# Evolutionary Inference Predicts Novel ACE2 Protein Interactions Relevant to COVID-19 Pathologies

**DOI:** 10.1101/2021.05.24.445517

**Authors:** Austin A. Varela, Sammy Cheng, John H. Werren

**Author notes:** contributed equally (Austin A. Varela, John H. Werren).

## Abstract

Angiotensin-converting enzyme 2 (ACE2) is the cell receptor that the coronavirus SARS-CoV-2 binds to and uses to enter and infect human cells. COVID-19, the pandemic disease caused by the coronavirus, involves diverse pathologies beyond those of a respiratory disease, including micro-thrombosis (micro-clotting), cytokine storms, and inflammatory responses affecting many organ systems. Longer-term chronic illness can persist for many months, often well after the pathogen is no longer detected. A better understanding of the proteins that ACE2 interacts with can reveal information relevant to these disease manifestations and possible avenues for treatment. We have undertaken an approach to predict candidate ACE2 interacting proteins which uses evolutionary inference to identify a set of mammalian proteins that “coevolve” with ACE2. The approach, called evolutionary rate correlation (ERC), detects proteins that show highly correlated evolutionary rates during mammalian evolution. Such proteins are candidates for biological interactions with the ACE2 receptor. The approach has uncovered a number of key ACE2 protein interactions of potential relevance to COVID-19 pathologies. Some proteins have previously been reported to be associated with severe COVID-19, but are not currently known to interact directly with ACE2, while additional predicted novel ACE2 interactors are of potential relevance to the disease. Using reciprocal rankings of protein ERCs, we have identified strongly interconnected ACE2 associated protein networks relevant to COVID-19 pathologies. ACE2 has clear connections to coagulation pathway proteins, such as Coagulation Factor V and fibrinogen components FGG, FGB, and FGA, the latter possibly mediated through ACE2 connections to Clusterin (which clears misfolded extracellular proteins) and GPR141 (whose functions are relatively unknown). ACE2 also connects to proteins involved in cytokine signaling and immune response (e.g. IFNAR2, XCR1, and TLR8), and to Androgen Receptor (AR). The ERC prescreening approach has also elucidated possible functions for previously uncharacterized proteins and possible new functions for well-characterized ones. Suggestions are made for the validation of ERC predicted ACE2 protein interactions. We propose that ACE2 has novel protein interactions that are disrupted during SARS-CoV-2 infection, contributing to the spectrum of COVID-19 pathologies.

## Introduction

The coronavirus SARS-CoV-2 is causing severe pathologies and death among infected individuals across the planet. In addition to the symptoms expected from a respiratory disease, the infection can develop systemic manifestations (Gupta et al., 2020; Terpos et al., 2020; Siddiqi, Libby & Ridker, 2021). As a consequence, a wide range of pathologies are associated with COVID-19, including vascular system disruption, the extensive formation of blood clots (thrombosis) resulting in microvascular injury and stroke (Magro et al., 2020; Connors & Levy, 2020), gastrointestinal complications (Luo, Zhang & Xu, 2020) cardiac and kidney pathologies, ocular and dermatological symptoms (Bouaziz et al., 2020), neurological manifestations (Niazkar et al., 2020; Taquet et al., 2021), male infertility (Khalili et al., 2020), and a Kawasaki-like blood and heart disorder in children (Jones et al., 2020; Morand, Urbina & Fabre, 2020). A severe and often lethal immunoreaction can occur from respiratory and other infection sites, termed a “cytokine storm” (Chen et al., 2020). Even after acute SARS-CoV-2 infection has passed, individuals can suffer a suite of complications for many months, termed “Long Haul” syndrome (López-León et al., 2021), and the causes of these syndromes are not well understood.

Angiotensin-converting enzyme 2 (ACE2) is of obvious interest because it is a primary receptor for SARS-CoV-2 entry into human cells (Lan et al., 2020). However, ACE2 also plays a role in other important processes, such as regulation of blood pressure and vasodilation by the renin-angiotensin system (RAS), and protein digestion in the gut (Kuba et al., 2010). SARS-CoV-2 binding to ACE2 leads to a downregulation in ACE2 function (Verdecchia et al., 2020) which may be linked to the systemic damage by COVID-19 (Medina-Enríquez et al., 2020). It has been proposed that ACE2 receptor degradation during SARS-CoV-2 infection disrupts ACE2 protection from inflammatory processes through the RAS and bradykinin pathways, possibly explaining patterns of COVID-19 severity with age and sex (Bastolla, 2020; Bastolla et al., 2021). As well as being a cell receptor, a circulating soluble form of the ectodomain of ACE2 (sACE2) is shed from cells and found in blood plasma, but the biological function of circulating ACE2 remains relatively unknown. Elevated levels of sACE2 have been detected in critically ill COVID-19 patients (van Lier et al., 2021) which coincides with a reduced expression of membrane-bound ACE2 (Medina-Enríquez et al., 2020), and a recent study indicates that sACE2 may assist SARS-CoV-2 entry into cells via other receptors (Yeung et al., 2021).

In general, ACE2’s protein-protein interaction network is likely to contribute to COVID-19 pathologies, due to ACE2’s role in systemic processes that are disrupted by the infection. Therefore, a fuller knowledge of ACE2 protein interactions is important to a better understanding of COVID-19 pathologies, including those that go beyond respiratory illness.

Common methods to identify protein-protein interactions include protein co-localization and precipitation, genetic manipulation, and proteomic profiling (Rao et al., 2014). Evolutionary approaches have also been used to evaluate protein interactions (De Juan, Pazos & Valencia, 2013), particularly to identify functional domains within proteins by sequence conservation. Another set of methods utilize evolutionary rate correlations (also called evolutionary rate covariance or evolutionary rate coevolution). The concept is that coevolving proteins will show correlated rates of change across evolution (Wolfe & Clark, 2015). The approach has been used to detect physical interactions within and among proteins, as well as shared functionality (Clark, Alani & Aquadro, 2012). For example, it has been employed to identify gene networks for post-mating response (Findlay et al., 2014), ubiquitination (Böhm et al., 2016), and recombination (Godin et al., 2015), and more recently to identify DNA repair genes (Brunette et al., 2019), cadherin-associated proteins (Raza et al., 2019), and a mitochondrial associated zinc transporter (Kowalczyk et al., 2021), with subsequent experimental support. Evolutionary rate correlation (ERC) approaches are relatively inexpensive screening tools for detecting candidate protein interactions, and can also detect novel protein interactions that are not readily found in more traditional proteomic and genetic approaches (Colgren & Nichols, 2019; Yan, Ye & Werren, 2019). As such, “the ERC method should be a part of the toolkit of any experimental cell or developmental biologist” (Colgren & Nichols, 2019).

We have developed an evolutionary rate correlation (**ERC**) method that uses well-established phylogenies based on multiple lines of evidence (e.g. Misof et al. 2014 for insects and Kumar et al. 2017 for mammals) and calculates protein evolutionary rates for terminal branches for different proteins across a set of related species (Fig. 1). The approach is predicated on the idea that proteins that have strong evolutionary rate correlations are more likely to have functional interactions that are maintained by their coevolution, a conclusion supported by its predictive power in identifying known nuclear-mitochondrial encoded protein interactions in insects (Yan, Ye & Werren, 2019). The study also found that nuclear-encoded proteins and amino acids in contact with their mitochondrial-encoded components (e.g. oxidative phosphorylation proteins or mitochondrial ribosomal RNA) have significantly stronger ERCs than those not directly in contact. This result implicates physical interactions between proteins as one driver of evolutionary rate correlations, at least among nuclear-mitochondrial interactions.

**Figure 1.**
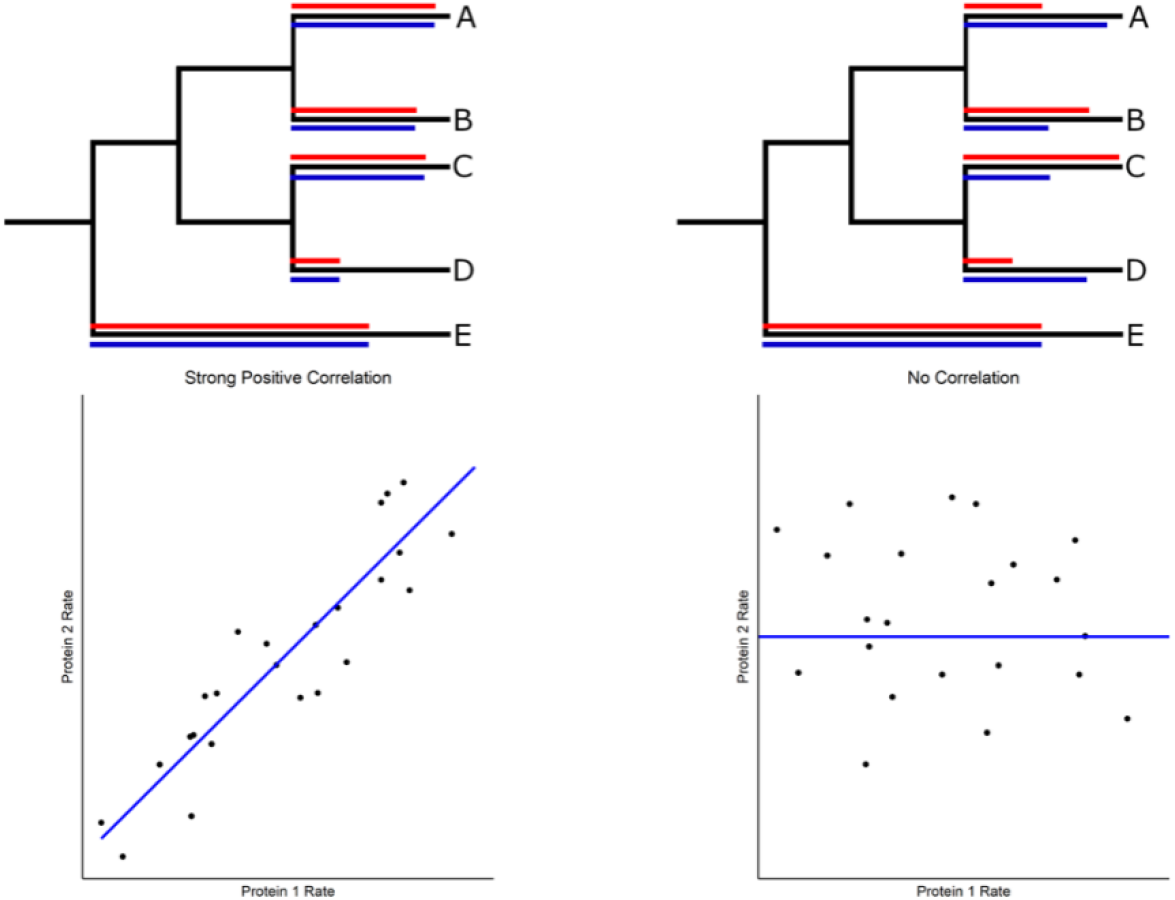
Evolutionary rate correlations. The Spearman rank correlations between two proteins are calculated based on rates of protein evolution on terminal branches of a phylogeny. The relative rates of two proteins (red and blue lines) are shown in the hypothetical phylogenetic trees. Correlated and uncorrelated protein rates are illustrated below using a larger number of terminal branches (data points) than presented in the phylogeny.

We have developed a reciprocal rank approach to identify ACE2 associated networks and propose that these strongly coevolving proteins reveal ACE2 protein interactions that could be disrupted by COVID-19, thus contributing to its diverse pathologies. Particularly noteworthy are strong connections to coagulation pathway proteins, cytokine signaling, inflammation, immunity, and viral disease response. We recognize the importance of validation of these predicted ACE2 protein interactions and therefore propose steps for investigating possible applications of these findings to COVID-19 disease and treatment.

## Results

### A. Basic Approach

The basic methods are outlined here to provide context for the results which follow (details are provided in Methods). To identify candidate protein interactions using evolutionary rate correlation, we utilized the consensus TimeTree phylogenetic reconstruction for mammalian species (Kumar et al., 2017). A total of 1,953 proteins (including ACE2) were aligned and evolutionary rates for each protein were then calculated for terminal branches of the tree (Fig. 1). This was determined by dividing the protein-specific branch length on each terminal branch by terminal branch time from the consensus tree (Yan, Ye & Werren, 2019). Maximum likelihood branch lengths were estimated in IQ-TREE (Minh et al., 2020) using an empirical amino acid substitution matrix (see methods for details). To investigate evolutionary rate correlations (ERCs) among proteins, Spearman rank correlations were calculated for every protein pair using terminal branch rates (Fig. 1). Due to the large number of comparisons, a Benjamini-Hochberg false discovery rate (FDR) correction was calculated for each protein’s ERC set (significance threshold α = 0.05). We subsequently found that many proteins show a positive correlation between terminal branch time and evolutionary rate, and observed that short branches in relatively oversampled taxa significantly contributed to this correlation (Supplementary Text). We, therefore, removed species that accounted for short branches, which eliminated the protein evolutionary rate to branch time correlation (see Methods and Supplementary Text for details). ERCs were then recalculated, and our ERC analyses are based on this set of 60 taxa.

In addition, we tested whether the observed lower rates of protein evolution for shorter branches in the phylogeny are due to rates actually increasing over evolutionary time. This was accomplished by quantifying protein rates while sequentially extending branch lengths in different clades. The analysis shows that evolutionary rates for many proteins increase as branch length is increased (Supplementary Text, Supplementary Fig. S4). A likely explanation for the pattern is that protein coevolution is mostly episodic, and short branches in a phylogeny are less likely to capture such events. In additional analyses, we tested for but did not find significant confounding effects of taxon on the ERC results (Supplementary Text).

Our analyses are focused on candidate protein interactions involving ACE2 using evidence of highly significant ERCs. For this purpose, we first examine proteins in ACE2’s highest 2% of ERCs (top 40 proteins), all of which are highly significant after FDR correction (Table 1). Some of these ACE2 ERC proteins have been previously implicated in severe COVID-19 or SARS-CoV-2 gene expression effects on infected cells. However, while they have not been previously identified as having direct protein interactions with ACE2, this is predicted by our ERC analysis.

**Table 1.**
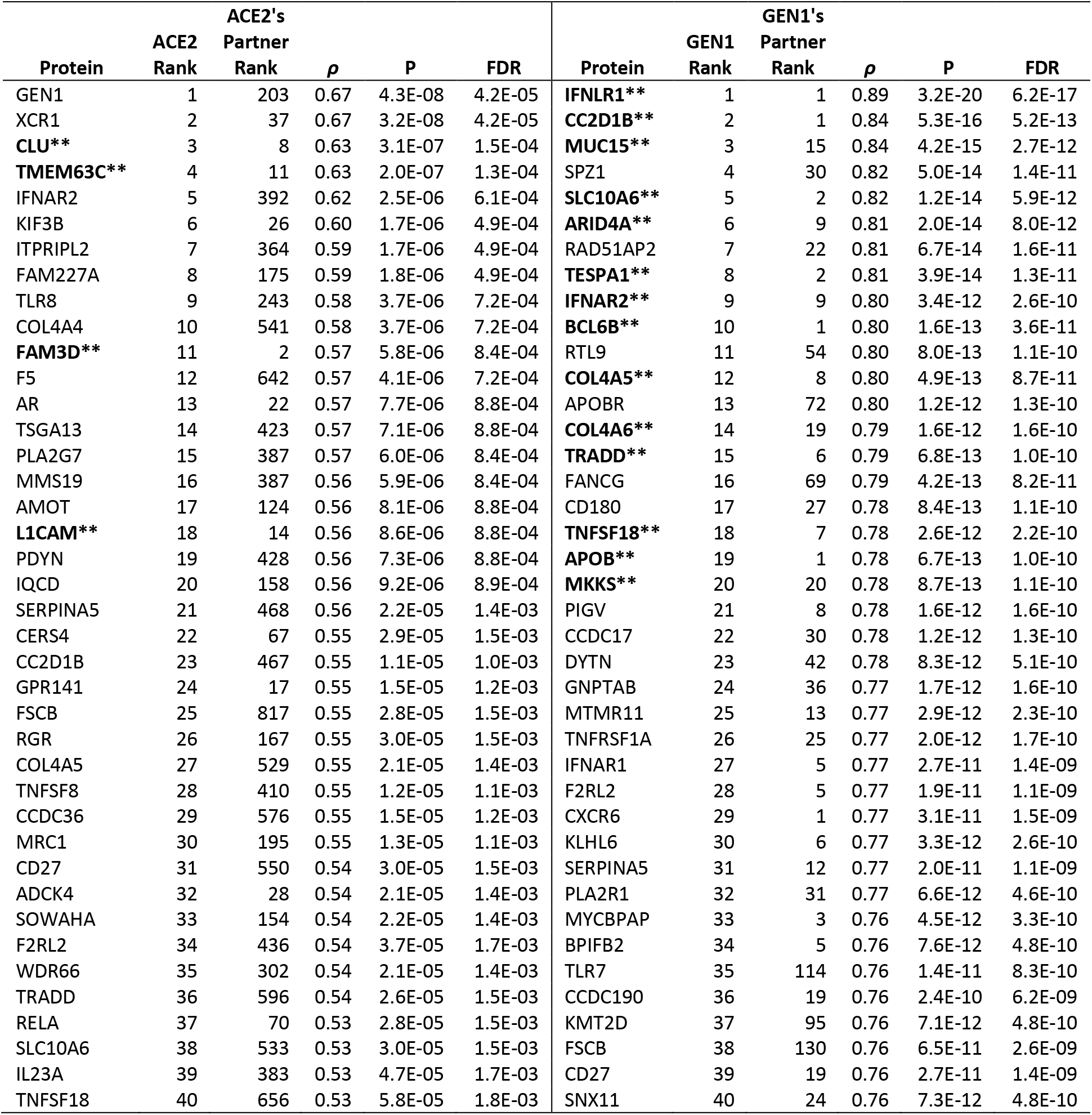
The top two percent (2%) of ERCs are shown for ACE2 and GEN1, ranked by descending ρ value. The table illustrates how reciprocal ranks can differ between proteins with significant evolutionary correlations, depending on how interconnected proteins are. GEN1 has many partners which rank GEN1 highly in their respective ERCs. Also indicated in the table are examples of reciprocal rank correlations in which both partners rank the other in their top 20 (indicated by bold and asterisks). These are used to construct reciprocal rank protein interaction networks.

X-C Motif Chemokine Receptor 1 (XCR1) provides an illustrative example. XCR1 is a cytokine signaling receptor and ACE2’s 2^nd^ highest ranked ERC, with a highly significant evolutionary rate correlation suggestive of a protein-protein interaction. XCR1 is in a small genomic region that is implicated in severe COVID-19 by additional genome-wide association studies (Severe Covid-19 GWAS Group, 2020; Fricke-Galindo & Falfán-Valencia, 2021). Another example is Interferon alpha/beta receptor 2 (IFNAR2) which, in a genome-wide association study (GWAS) and multi-omic analysis by Pairo-Castineira et al. (2021), was implicated in severe COVID-19. We therefore added it to our analysis, and surprisingly found it to be highly ranked (5^th^) among ACE2 ERCs. Clusterin (CLU) is the 3^rd^ strongest ERC of ACE2 and the ACE2-CLU pair show high reciprocal ranks to each other (3^rd^ in ACE2’s set, 8^th^ in CLU’s set). CLU prevents the aggregation of misfolded proteins in the blood and delivers them to cells for degradation in lysosomes (Sánchez-Martín & Komatsu, 2020). CLU connects to key proteins in the coagulation pathway based on its reciprocal rank network (Section C, Fig. 2). CLU has been implicated in coronavirus infections, as one of only two proteins showing significant expression changes in cells infected by three different coronaviruses tested, including SARS-CoV-2 (Singh et al., 2021). The examples above lend credence to the proposition that the ERC approach is detecting ACE2 protein interactions that have implications to COVID-19.

**Figure 2.**
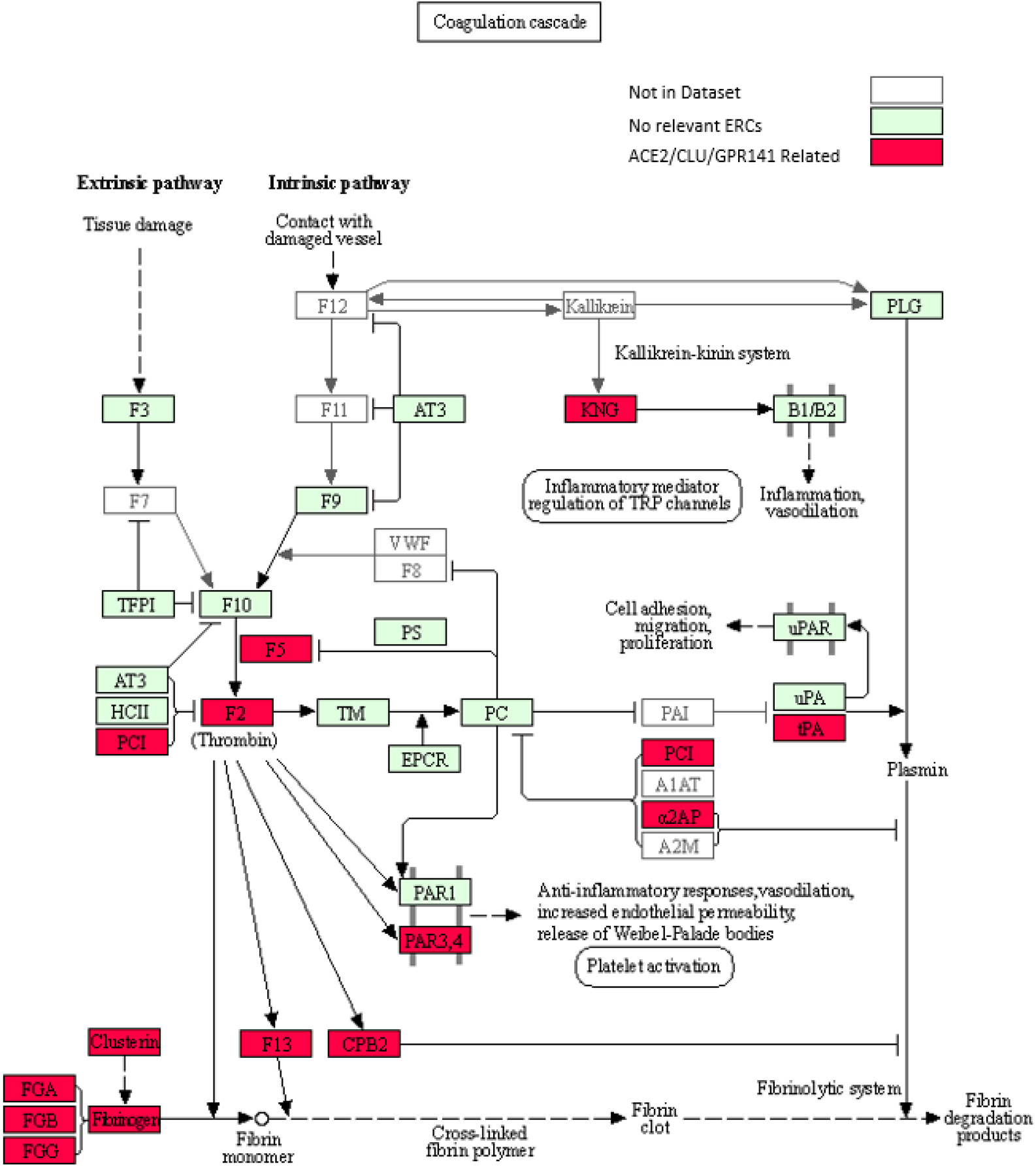
KEGG Coagulation cascade pathway (Kanehisa & Goto, 2000), with ACE2-CLU-GPR141 associated proteins (based on presence on any of their top 2% ERCs or in the ACE2 CRR network) indicated in red. The KEGG pathway has been supplemented to indicate the three fibrinogen proteins and clusterin associations previously discussed. Note the alternate protein names: PAR3,4 = F2RL2 & F2RL3 = Thrombin receptors; α2AP = Alpha-2-antiplasmin = SERPINF2; PLAT = tPA, and PCI = SERPINA5 = Protein C Inhibitor.

Differences in ERC rank between protein pairs for the same correlation can occur because some proteins have higher and more extensive ERC connections than do. As a result, while two proteins can have a significant ERC with each other, each one’s rank may differ in their respective ERC lists, as illustrated for ACE2 and GEN1 (Table 1). GEN1 (Flap endonuclease GEN homolog 1) is ACE2’s top-ranked ERC, and is a DNA nuclease whose primary functions are resolution of DNA Holliday junctions and DNA damage checkpoint signaling (Chan & West, 2015). This protein shows high ERCs and is ranked highly in the ERC sets for many other proteins, suggesting central connectivity. As described further in Section C2, GEN1 shows unexpected enrichments for immune functions, perhaps related to its role in DNA damage checkpoint signaling.

Because our focus is on identifying strong candidate interactions involving ACE2 and its predicted partners, we utilize the rank information to identify proteins with high reciprocal ranks. Specifically, we focus on the strongest reciprocal ranks (RR) defined by ranks of less than or equal to 20 (RR20), which is the highest one percent of each protein’s ERCs, and use these to develop reciprocal rank networks (Section C). Although speculative, we posit that protein pairs with high reciprocal ranks are likely to be strongly coevolving (i.e. both partners evolving reciprocally due to selective pressures acting on interacting domains between them). In contrast, protein pairs with a significant evolutionary rate correlation only one ranks highly (e.g. within the top two percent) in the ERC set of the other, are more likely to be due to “unidirectional” evolution. The rationale is that proteins with many significant ERC partners are under selective pressures primarily from their top evolutionary partners, whereas other interactors evolve primarily in response to the forces shaped by their stronger partner(s).

The view that ERCs are detecting protein interactions relevant to COVID-19 is further supported by the analysis of ACE2 reciprocal rank ERC networks (Section C below). Noteworthy in this regard are additional proteins in the coagulation pathway, such as Coagulation Factor V (F5), Fibrinogen Alpha Chain (FGA), Fibrinogen Beta Chain (FGB), and Fibrinogen Gamma Chain (FGG). Thrombosis (blood clotting) is a major pathology of COVID-19 (Gupta et al., 2020). Connections of ACE2 with the proteins above could relate to severe blood clotting problems in COVID-19 infections. ACE2 networks also show strong enrichments of cytokine signaling, viral (and pathogen) infections, and inflammatory response terms (Supplementary File S3), which are clearly relevant to COVID-19 pathologies such as cytokine storms and systemic inflammation.

In yet other cases, we have found proteins with significant ACE2 ERCs or ACE2 network connections, but for which there is little functional information. We can use their ERCs to suggest possible functions for future investigation. Examples of this are GPR141, C16orf78, and CCDC105. Finally, ERCs for proteins of known function (such as Coagulation Factor F5 and GEN1) indicate likely additional roles, suggesting these proteins have unrecognized “moonlighting” functions (Jeffery, 1999).

Below, we first describe proteins of interest to which ACE2 has significant ERCs, summarize aspects of their known biological functions, and examine significantly enriched functional categories for these ERCs. We then build and evaluate two different networks for ACE2 interacting proteins (Section C), one of which reveals connections to coagulation pathways and the other to cytokine-mediated signaling, viral response, and immunity. Finally, we discuss the potential implications of these predicted ACE2 interactions to COVID-19 pathologies and propose some specific hypotheses that emerge from this analysis.

### B. Top ERC Interactions Link ACE2 to COVID Pathologies

To investigate protein associations of ACE2, we first determined the protein enrichment categories for its top 2% ERC proteins (based on Spearman rank correlation coefficients, **ρ**) using the gene set enrichment package Enrichr (Xie et al., 2021) (Table 2). The top two KEGG_2019_Human enrichments are for complement and coagulation cascade related (FDR = 2.0E-03) and cytokine-cytokine receptor interaction related (FDR = 2.0E-03) terms. This finding is consistent with two hallmarks of COVID-19 pathology, abnormal systemic blood-clotting (thrombosis) and cytokine storms (Coperchini et al., 2020; Fei et al., 2020). Additionally, several terms related to viral/bacterial-specific infection are significantly enriched, such as Tuberculosis (FDR = 1.4E-02), HPV infection (FDR = 1.4E-02), measles (FDR = 2.4E-02) and Hepatitis C (FDR = 3.1E-02). Gene Ontology Biological Process also shows enrichment for tumor necrosis factor (TNF) pathways, including the signaling pathway (FDR = 3.9E-03) and cellular responses (FDR = 1.6E-02). Additional terms are shown in Table 2.

**Table 2.** Enrichment categories for ACE2’s top 2% proteins by ERC. Key enrichments include complement and coagulation cascades, cytokine-cytokine signaling, and different pathogen infections.

| Enrichr Gene set | Term | FDR<br>P-value | Odds<br>Ratio | Gene List |
| --- | --- | --- | --- | --- |
| KEGG_2019_Human | Complement and coagulation cascades | 2.03E-03 | 25.9 | CLU, F2RL2, F5, SERPINA5 |
| KEGG_2019_Human | Cytokine-cytokine receptor interaction | 2.03E-03 | 10.5 | TNFSF18, IFNAR2, XCR1, IL23A, CD27, TNFSF8 |
| KEGG_2019_Human | Tuberculosis | 1.38E-02 | 11.0 | IL23A, TRADD, MRC1, RELA |
| KEGG_2019_Human | Human papillomavirus infection | 1.38E-02 | 7.6 | IFNAR2, COL4A4, TRADD, COL4A5, RELA |
| KEGG_2019_Human | Protein digestion and absorption | 1.38E-02 | 16.3 | ACE2, COL4A4, COL4A5 |
| KEGG_2019_Human | Pathways in cancer | 1.38E-02 | 5.7 | IFNAR2, AR, IL23A, COL4A4, COL4A5, RELA |
| KEGG_2019_Human | Small cell lung cancer | 1.38E-02 | 15.8 | COL4A4, COL4A5, RELA |
| KEGG_2019_Human | Amoebiasis | 1.38E-02 | 15.3 | COL4A4, COL4A5, RELA |
| KEGG_2019_Human | AGE-RAGE signaling pathway in diabetic complications | 1.38E-02 | 14.6 | COL4A4, COL4A5, RELA |
| KEGG_2019_Human | Toll-like receptor signaling pathway | 1.39E-02 | 14.0 | IFNAR2, TLR8, RELA |
| KEGG_2019_Human | Sphingolipid signaling pathway | 1.85E-02 | 12.2 | CERS4, TRADD, RELA |
| KEGG_2019_Human | Relaxin signaling pathway | 2.18E-02 | 11.2 | COL4A4, COL4A5, RELA |
| KEGG_2019_Human | Measles | 2.38E-02 | 10.5 | IFNAR2, TRADD, RELA |
| KEGG_2019_Human | Hepatitis C | 3.05E-02 | 9.3 | IFNAR2, TRADD, RELA |
| KEGG_2019_Human | Cocaine addiction | 3.05E-02 | 19.7 | PDYN, RELA |
| KEGG_2019_Human | PI3K-Akt signaling pathway | 4.17E-02 | 5.5 | IFNAR2, COL4A4, COL4A5, RELA |
| KEGG_2019_Human | Kaposi sarcoma-associated herpesvirus infection | 4.17E-02 | 7.7 | IFNAR2, TRADD, RELA |
| KEGG_2019_Human | Inflammatory bowel disease (IBD) | 4.34E-02 | 14.7 | IL23A, RELA |
| KEGG_2019_Human | Epstein-Barr virus infection | 4.34E-02 | 7.1 | IFNAR2, TRADD, RELA |
| KEGG_2019_Human | Adipocytokine signaling pathway | 4.34E-02 | 13.8 | TRADD, RELA |
| KEGG_2019_Human | RIG-I-like receptor signaling pathway | 4.34E-02 | 13.6 | TRADD, RELA |
| KEGG_2019_Human | Pertussis | 4.85E-02 | 12.5 | IL23A, RELA |
| GO_Biological_Process_2018 | tumor necrosis factor-mediated signaling pathway (GO:0033209) | 3.86E-03 | 21.0 | TNFSF18, TRADD, CD27, TNFSF8, RELA |
| GO_Biological_Process_2018 | cellular response to tumor necrosis factor (GO:0071356) | 1.63E-02 | 13.1 | TNFSF18, TRADD, CD27, TNFSF8, RELA |
| GO_Biological_Process_2018 | immunoglobulin mediated immune response (GO:0016064) | 1.63E-02 | 154.6 | CD27, TLR8 |
| GO_Biological_Process_2018 | B cell mediated immunity (GO:0019724) | 1.63E-02 | 154.6 | CD27, TLR8 |
| GO_Biological_Process_2018 | positive regulation of NF-kappaB transcription factor activity (GO:0051092) | 1.85E-02 | 15.6 | TNFSF18, TRADD, CLU, RELA |
| GO_Biological_Process_2018 | I-kappaB kinase/NF-kappaB signaling (GO:0007249) | 2.26E-02 | 26.3 | TRADD, TLR8, RELA |
| GO_Biological_Process_2018 | cytokine-mediated signaling pathway (GO:0019221) | 3.11E-02 | 5.7 | TNFSF18, IFNAR2, IL23A, TRADD, CD27, TNFSF8, RELA |
| GO_Biological_Process_2018 | regulation of inflammatory response (GO:0050727) | 3.11E-02 | 11.9 | ACE2, IL23A, PLA2G7, RELA |
| GO_Biological_Process_2018 | positive regulation of defense response (GO:0031349) | 3.26E-02 | 20.0 | IL23A, TLR8, PLA2G7 |
| WikiPathways_2019_Human | Complement and Coagulation Cascades WP558 | 2.38E-02 | 25.8 | CLU, F5, SERPINA5 |
| WikiPathways_2019_Human | EBV LMP1 signaling WP262 | 4.24E-02 | 44.2 | TRADD, RELA |
| WikiPathways_2019_Human | Toll-like Receptor Signaling Pathway WP75 | 4.24E-02 | 14.2 | IFNAR2, TLR8, RELA |
| WikiPathways_2019_Human | Toll-like Receptor Signaling WP3858 | 4.36E-02 | 32.0 | TLR8, RELA |
| WikiPathways_2019_Human | miRNAs involvement in the immune response in sepsis WP4329 | 4.95E-02 | 26.5 | TLR8, RELA |
| WikiPathways_2019_Human | Regulation of toll-like receptor signaling pathway WP1449 | 4.96E-02 | 10.4 | IFNAR2, TLR8, RELA |

The ACE2 ERC analysis indicates that ACE2 is “coevolving” with proteins involved in the complement and coagulation pathways, cytokine signaling, TNF, and pathogen response pathways. Here, we summarize results and background information on some of the key proteins among ACE2’s ERCs (more extended summaries of each protein are in the Supplementary Text).

Among ACE2’s strongest ERCs are proteins involved in immunity. For example, XCR1 (X-C Motif Chemokine Receptor 1) is ACE2’s 2^nd^ top-ranked ERC (ρ = 0.67, FDR = 6.18E-05). It is a chemokine XCL1 receptor involved in immune response to infection and inflammation (Lei & Takahama, 2012). Strikingly, the Severe Covid-19 GWAS Group (2020) detected a small genomic region containing six genes that significantly associate with severe COVID-19, one of which is XCR1. Our finding that XCR1 is ACE2’s 2^nd^ highest ERC interactor lends independent support for a relationship between COVID-19 and XCR1. Furthermore, it suggests that a direct interaction between ACE2 and XCR1 could be involved in COVID-19 pathologies. To our knowledge, there are no other reports of interactions between these two proteins.

Another striking connection of ACE2 ERC to immunity is through IFNAR2 (Interferon alpha/beta receptor 2), which has a highly significant ACE2 ERC correlation (ρ = 0.62, FDR = 6.1E-04). IFNAR2 forms part of an important receptor complex with IFNAR1 (Thomas et al., 2011) involved in interferon signaling through the JAK/STAT pathway to modulate immune responses. IFNAR2 has been implicated in severe COVID-19, based on mendelian randomization, genome-wide associations, and gene expression changes (Zhang et al., 2020; Liu et al., 2021; Pairo-Castineira et al., 2021). Our data provide independent support for a role, possibly mediated through ACE2 interactions. Interferon pathways are important in antiviral defense, but also can contribute to cytokine storms and COVID-19 pathologies (McKechnie & Blish, 2020). Other immune-related proteins with high ERC connections to ACE2 include TLR8 (Toll-like Receptor 8), FAM3D (FAM3 metabolism regulating signaling molecule D), and PLA2G7 (phospholipase A2 group VII).

Coagulation pathway proteins figure prominently in ACE2 ERC-predicted protein interactions (Table 3, Fig. 2). This is reflected both in significant enrichment for coagulation cascade proteins in the top 2% strongest ACE2 ERCs (Table 2) and the strong reciprocal rank network for ACE2 (Section C, Fig. 3). The finding has obvious potential implications to a hallmark pathology of COVID-19, systemic coagulopathy (Wright et al., 2020; Medcalf, Keragala & Myles, 2020). A list of coagulation and blood-related proteins associated with ACE2 is presented in Table 3. Among ACE2’s top 2% ERCs associated with coagulation pathway are Coagulation Factor V (F5), Protein C inhibitor (SERPINA5 aka PCI), and Thrombin Receptor 2 (F2RL2) (Table 1).

**Table 3.** Coagulation and blood-related proteins in the ACE2 CRR and URR Networks as well as the top 1% ACE2 ERC list.

| Name | Full Name | Brief Description |
| --- | --- | --- |
| ACE2 | Angiotensin Converting Enzyme 2 | Catalyzes the cleavage of angiotensin I to angiotensin 1-9 and angiotensin II to angiotensin 1-7 (Burrell et al., 2004) |
| FGA | Fibrinogen alpha chain | Bind to FGB and FGG to form fibrinogen, used to form blood clots (Mosesson, 2005) |
| FGB | Fibrinogen beta chain | Bind to FGA and FGG to form fibrinogen, used to form blood clots (Mosesson, 2005) |
| FGG | Fibrinogen gamma chain | Bind to FGA and FGB to form fibrinogen, used to form blood clots (Mosesson, 2005) |
| CPB2 | Carboxypeptidase B2 | Inhibits fibrinolysis (Leenaerts et al., 2018) |
| SERPINF2 | Serpin family F member 2 (alpha-2-antiplasmin) | Inhibits Plasmin, a protein involved in fibrinolysis (Kanehisa & Goto, 2000) |
| CD34 | CD34 molecule | Associated with hematopoiesis and stem cells (Fina et al., 1990) |
| CLU | Clusterin | Binds to Fibrinogen (Wyatt & Wilson, 2010) |
| MAS1 | MAS1 Proto-Oncogene, G Protein-Coupled Receptor | Receptor for angiotensin-(1-7) (Burrell et al., 2004) |
| FAM3D | FAM3 Metabolism Regulating Signaling Molecule D | Implicated in inflammatory responses in the gastrointestinal tract and is a chemoattractant for neutrophils and monocytes (Peng et al., 2016) |
| GPR141 | G Protein-Coupled Receptor 141 | High expression in blood, granulocytes, Kupfer cells, and macrophages (Stelzer et al., 2016) |
| TMEM63C | Transmembrane Protein 63C | Interacts with angiotensin II (Eisenreich et al., 2020) |
| LECT2 | Leukocyte Cell-derived Chemotaxin 2 | Involved in macrophage activation, insulin resistance and diabetes, and neutrophil chemotaxis (Yamagoe et al., 1996; Zhang et al., 2018; Takata et al., 2021) |
| ETS1 | ETS proto-oncogene 1, transcription factor | Transcription factor involved in cytokine/chemokine processes and angiogenesis (Stelzer et al., 2016) |
| ZBTB43 | Zinc Finger and BTB Domain containing 43 | Associated with Diamond-Blackfan Anemia 4, in which the bone marrow is unable to make enough red blood cells to carry oxygen (Stelzer et al., 2016) |
| COL4A4 | Collagen Type IV Alpha 4 | Subunit of Collagen Type 4, which are a part of the basement membrane which resides between epithelial cells (Stelzer et al., 2016) |
| F13B | Coagulation Factor XIII B chain | Stabilizes F13A subunits, while it does not have enzymatic abilities it is thought to be a plasma carrier molecule (Stelzer et al., 2016) |
| AMOT | Angiomotin | Associated with angiogenesis and endothelial cell movement (Bratt et al., 2005; Aase et al., 2007) |
| PDYN | Prodynorphin | Inhibits vasopressin secretion (Yamada et al., 1988) |

**Figure 3.**
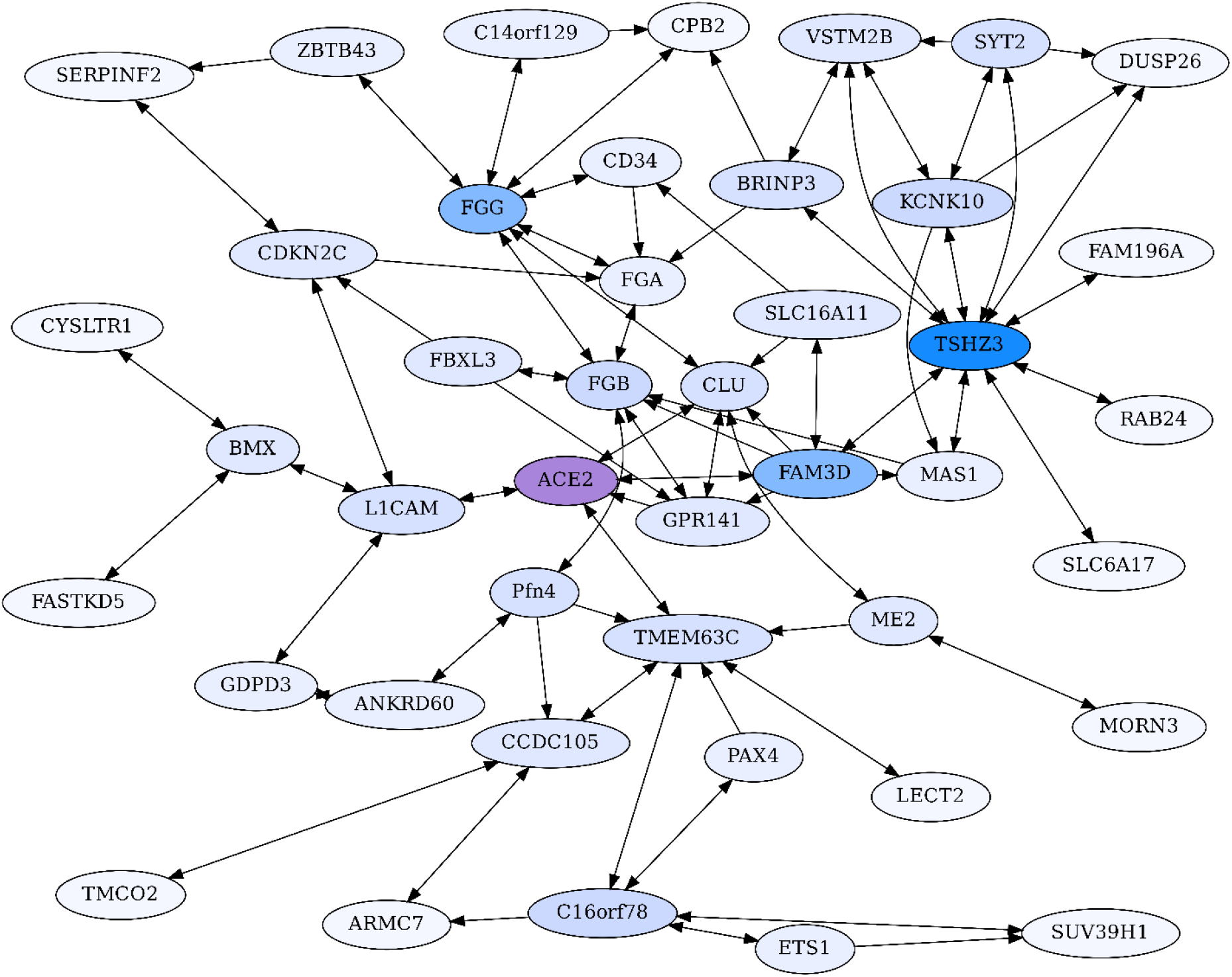
ACE2 Centric Reciprocal Rank Network (CRR). Proteins with ERC reciprocal ranks ≤ 20 are shown by double-headed arrows, and unidirectional ranks ≤ 20 connecting to the RR backbone are indicated by single-headed arrows. ACE2 has extensive connections to coagulation proteins mediated primarily through Clusterin (CLU) and GPR141. ACE2 is highlighted in purple, and blue shading intensity indicates the level of reciprocal connectivity for different proteins.

Also relevant to coagulopathy are Clusterin (CLU) and the orphan G protein-coupled receptor 141 (GPR141). The chaperone protein CLU has a soluble form that circulates in the blood and is part of the “cleaning squad” that clears misfolded extracellular proteins for delivery to lysosomes and degradation (Itakura et al., 2020; Sánchez-Martín & Komatsu, 2020). It is the 3^rd^ highest ACE2 ERC (ρ = 0.63, FDR = 1.5E-04), and these two proteins show strong reciprocal ranks (3, 8), likely supporting biological interactions. Relevant to this point is that both ACE2 and CLU have soluble forms that circulate in the blood (Itakura et al., 2020). Of direct relevance to COVID-19 and possible ACE2-CLU protein interactions, Singh et al. (2021) found in cells infected with different coronaviruses (SARS-CoV-2, SARS-CoV, and MERS-CoV), only two genes were found to be differentially expressed in all three, with CLU being one.

CLU’s top 2% strongest ERCs show highly significant enrichment for terms relating to coagulation cascades and clot formation (Supplementary File S3, e.g. “Complement and coagulation cascades”, FDR = 6.28E-12), as well as significant terms that are relevant to immunity, such as “Immune system” (FDR = 4.8E-03) and “activated immune cell type” (FDR = 3.4E-05). Among its top ERC proteins relevant to coagulation process are Coagulation Factor V (F5, ρ = 0.67, FDR = 9.1E-06, rank 3), Fibrinogen Gamma chain (FGG, ρ = 0.59, FDR = 1.7E-04, rank 18), Coagulation Factor XIII B chain (F13B, ρ = 0.63, FDR = 2.8E-05, rank 19), and Fibrinogen Alpha chain (FGA, ρ = 0.57, FDR = 2.9E-04, rank 27) (Fig. 3, Supplementary File S1). Notably, fibrinogen is a major binding “client” of Clusterin in stressed plasma (Wyatt & Wilson, 2010). Little is known about GPR141; however, the ERC analysis suggests an important role in blood coagulation. Among GPR141’s top ERC proteins relevant to coagulation process are Kininogen 1 (KNG1, ρ = 0.60, FDR = 9.3E-04, rank 5), Plasminogen Activator (PLAT, ρ = 0.58, FDR = 6.5E-04, rank 6), Thrombin (Coagulation Factor II or F2, ρ = 0.58, FDR = 6.5E-04, rank 7), Fibrinogen Beta chain (FGB, ρ = 0.57, FDR = 6.5E-04, RR 11, 11), Complement C1s (C1S, ρ = 0.54, FDR = 1.6E-03, rank 22), F2R-like thrombin (also called trypsin receptor 3; F2RL3, ρ = 0.52, FDR = 2.6E-03, rank 37), and Coagulation Factor V (F5, ρ = 0.52, FDR = 1.7E-03, rank 39) (Fig. 2, Supplementary File S1).

GPR141 has a highly significant ERC to CLU, with these two proteins being each other’s first ranking ERCs (ρ = 0.68, FDR = 9.06E-06, RR 1,1). The pattern suggests a strong biological interaction, although none is described in the literature. The result supports investigating functional interactions between CLU and GPR141, based upon their high ERC and reciprocal ranks. Our network analysis (Section C) further supports extensive interconnections among ACE2, Clusterin, GPR141, and coagulation pathway proteins, implicating the protein interaction pathway as a possibly significant contributor to disruption of coagulation in COVID-19 disease. Coagulation cascade proteins found in the ACE2’s top 2% ERCs, ACE2 reciprocal rank network, and Clusterin-GPR141 associated proteins are highlighted in Figure 2.

Androgen Receptor (AR, ρ = 0.57, FDR = 8.8E-04, rank 13), the receptor for the male hormone androgen, plays a major role in reproductive system development, somatic differentiation, and behavior (Matsumoto et al., 2008). Androgen-AR signaling induces ACE2 (Wu et al., 2020), while knockdowns of AR result in downregulation of ACE2 (Samuel et al., 2020). AR agonists also reduce SARS-CoV-2 spike protein-mediated cellular entry (Deng et al., 2021). Additionally, AR is associated with COVID comorbidities (Dolan et al., 2020), and recently implicated in the severity of COVID-19 in women with polycystic ovarian syndrome, a disorder associate with high androgen levels and androgen sensitivity (Gotluru et al., 2021). Our ERC finding indicating ACE2 and AR coevolution suggests a regulatory feedback between these two proteins, which could be relevant to COVID-19 severity and other sex differential pathologies, such as cardiovascular disease (Viveiros et al., 2021).

Other notable significant ACE2 ERCs (Table 1) include Metabolism regulating signaling molecule D (FAM3D), Transmembrane-protein 63C (TMEM63C); Collagen Type IV Alpha 4 (COL4A4), L1 cell adhesion molecule (L1CAM), and ITPRIP-like 2 (ITPRIPL2). More detailed information on these and other proteins mentioned in this section is provided in Section C and the Supplementary Text.

### C. ERC Reciprocal Rank Networks implicate coagulation pathways and immunity

As mentioned previously, two proteins with a significant evolutionary rate correlation may often “rank” each other differently in their respective top ERC connections. This occurs because some proteins have more extensive ERC connections than others. High reciprocal ERC ranks between protein pairs may be more indicative that they are under strong coevolutionary pressure in their sequence and function. We have thus found it useful to evaluate these reciprocal rank connections as a network. The rationale is that such proteins are likely to be reciprocally evolving (“coevolving”). To build reciprocal rank networks, we use protein pairs that reciprocally share ranks less than or equal to 20 (RR20).

A core ACE2 reciprocal rank network was generated by building reciprocal rank connections (RR20) outward of ACE2, to provide a backbone set of RR20 protein connections. The backbone was expanded on by adding the RR20 connections of the non-ACE2 backbone proteins. Unidirectional ERCs (≤ rank 20) were then added between proteins within the RR set to produce an ACE2 Core Reciprocal Rank (**CRR**) Network (Fig. 3). The network is designed to capture features of ACE2’s protein interactions as revealed by the strong reciprocal evolutionary correlations among proteins.

ACE2 also has highly significant ERCs to proteins that do not rank ACE2 within their top 1% of ERCs, due to those proteins having more protein interactions with higher ERCs. A second network was therefore generated using ACE2’s top ten unidirectional ERCs, followed by calculating the RR20 associations for those proteins. This second network is referred to as the ACE2 Unidirectional Reciprocal Rank Network (**URR**).

These are presented below. In general, the reciprocal ranks analysis lends credence to our proposition that ERCs reveal real biological interactions, as well as providing predictions for novel protein interactions possibly of importance to COVID-19 pathologies and protein-interaction networks.

### C1 The ACE2 Core Reciprocal Rank (CRR) Network

The CRR network (Fig. 3) is designed to capture essential features of ACE2’s protein interactions as revealed by the strong reciprocal correlations among proteins.

The most striking aspects of the ACE2 CRR Network are extensive connections to the coagulation pathway and blood-associated proteins (Fig. 3, Table 3). This is, of course, very relevant to COVID-19 due to extensive clotting pathologies and stroke associated with COVID-19 (Bonaventura et al., 2021), as well as microvascular clotting and the apparent shut-down of fibrinolysis (Wright et al., 2020). Extensive blood coagulation of COVID-19 patients can even lead to clogging of dialysis equipment (Rabb, 2020). This hallmark pathology of COVID-19 indicates a disruption in coagulation and fibrinolysis pathways, and our findings of extensive network connections between ACE2 and coagulation-fibrinolysis pathway proteins could be relevant.

ACE2 connects to coagulation pathway proteins through F5, CLU, FAM3D, and GPR141 (Fig. 2, Fig. 3). CLU-GPR141 form a high RR ERC (ranks 1,1), strongly suggesting coevolution of these proteins and physical/functional interactions. Both CLU and GPR141 then connect to the fibrinogen proteins FGB and FGGB. FGA, FGB, and FGG are the three protein components that make up fibrinogen, which during the clotting process are converted into fibrin monomers, which subsequently cross-link to form the fibrin clot (Mosesson, 2005). All three proteins form a RR20 triad, indicating protein coevolution. FGG is a hub for RR ERCs to several other proteins (e.g. CD34, CPB2, C14or129, and ZBTB43). ZBTB43 is noteworthy, as it is associated with the blood diseases Diamond-Blackfan Anemia 4 and Hemochromatosis Type 2 (Stelzer et al., 2016). The former disrupts red blood cell formation in the bone marrow and the latter causes iron accumulation in the body. In terms of tissue distribution, ZBTB43 is enhanced in bone marrow (Uhlén et al., 2015). Cellularly, it is found mainly in nucleoplasm and nucleoli, suggesting regulatory functions, as might be expected for a transcription factor-like zinc finger domain protein. Most noteworthy, Mamoor (2020) has shown that ZBTB43 is differentially expressed in human microvascular endothelial cells and human cell cultures infected with coronaviruses (e.g MERS-CoV and human coronavirus 229E). So, this is yet another member of the ACE2 protein Network which is implicated in coronavirus infection. In turn, ZBTB43 has a RR connection with SERPINF2, which enhances clotting by inhibiting plasmin, an enzyme that degrades fibrin, the main component of clots. Mutations in SERPINF2 can cause severe bleeding disorders and upregulation of SERPINF2 is implicated in COVID-19 patient thrombosis (Jain et al., 2021; Lazzaroni et al., 2021). In turn, CPB2 (Carboxypeptidase B2) is a thrombin‐activated inhibitor of fibrinolysis, and therefore enhances clotting stability (Leenaerts et al., 2018), and also plays a role in activating the complement cascade (Morser et al., 2018; Leung & Morser, 2018).

FAM3D is a cytokine for neutrophils and monocytes in peripheral blood which may directly interact with ACE2 based on their reciprocal ranking. ACE2 is its 2^nd^ ranking ERC. Although ACE2 does not have a significant ERC to F13B (also known as Coagulation Factor XIII B Chain), it is FAM3D’s top-ranking ERC. F13B functions to stabilize clotting through cross-linking of fibrin (Stelzer et al., 2016). Thus, connections of FAM3D to F13B, possibly through physical binding as implied by their high reciprocal ranks, could be important in modulating the dynamics of clot formation.

Blood pressure and vasoconstriction regulation also show functional enrichment in the CRR network. Naturally, ACE2 is a crucial component of the Renin-Angiotensin System (RAS), which converts angiotensin II to angiotensin (1-7). This, in turn, binds to the MAS1 receptor, promoting vasodilation and reduced blood pressure. As seen in Figure 2, MAS1 is part of the ACE2 CRR network. Although not significantly correlated with ACE2 directly, it has significant RR connection to TSHZ3 (ρ = 0.52, FDR = 7.8E-03, ranks 11, 4) and is FAM3D’s 19^th^ ranking ERC (ρ = 0.49, FDR = 1.5E-02). Biologically MAS1 and ACE2 are key elements promoting vasodilation in the renin-angiotensin system (RAS) (Burrell et al., 2004). Thus, the ERC RR network detects biologically significant connections of ACE2 to RAS signaling via the MAS1 receptor of angiotensin-(1-7). Samavati & Uhal (2020) posit that the loss of ACE2 due to SARS-CoV-2 infection reduces MAS1 signaling and increases AT1 & AT2 signaling via higher levels of angiotensin 2, promoting vasoconstriction, fibrosis, coagulation, vascular and cardio injury, and ROS production. Similar arguments are made by Sriram & Insel (2020). ACE2 and MAS1 do not have a signature of protein coevolution, even though they interact indirectly biologically through the short seven amino acid signaling peptide Ang (1-7). This observation is consistent with our model that ERCs (particularly RR ERCs) are driven by direct binding between protein partners. In contrast, MAS1 has significant RR with TSHZ3 (mentioned above). A biological connection between these proteins is not obvious, although the high ERC reciprocal ranks suggest possible binding affinities worth further investigation. Additionally, TMEM63C is one of four proteins that form a direct reciprocal rank ERC association with ACE2 (Figure 2). It functions in osmolarity regulation and like ACE2, interacts with angiotensin II, possibly reducing damage to kidney podocytes (Eisenreich et al., 2020).

FBXL3 has an RR20 connection to FGB and ranks GPR141 in its top 2%. This protein is a component of circadian rhythm regulation (Busino et al., 2007). Many aspects of the cardiovascular system have circadian cycling such as heart rate, blood pressure, and fibrinolysis (Reilly, Westgate & FitzGerald, 2007). Endogenous oscillators in the heart, endothelial cells, and smooth muscles may play significant roles in these cycles (Reilly, Westgate & FitzGerald, 2007), and the CRR network suggests that direct interactions between FBXL3 and FGB could play a role in circadian aspects of fibrinolysis.

CD34 (Hematopoietic Progenitor Cell Antigen CD34) is believed to be an adhesion protein for hematopoietic stem cells in bone marrow and for endothelial cells (Fina et al., 1990). Our ERC analysis indicates connections to coagulation pathway proteins and lipoproteins. In addition to its RR association with FGG (ρ = 0.60, FDR = 2.2.0E-04, ranks 18,9), CD34 also forms significant reciprocal rank correlations with coagulation factor F2 (ρ = 0.69, FDR = 7.9E-06, ranks 1,6), lipoprotein APOE (ρ = 0.64, FDR = 6.0E-05), lipid droplet-associated protein PLIN1 (ρ = 0.64, FDR = 1.1E-04, ranks 8,7), and inflammation associated pentraxin protein PTX3 (ρ = 0.65, FDR = 6.8E-05, ranks 3,11) (Supplementary File S1). As expected from these protein associations, CD34's top enriched term is to complement and coagulation cascade (FDR = 1.4E-08). There is also enrichment for HUVEC cells (FDR = 3.1E-05) and Blood Plasma (FDR = 1.7E-04) (Supplementary File S3).

Consistent with the descriptions above, the CRR network shows enrichment (full enrichment table in Supplementary File S3) for negative regulation of blood coagulation (FDR = 4.3E-08), platelet alpha granule-related terms (FDR = 1.7E-05), plasma cell (FDR = 8.3E-4) and blood clot (FDR = 4.5E-02). These enrichments indicate that the network involves protein interactions related to blood clotting pathways. There are also several significantly enriched terms which are driven in part by ACE2, such as regulation of systemic arterial blood pressure by renin-angiotensin (FDR = 1.6E-03; members: ACE2 and SERPINF2), metabolism of angiotensinogen to angiotensin (FDR = 6.9E-03; members: ACE2 and CPB2), regulation of blood vessel diameter (FDR = 1.5E-02; members: ACE2 and SERPINF2), and renin-angiotensin system (FDR = 1.8E-02; members: ACE2 and MAS1).

### CRR Subnetworks

Here we briefly describe other subnetworks within the CRR network with implications to COVID-19.

#### TMEM63C RR Subnetwork

TMEM63C is one of four proteins that form a direct reciprocal rank ERC association with ACE2 (RR20). It functions in osmolarity regulation. In addition to ACE2, TMEM63C has direct RR20 connections to three proteins, CCDC105, LECT2, and C16orf78, and through them forms a subnetwork also containing TMCO2, ARMC7, PAX4, ETS1, and SUV39H1 (Fig. 3). LECT2 (Leukocyte Cell-derived Chemotaxin 2) is involved in macrophage activation, insulin resistance and diabetes, and neutrophil chemotaxis (Yamagoe et al., 1996; Zhang et al., 2018; Takata et al., 2021). TMEM63C and LECT2 are significantly correlated (ρ = 0.64, FDR = 7.5E-04) with high reciprocal ranks (1,6). Thus, LECT2 is connected to ACE2 through reciprocal ranks between TMEM63C and ACE2.

Little is known about C16orf78, except that it is enriched in testes, and specifically in spermatids (Uhlén et al., 2015). We, therefore, looked at its ERC protein associations as an exploratory tool for possible function. C16orf78 forms a strong RR association with TMEM63C (RR 1,4) and also has reciprocal ERCs with PAX4 (3,9), ETS1 (11,2), and SUV39H1 (10,15). ETS1 is a transcription factor involved in cytokine and chemokine processes. Whereas SUV39H1 is a suppressor of variegation protein whose function loss leads to chromosome instability. It has only 24 significant ERC proteins. These show enrichment for pri-miRNA transcription from RNA polymerase II promoter (FDR = 7.8E-03, members: ETS1;RELA), scavenging by class A receptors Homo sapiens (2.6E-02, members: COL4A1;APOB), endosomal part (FDR = 2.8E-02, members: WDR81;APOB) and striated muscle tissue development (FDR = 4.9E-02; members: IGSF8;BVES). Pri-miRNAs are processed into miRNAs, whereas scavenging class A receptors play a role in innate immunity as phagocytic receptors in macrophages and dendritic cells (Areschoug & Gordon, 2009), which ties to the endosome term enrichment. These observations may serve as a guide for further investigations into C16orf78 function.

There is also little information on CCDC105, except an intriguing paper using phylogenetic profiling to implicate it as functioning in meiosis-specific chromatin and spermatogenesis (Tabach et al., 2013) and an association of a human variant with infertility (Handel & Schimenti, 2010). Consistent with those two studies, enrichment of the top 2% ERCs for CCDC105 has one significant term, from proteomicsDB for spermatozoon (members: UBXN11, CCDC105, C10orf62, MYCBPAP, PMFBP1). It forms strong RR ERCs with the transcription factor PAX4 (3,9) and TMEM63C (1,4).

These findings suggest that TMEM63C protein connections involve innate immunity and spermatogenesis and indicate possible avenues for elucidating interactions of its protein partners of relatively unknown function, such as C16orf78 and CCDC105.

#### TSHZ3 Subnetwork

TSHZ3 does not have a significant ERC to ACE2, yet it connects to ACE2 through FAM3D, with which it has significant reciprocal ranks (1,9). TSHZ3 is a key regulator of airflow and respiratory rhythm control (Caubit et al., 2010), phenotypes that could be important in COVID-19 respiratory distress. Therefore, potential signaling interactions between TSHZ3 and FAMD3, possibly mediated by physical binding, warrant further examination. Additionally, TSHZ3 variants are associated with amyloid-β processing and Alzheimer’s disease (Louwersheimer et al., 2017), and it plays a role in smooth muscle development (Caubit et al., 2008). TSHZ3 is highly connected within the ACE2 CRR network, with 10 reciprocal rank connections (Fig. 3). One of these, BRINP3, connects back to coagulation through FGA and CPB2.

#### L1CAM RR Subnetwork

L1CAM is the fourth protein with a direct RR20 connection to ACE2 (Figure 2). It was originally discovered as an important protein in nervous system development (Moos et al., 1988; Samatov, Wicklein & Tonevitsky, 2016). Among its other functions may be stem cell differentiation to vascular endothelial cells (Rizvanov et al., 2008), and it also plays a role in tumor vascular development (Angiolini & Cavallaro, 2017). These functions may play a role in its protein coevolution with ACE2. Interestingly, BMX is an RR20 to L1CAM and is known as a tyrosine kinase that is present in endothelial and bone marrow cells and may play a role in inflammatory response (Chen et al., 2014). The BMX and ACE2 proteins are encoded by neighboring genes on the X chromosome (Navarro Gonzalez et al., 2021) and BMX has been shown to potentially have two SNPs with strong linkage disequilibrium to an ACE2 SNP associated with the lowered circulation of angiotensin (1-7) (Chen et al., 2018). L1CAM’s top 2% ERC enrichment analysis shows significant terms for coagulation pathway-related (FDR = 5.4E-04), tumor necrosis factor signaling (FDR = 6.4E-03), and gamma-carboxylation of proteins (FDR = 6.7E-03). It connects to FGA through CDKN2C and to FGB through Pfn4.

### C2 The ACE2 Unidirectional Reciprocal Rank (URR) Network

ACE2 also has highly significant ERCs with interacting proteins that are unidirectional, meaning that ACE2 ranks these proteins in its top 2%, but the partner protein does not rank ACE2 within its top 2% due to higher ERC correlations with other partners (Table 1). Some of ACE2’s highest-ranking proteins fall into this category, including GEN1 (rank 1), XCR1 (2), IFNAR2 (5) KIF3B (6), and ITPRIPL2 (7), FAM227A (8), TLR8 (9), COL4A4 (10), F5 (12), and AR (13). To focus on strong protein connections in this set, we took the top ten proteins with unidirectional ERCs for ACE2 and then added their reciprocal rank 20 (RR20) partners. The resulting ACE2 Unidirectional Reciprocal Rank (URR) Network contains 69 proteins (Fig. 4).

**Figure 4.**
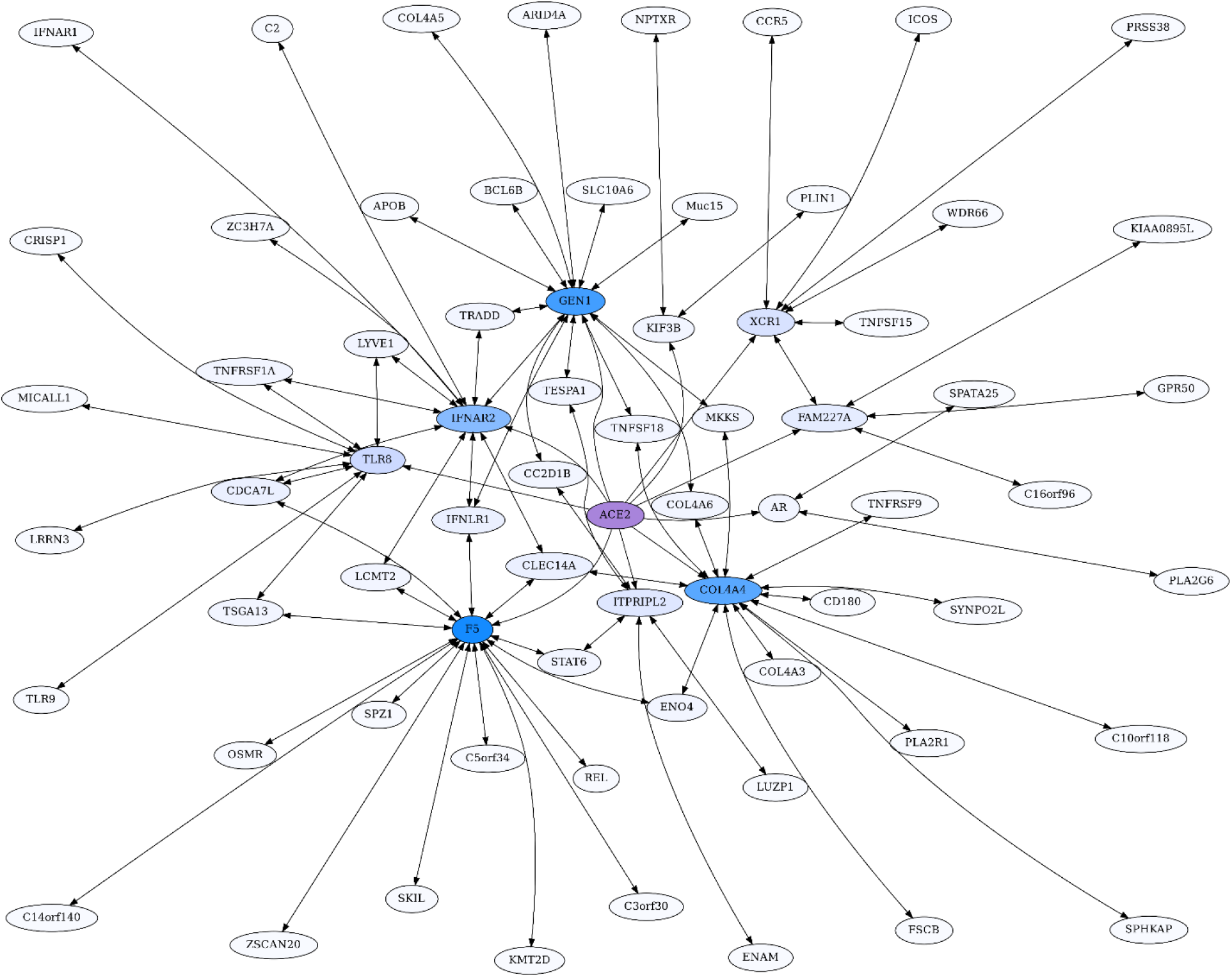
ACE2 Unidirectional Reciprocal Rank (URR) Network. ACE2’s top 10 unidirectional ERC proteins for a web of reciprocal rank (RR20) connections. The network is particularly enriched for cytokine signaling and immunity. Highly interconnected proteins include COL4A5, F5, GEN1, and IFNAR2. ACE2 is highlighted in purple, and blue shading intensity indicates the level of reciprocal connectivity for different proteins.

Notable in the network are many proteins involved in immunity and cytokine signaling, such as IFNAR2 (Interferon alpha/beta receptor 2), XCR1 (X-C Motif Chemokine Receptor 1), and ICOS (Inducible T Cell Costimulator). There are also Toll-Like Receptors TLR8 and TLR9, which stimulate innate immune activity (Forsbach et al., 2011), and Tumor Necrosis Factor related proteins such as TNSFS18, TNTSF15, TNFRSF9, and TNRRSF1A.

Enrichment analysis of the URR network generates 72 significant terms (Supplementary File S3). The network is highly enriched for cytokine-cytokine receptor interaction (FDR = 6.5E-06), I-kappaB kinase/NF-kappaB signaling (FDR = 1.6E-06), necroptosis (FDR = 3.3E-03), viral infections, such as Human Papillomavirus (FDR = 5.7E-04) and Herpes virus (FDR = 3.5E-03), JAK-STAT and PI3K-AKT signaling pathways, Toll-like receptor signaling, and immune system Homo sapiens (FDR = 3.7E-03).

XCR1 is the 2^nd^ highest ACE2 ERC. It is the receptor for chemokine XCL1, which is produced in response to infection and inflammation, and during development of regulatory T cells (Lei & Takahama, 2012). Furthermore, XCR1 maps to a region implicated in severe COVID-19 by a genome-wide association study (Severe Covid-19 GWAS Group, 2020). As seen in Figure 4, XCR1 forms a RR subnetwork with six other proteins (ICOS, CCR5, WDR66, TNSFS15, PRSS38, and FAM227A), three of which are known to be involved in immunity. ICOS (Inducible T Cell Costimulator) is reciprocally evolving with XCR1 based on their ERC interaction. It is an inducible T Cell stimulator that is essential for T helper cell responses (Hutloff et al., 1999; Tafuri et al., 2001). In addition, ICOS signaling is impaired in COVID-19 patients requiring hospitalization (Hanson et al., 2020). The high ERC between ACE2 and XCR1, RR of XCR1 to ICOS, and evidence that both are associated with ICOS pathologies, suggest that disruption of ACE2-XCR1 interaction could have a contributory role. C-C Motif Chemokine Receptor 5 (CCR5) forms a significant RR ERC with XCR1 as well. Several studies have implicated CCR5 variation and expression to be associated with COVID-19 severity (Gómez et al., 2020; Hubacek et al., 2021; Kasela et al., 2021), while others have not (Bernas et al., 2021). TNFSF15 is a third immune response protein in the XCR1 RR subnetwork that shows elevated expression in patients with severe COVID-19 (Jain et al., 2021). We recognize that the involvement of these immune-related proteins does not require an effect mediated through ACE2; however, their protein evolutionary correlations suggest that ACE2 may be part of a signaling network (perhaps modulated through XCR1) that is contributory.

IFNAR2 is another protein that is highly correlated with ACE2 (ρ = 0.62, FDR = 6.1E-04) and is also implicated in severe COVID-19 by GWAS and expression data (Zhang et al., 2020; Liu et al., 2021; Pairo-Castineira et al., 2021). It has RR20 ERCs with ten other proteins and is embedded in a complex web of interactions with members of the ACE2 network. Here we draw attention to a few key features. Notably, IFNAR2 and IFNAR1 are RR partners, as expected given that they combine to form the IFN-alpha/beta receptor, which is the receptor for both alpha and beta interferons. IFNAR2 forms a high RR relationship with TNFRSF1A (ρ = 0.84, FDR = 4.8E-12, 1,1 reciprocal ranks). This protein is the receptor for TNFα and the pathway affects apoptosis and inflammation regulation. Jin et al. (2015) found that ACE2 deletion increases inflammation through TNFRSF1A signaling, lending further support to a functional association between ACE2 and this protein.

GEN1 is the highest-ranking ACE2 ERC protein (ρ = 0.67, FDR = 4.2E-05), and it functions as a resolvase of Holliday junctions and a DNA damage checkpoint signaling (Chan & West, 2015). Frankly, we are perplexed by the functional significance of ACE2-GEN1 correlated evolution. As observed in the ACE2 network, GEN1 is a highly interconnected protein, with 14 RR20 connections in the network. This result suggests that GEN1 may have additional functions beyond DNA replication. Indeed, although its second-highest RR is to CC2D1B (2,1), a protein involved in mitosis, its highest RR is to Interferon Lambda Receptor 1 (IFNLR1), with an impressive Spearman correlation of ρ = 0.89 (FDR = 6.2E-17). As IFNLR1 binds cytokine ligands and stimulates antiviral response, this suggests some feedback mechanism between GEN1 and the immune system, possibly related to its functional role in DNA damage checkpoint signaling.

Indeed, its top 2% ERCs show enrichment for multiple viral infection terms (Supplementary File S3). Therefore, it appears that GEN1 has a “hidden life” that ERC analysis suggests warrants exploration.

The Collagen Type IV A4 subnetwork (Fig. 4, Fig. 5) lends further credence to the view that ERCs can detect proteins with likely binding partners. COL4A4 is a component of the Collagen Type IV protein complexes in basement membranes in the extracellular matrix of various tissues, including the kidney glomerulus and vascular endothelial cells, and lung alveoli (Myllyharju & Kivirikko, 2001). COL4A4, COL4A3, and COL4A5 complex with each other in the basement membranes of kidney glomeruli – mutations in these COL4A proteins are known to cause different kidney disorders (Torra et al., 2004; Wiradjaja, DiTommaso & Smyth, 2010). Consistent with their expected binding, COL4A4 and COL4A3 are each other’s reciprocal best partners (ranks 1,1) and highly correlated with each other (ρ = 0.88, FDR = 4.4E-16). Both show highly significant ERCs to COL4A6 (rank 6,5 for COL4A4 ρ = 0.83, FDR = 2.1E-12; rank 22,30 for COL4A3 ρ = 0.78, FDR = 1.6E-10). Thus, evolutionary rate correlations show highly significant ERCs among Collagen Type IV proteins known to physically interact. A future direction is to use ERCs to more precisely define predicted coevolving protein segments, which could be used to inform docking simulations and experimental studies.

**Figure 5.**
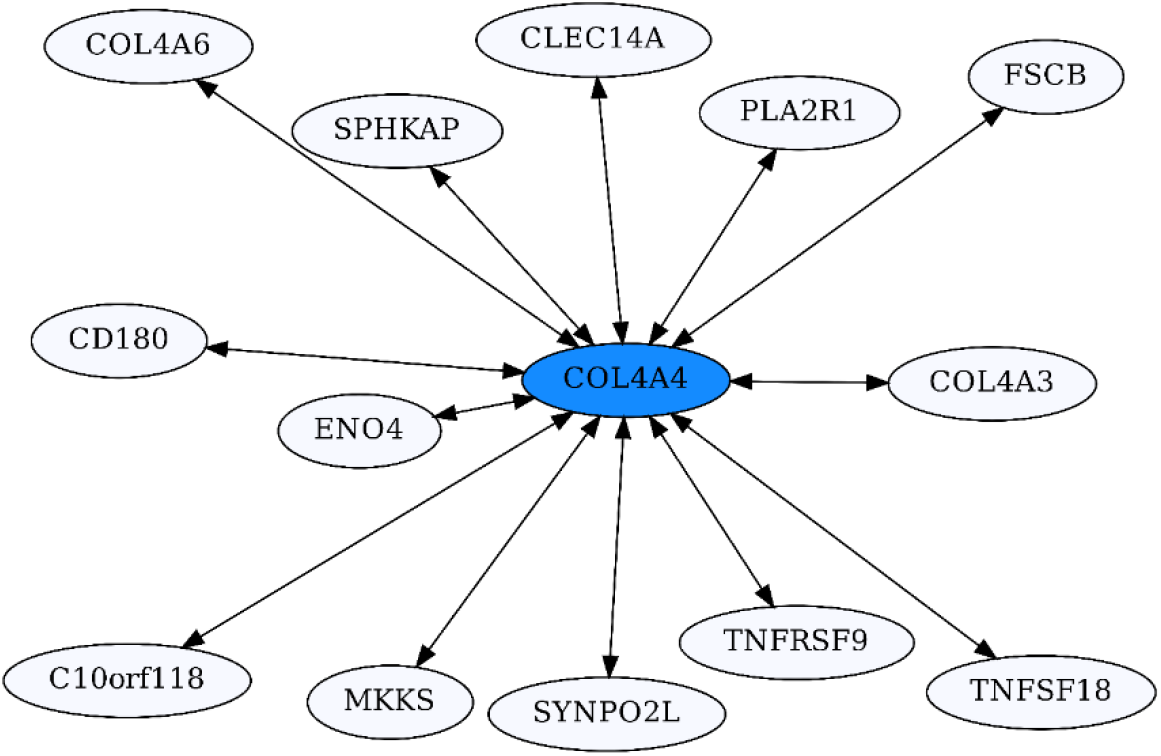
COL4A4-Centric RR20 Network. This network detects reciprocal ERCs of different proteins to COL4A4, including other COL4A proteins known to form complexes with COL4A4.

COL4A5 also has significant ERCs to COL4A3 (ρ = 0.71, FDR = 2.2E-08) and COL4A4 (ρ = 0.71, FDR = 1.7E-08), but these do not qualify as RR20 due to the large number of high ERCs for COL4A5. Interesting, COL4A5-MUC15 are top-ranking partners (ranks 1,1) with a very high ERC (ρ = 0.89, FDR = 3.2E-16). MUC15 is a cell surface protein that is believed to promote cell-extracellular matrix adhesion and it is implicated in affecting influenza infection (Chen et al., 2019), which may increase its relevance in the context of COVID-19 infection. ERCs may help to inform candidate domains within each protein that are involved in their expected binding affinity.

Coagulation Factor V (F5) is known for its role in the coagulation cascade. However, F5 is a highly ERC-connected protein, with 43 proteins ranking it in their respective top 5 highest ERCs. This connectedness is also reflected in the RR20 network shown below (Fig. 6). F5 has 16 direct RR20 connections out of 20 possible. Although F5 is a vital protein in the coagulation cascade, its top 16 RR connections indicate immune functions, including Interferon λ receptor 1 (IFNLR1; RR 4,10) and Oncostatin M Receptor (OSMR; RR 1,4). This is reflected in the enrichments among its 16 RR proteins for the JAK-STAT signaling pathway (FDR = 8.7E-03) and response to cytokine (FDR = 2.5E-02). Similarly, the F5 top 2% ERC show enrichments for 54 terms (Supplementary File S3); notably many related to inflammatory response (FDR = 1.1E-03) and the complement system (FDR = 8.4E-03). The functions of several of F5’s RR20 partners are not well known, such as C14orf140 and C5orf34. Their top 2% enrichment suggests cytokine receptor activity (FDR = 2.7E-02) for C14orf140, and Human Complement System (FDR = 1.9E-03) and cytokine receptor activity (FDR = 2.1E-02) for C5orf34. In conclusion, F5 appears to have a “secret life” of strong protein interactions reflecting moonlighting functions with extensive signaling or modulation roles beyond coagulation regulation.

**Figure 6:**
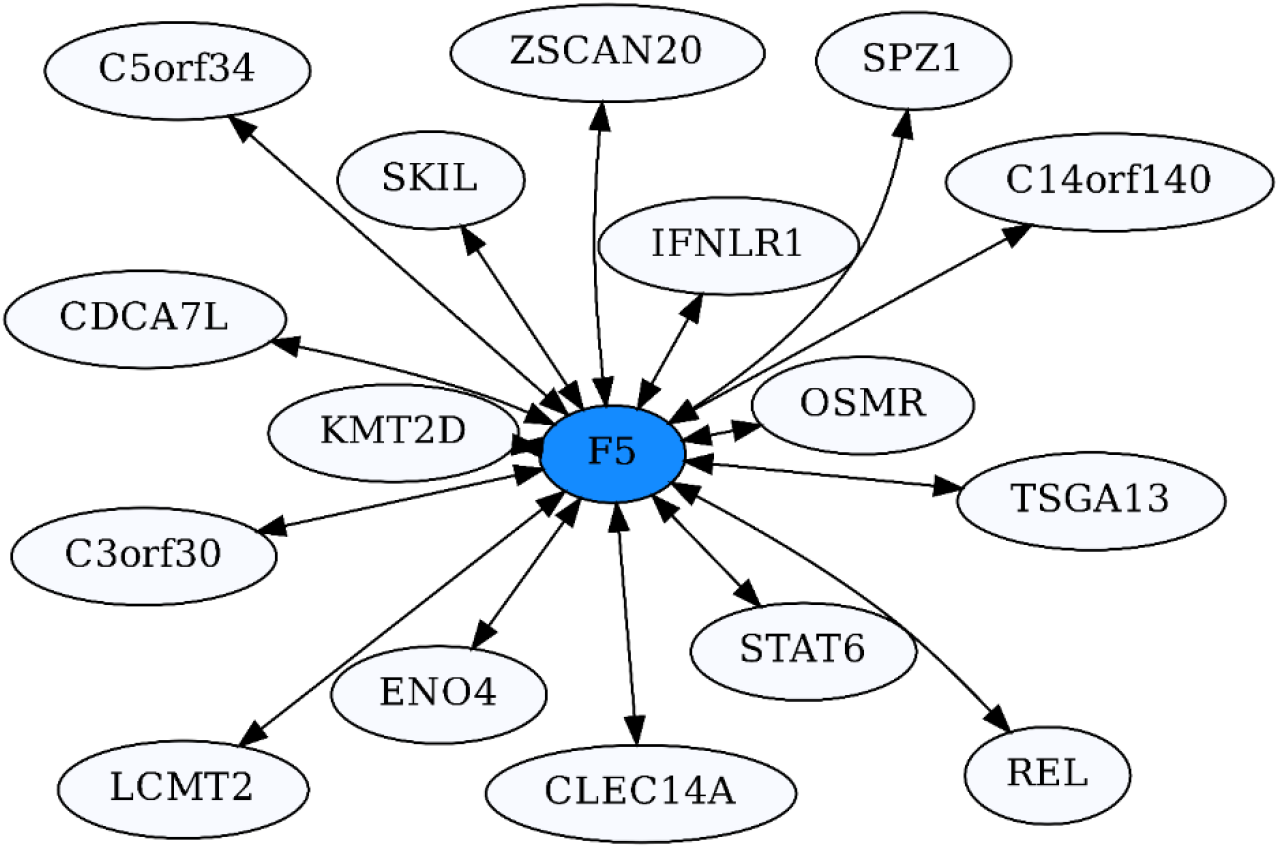
Coagulation factor V-centric RR20 Network. The network captures strong reciprocal ERCs between F5 and proteins related to immune function such as IFNLR1.

### D. ERCs and Protein Interactions

We postulate that ERCs detect proteins that are coevolving due to functional interactions. Furthermore, we propose that physical binding is an important mechanism contributing to significant ERCs between proteins. This is consistent with anecdotal observations from this study of high reciprocal rank ERCs among the fibrinogen components FGA, FGB, & FGG, the Collagen Type IVA proteins COL4A4, COL4A3, and COL4A6 proteins, and Interferon alpha/beta proteins IFNAR2 and IFNAR1.

To further investigate the role of binding affinity, we examined the mammalian protein complex database CORUM (Giurgiu et al., 2019) to determine whether significantly higher Spearman rank correlations (ρ values) are found among proteins within known protein complexes. A set of 139 protein complexes (excluding those with overlapping proteins) were identified which contain at least two members from our ERC data set, for a total of 258 pairwise comparisons. We compared the ρ values of within complex proteins to the median values for proteins outside the complex and found that Spearman rank correlations of within complex proteins were significantly higher than its between complex values according to Wilcoxon matched signs rank tests (WMRST) under a significance level of α = 0.05 (p = 5.2E-04), with a median increase of 6.3% (Supplementary File S11). Many of the complexes contain large numbers of proteins, reducing the probability of direct physical contact between individual members. We therefore also analyzed only proteins from complexes with 5 or fewer members (96 pairs). In this case, the median ρ value increase is 15.8% (WMSRT p = 6.2E-03). The results support the view that proteins within known complexes show higher ERCs than between complexes, and further implicate physical contact as a contributor to ERCs. However, more detailed studies are warranted.

## Discussion

In this paper, we take an exploratory approach to ACE2 protein interactions using evolutionary rate correlations. Our key findings are that the ERC analysis predicts ACE2 to have previously unidentified protein partners, and to be part of interaction networks relevant to COVID-19 pathologies. Most notably, ACE2 forms strong ERC networks relevant to coagulation and immunity. These networks suggest that disruptions in ACE2 function may contribute to COVID-19 pathologies. A potential mechanism for this is a reduced abundance of membrane-bound ACE2 that can disrupt signaling networks. Additionally, the presence of the soluble ACE2 ectodomain may explain the systemic pathologies of COVID-19 infection as its circulation in the blood can affect pathways throughout the body.

An overwhelmingly strong pattern is an association between ACE2, its partners, and the coagulation pathway. This is exemplified by the enrichment for terms related to coagulation pathways in the CRR network, and the presence of the three proteins that form fibrinogen (FGA, FGB, FGG) which constitutes the clotting molecule fibrin. Abnormal clotting and coagulation such as “hypercoagulability” has been observed as a major symptom of COVID-19 infection (Fei et al., 2020). Additionally, disseminated intravascular coagulation (DIC) due to COVID-19 has been found more frequently in fatal cases of COVID-19 than non-fatal cases (Seitz & Schramm, 2020). Levi et al. (2020) have noted that low-grade DIC often seen in COVID-19 is associated with a sudden decrease in plasma fibrinogen before death. This makes the connection with the various fibrinogen subcomponents even more striking. Our network data suggest that ACE2’s connection to fibrinogen is mediated through Clusterin and GPR141 (Fig. 3). The chaperone protein Clusterin’s role in removing misfolded proteins in the blood and its common association with fibrinogen in blood plasma (Wyatt & Wilson, 2010) lend credence to these ERC findings. What remains unclear is the nature of potential functional interactions between ACE2 and Clusterin, but the ERC results suggest that this warrants further attention. The discovery of a strong ERC association of Clusterin and GPR141 is a novel finding, as functional information on GPR141 is largely lacking. ERC analysis indicates that these proteins functionally interact, likely involving coagulation processes.

Another mechanism for ACE2’s influence on the coagulation effects of COVID-19, based on ERCs, is through F5. F5 canonically is activated by the same enzyme (Thrombin) that converts fibrinogen into fibrin for clotting (Omarova et al., 2013). Omarova et al. (2013) further report that inhibition of F5 can enhance an anticoagulant ability of an alternate fibrinogen that utilizes a different isoform of FGG, fibrinogen γ′. Thus, we hypothesize that abnormal coagulation activity may (in part) be driven by disruptions in ACE2-F5 protein interactions, which could reduce anticoagulant feedback mechanisms. F5 is also found to have many significant ERCs outside of the coagulation pathway, connecting to various immunity-related pathways (Fig. 4, Supplementary File S1). The ERC results for GPR141 and F5 reveal how ERC analysis may be useful in providing testable hypotheses for functions of understudied proteins, and to investigate additional functional roles on well-studied proteins.

A second major finding is ACE2 protein-protein interactions that connect to immunity. “Cytokine storms”, an overreaction of the immune system which can lead to inflammation and organ failure, is a second major hallmark of severe COVID-19, and its management is a major target of medical treatment research (Luo et al., 2020; Mangalmurti & Hunter, 2020). Chemokines are a class of cytokines that act as immune cell attractants (Coperchini et al., 2020), and an increase in chemokine production may be characteristic of COVID-19 infection (Coperchini et al., 2020). XCR1 is a receptor of XCL1 chemokines, mostly expressed in dendritic cells, and plays a role in cytotoxic immune responses (Lei & Takahama, 2012). The XCR1 protein, strikingly, is the second-highest ERC to ACE2 and has already been implicated in severe COVID-19 infection (Severe Covid-19 GWAS Group, 2020). While the specific mechanism by which XCR1 might play a role in severe COVID-19 is not yet known, ERC results indicate its role may be mediated by ACE2 with XCR1’s ERCs also possibly indicating a broader functional role in coagulation. Excessive Inflammatory response, particularly as a consequence of cytokine storms, is a clear pathology or COVID-19.

Type 1 interferons are among the first types of cytokines produced after viral infection (García-Sastre & Biron, 2006; Sallard et al., 2020). A component of the type 1 interferon receptor, IFNAR2, is among the strongest ACE2 ERCs, possibly linking ACE2 to the type 1 interferon immunity response. Notably, IFNAR2 has been implicated in severe COVID-19 infection (Pairo-Castineira et al., 2021). Since type 1 interferons have shown some initial efficacy in treating COVID-19 infection (Sallard et al., 2020), it is possible that the SARS-CoV-2 virus interaction with both receptor and soluble ACE2 interferes with type 1 interferon response, as low levels of type 1 interferons have been found in COVID-19 patients (Salman et al., 2021). Another connection of ACE2 with immunity may be mediated by the toll-like receptor TLR8 (a strong ACE2 ERC), among TLRs believed to regulate platelet circulation in response to inflammation (Beaulieu & Freedman, 2010) providing possible avenues for interaction with soluble ACE2 in blood. Genetic variants in TLRs (including TLR8) may affect COVID-19 susceptibility (Lee, Lee & Kong, 2020). Thus, there are many potential avenues for ACE2 protein interactions contributing to immune dysregulation in COVID-19 disease, which may warrant further investigation given the strong ERC associations of ACE2 with proteins relevant to immunity, although the functional bases of such interactions are unknown. Other ACE2 network ERCs of interest are relevant to kidney disease, cardiovascular disease, male fertility, Alzheimer’s disease, and DNA damage checkpoint signaling. These are discussed further in the Supplementary Text.

Overall, the underlying concept behind the evolutionary rate correlation approach (also called evolutionary rate covariance or evolutionary rate coevolution) is that coevolving proteins will show correlated rates of change across evolution and that this reflects functional interactions (Clark, Alani & Aquadro, 2012; Wolfe & Clark, 2015). Clark and colleagues have developed a web interface (https://csb.pitt.edu/erc_analysis/) to screen for ERC interactions for Drosophila, yeast, and mammals. Their mammalian data set is based on 33 mammalian species (Priedigkeit, Wolfe & Clark, 2015; Wolfe & Clark, 2015). We have compared their output for ACE2 to our analyses and found only one overlapping protein (XCR1) between their significant ERCs (p < 0.05) and our top 2% ACE2 ERCs. There are many methodological differences between our approaches, including the number and specific mammalian taxa used, the method for calculating protein rates, and the phylogeny used for calculating branch lengths. In addition, their dataset includes 17,487 proteins, whereas our analysis is currently restricted to 1,953 proteins for which we were confident about 1:1 orthology and therefore for which there are minimal paralogy complications. Furthermore, we are uncertain how their database dealt with potential short branch artifacts on ERC calculations. In our case, we found that short branches in the phylogeny resulted in significant correlations between branch time and protein rate, thus both inflating estimated ERCs and introducing branch time as a confounding factor which can lead to spurious correlations, and we removed these by branch trimming. In another study, Braun et al. (2020) applied a “phylogenetic profiling” approach to identify ACE2 interacting proteins relevant to possible drug targets for COVID-19. Phylogenetic profiling generally screens multiple genomes for presence-absence correlations of protein combinations, as a method to detect candidate protein interactions (Pellegrini et al., 1999). However, Braun et al. (2020) use a modification of the method that also incorporates a BLAST-based distance metric from human ACE2 across taxa ranging from humans to fungi. We find no overlap in our findings and their results, and suggest that our direct measures of protein evolutionary rates may be a more sensitive approach for finding evolutionary interactions among proteins in mammals. Obviously, future validation studies are needed to reveal which approaches are most effective at detecting candidate protein interactions, or whether each has its own merits for detection of different interactions.

Experimental validations of novel ACE2 protein associations predicted by our ERC approach are clearly needed. A necessary first step is to establish whether ACE2 has binding affinities *in vitro* and *in vivo* with proteins showing high evolutionary correlation to it, in particular CLU, XCR1, GEN1, and IFNAR2. Similar binding affinity is predicted between CLU and GPR141 based on their high reciprocal rank ERCs. CLU-FGG and GPR141-FGB provide connections to fibrinogen based on their evolutionary correlations, suggesting binding affinities. Applicable methods could include protein complex immunoprecipitation, tagged protein analysis, and yeast-two-hybrid analysis (Rao et al., 2014).

We have begun preliminary analyses using short (10mer) amino acid sequences to identify predicted sites of interaction among protein partners. These data may be able to inform protein-protein docking simulations for protein pairs using software that allows for the incorporation of a priori predicted interfaces (Van Zundert et al., 2016; Pagadala, Syed & Tuszynski, 2017). For example, these 10mer analyses can be used to determine likely regions of binding affinity between ACE2 and Clusterin, for experimental validation through mutational analysis. Similarly, coagulation factor V shows high ERCs for non-canonical proteins, which can be investigated to determine whether F5 has novel functions outside of the coagulation pathway.

Further computational analyses of ERCs are needed to better understand their relationship to protein function and evolution. For instance, machine learning and simulation approaches can be used to determine which aspects of protein structure, amino acid properties, and rates of protein evolution improve ERC predictive power. We are currently expanding the mammalian protein set for such analyses. Finally, if evidence mounts that ERCs can be informative in predicting protein interactions, the approach can be applied more broadly as an additional tool for detecting protein interaction networks involved in biological processes and disease.

## Methods

### Taxon Selection and Data Collection

Our evolutionary rate correlation (ERC) approach requires orthologous protein sequence data across a large number of taxa with well-defined phylogenetic relationships. Calculation of evolutionary rates requires a resolved phylogeny of the taxa analyzed that is scaled to evolutionary time. ERC calculations utilize the TimeTree (Kumar et al., 2017) service to generate a time-scaled phylogenetic tree using the mammalian taxa that are represented in OrthoDB sequence data (Supplementary Fig. S1). The tree generated is in units of millions of years and is based on a compilation of thousands of phylogenetic-dating related studies. The tree is utilized as a base topology in phylogenetic analysis and its branch lengths are used to measure time for calculating evolutionary rates from the resultant protein trees (Supplementary Fig. S2).

Well-defined orthologous sequence data is sourced from OrthoDB (Kriventseva et al., 2019) at the “mammalia” (taxonomic id: 40674) taxonomic level. Since OrthoDB sequence data is gathered from a variety of sources and clustered algorithmically (unsupervised), primarily based on sequence similarity (Kriventseva et al., 2015), related paralogous proteins are often clustered with each other even if canonically annotated as functionally distinct proteins. Additionally, since the data sources for sequences can have varying levels of completeness, most ortholog groups on OrthoDB are missing sequence data for one or more taxa represented in the database. So, a majority of the data we selected is from single-copy ortholog groups with at least 90 of the 108 possible mammalian taxa present. This set is fairly restrictive, however, so proteins with a possibly relevant function in COVID-19 pathologies (such as XCR1, and IFNAR2) in ortholog groups that do not meet the initial criteria (but with minimal paralogy issues) were included with paralogous sequences manually disambiguated based on published protein annotations. If a taxon in a given sequence has duplicate sequences that cannot be disambiguated, the taxon is excluded in phylogenetic and ERC calculations for the specific proteins involved. In total, 1,953 orthologous protein groups are used in analyses. Additional details on the data set are provided in the Supplementary Text.

### Phylogenetic Calculations and Protein Alignments

To prepare orthologous sequence data for ERC calculation, each set of protein sequences are first aligned using the MAFFT software package (Katoh & Standley, 2013) using the following arguments: “--maxiterate 1000 --localpair --anysymbol”. Since the sequences come from data sources with varying levels of quality and multiple alignment programs can be imperfect, the aligned sequences must then be trimmed. The alignments are trimmed using the trimAl software package (Capella-Gutiérrez, Silla-Martínez & Gabaldón, 2009) using the “-automated1” argument to remove poorly aligned regions. These final prepared alignments are then used to generate maximum-likelihood phylogenies. The IQ-TREE software package (Minh et al., 2020) is used to estimate protein branch lengths (equivalent to average substitution counts per site). Specifically, the “LG+F+G+I” model (which utilizes an empirically derived amino acid substitution matrix) is used with the following additional parameters: “-B 1000 -st AA -seed 1234567890” and the TimeTree phylogeny is provided to constrain output tree topology to reduce possible branch length estimation errors with the “-g” option. These trees are the basis of ERC calculations. Protein branch lengths are based on the average number of changes in amino acids at each residue in the alignment. The resultant branch lengths are paired with corresponding branches in the TimeTree to quantify branch-specific rates to be used for ERC calculations (described below).

### Calculation of ERCs

Our evolutionary rate correlation (ERC) method is designed to predict protein-protein interactions using evolutionary data (Yan, Ye & Werren, 2019), and is based on protein evolutionary rates on terminal branches of the mammalian phylogeny (Fig. 1). Specifically, we use a reduced mammalian phylogeny consisting of 60 taxa for the final ERC. This is because ERCs calculated with a more complete phylogeny (108 species) had short branch problems that inflate ERC spearman rank correlations (discussed in Supplementary Text). Most notably, there was an association between branch time and protein rate for many proteins, with taxonomic oversampling (e.g. in Primates and Rodentia) leading to many ERCs being driven by relatively short branches (Supplementary Text). We attempted to control for these effects initially by using a partial correlation method, but found that it was not sufficient (Supplementary Text). We then removed taxa that contributed short branches in our phylogeny based on either a 20MY or 30MY divergence time threshold and recalculated branch rates for all proteins. We found that the 30MY threshold short branch removal eliminated significant branch time to protein rate correlations for the majority of proteins (96%). The resultant rate data no longer has branch time to branch rate as a confounding cofactor, and the ERCs themselves are no longer biased by extremely short branches and taxonomic oversampling (Supplementary Text). Using the adjusted data set, ERCs are calculated for every possible combination of protein pairs for which a tree has been generated. Every protein pair for which an ERC is calculated has each respective tree and the TimeTree topology is pruned to only include the intersection of shared taxa between the two, using the “ETE3” Python package (Huerta-Cepas, Serra & Bork, 2016). ETE3 is also used to extract the terminal branch lengths of each pruned protein tree and the TimeTree. Evolutionary rates are calculated by dividing the terminal protein-tree branch lengths (average substitutions per site) by the corresponding branch in the TimeTree (measured in millions of years). Terminal branches are used for calculations as they do not have shared evolutionary histories and are therefore independent. The resulting rates have the unit of average substitution per site per millions of years. Given the resultant rates, evolutionary rate correlations can then be calculated by performing a Spearman’s rank correlation test (Yan, Ye & Werren, 2019) using the Python package “SciPy” (Virtanen et al., 2020).

### Multiple Test Corrections

P-values are corrected using the Benjamini-Hochberg FDR multiple-test correction procedure implemented in the Python package “statsmodels” (Seabold & Perktold, 2010). The FDR correction is applied to each respective protein’s set of ERCs. Correlation test results are non-directional, but FDR corrections are dependent on the rank of each correlation’s p-values. Since the rank of each correlation test value on respective protein lists vary, the FDR-corrected p-values of a given protein pair can differ depending on the specific ERC set. An ERC is considered significant if the FDR-corrected p-value is less than 0.05.

### ERC Set Enrichment Analysis

To summarize the common biological function of proteins that tend to have strong ERCs to a protein(s) of interest, gene set enrichment analysis is performed on the top 2% of ERCs (by ρ) of each protein(s) of interest (including the protein itself), including only proteins with ERCs that are significant following an FDR correction at a significance level of 0.05. At most, a protein of interest will have 41 proteins included for enrichment analysis (including itself) (2% of the total 1953 proteins plus self). Protein set enrichment analyses are performed using the Enrichr service (Xie et al., 2021) via the Python bindings provided by the “GSEApy” Python package (Fang et al., 2021) given the background of the full set of 1,953 proteins. We calculate enrichment results for ACE2 and all of its top 20 ERC partners. Additional enrichment analyses were also performed on a case-by-case basis based on relevance, including the reciprocal rank networks. Enrichments are performed using selected relevant term databases: KEGG_2019_Human, GO_Biological_Process_2018, GO_Cellular_Component_2018, GO_Molecular_Function_2018, Reactome_2016,WikiPathways_2019_Human, Tissue_Protein_Expr_from_Human_Proteome_Map, Tissue_Protein_Expr_from_ProteomicsDB, and Jensen_TISSUES.

All enrichment results for terms that are significant at FDR-adjusted p < 0.05 for all analyses are placed into a single table, organized by the enrichment analysis focus (Supplementary File S3). The outputs from different databases can contain redundant terms to each other, so only the most significant of the redundant terms are reported for any enrichment analysis in the main text.

### Reciprocal Rank Network (RRN) Generation

To evaluate and visualize the strongest ERCs centered around proteins of interest, “reciprocal rank networks” (RRNs) are produced. Reciprocal ranks refer to the fact that a significant ERC between two proteins can have different ranks in the two respective protein ERC lists because some proteins have more and higher ERCs than others. To focus on networks of proteins with strong reciprocal rank correlations, we have constructed networks based on proteins with reciprocal ranks of 20 or less (RR20), which is the top 1% in each protein’s highest ERCs based on ρ values. Specifically, we have developed an ACE2 centric reciprocal rank network by the following steps (1) for ACE2, select its top 1% (20) proteins, (2) for each of those proteins, select additional proteins in their ERC list with reciprocal rank 20 or less, and then (3) Given the core set of proteins generated in the previous two steps, connect proteins which have a unidirectional rank of 20 or less.

The resultant network represents the strongest ERCs centered around a protein of interest (in this case ACE2), along with the immediate neighborhood of the strongest ERCs surrounding the protein of interest. The ACE2 Core Reciprocal Rank (**CRR**) was initiated with four proteins to which it has RR20 ERCs: CLU, TMEM63C, FAM3D, and L1CAM), with GPR141 add due to its RR1 strong connection to CLU and unidirectional connection to ACE2. A similar approach has been used to generate an ACE2 reciprocal network, initiated with the top 10 proteins to which ACE2 has highly significant ERCs, but that are not RR20, because the partner ranks other proteins higher in its ERC list, with subsequent one cycle of RR20 built upon these. This ACE2 Unidirectional Reciprocal Rank Network” (**URR**) indicates strong network connections to ACE2 through initial non-reciprocal ranking. Steps 2-3 were omitted as the network becomes extremely large following just the first step, and our focus is on close connections to ACE2 based on ERC analysis.

### ERCs Within and Between Protein Complexes

To compare whether calculated ERCs are stronger between known interactions versus non-interactions, the protein complex database, CORUM (Giurgiu et al., 2019), was used to retrieve known complexes. The “Core Complex” dataset was downloaded and filtered for human complexes to eliminate redundancy, resulting in 233 protein complexes from this CORUM data set which have two or more components present in our 1,953 protein ERC set, representing 258 pairwise ERC comparisons. As these protein complexes have redundancy (i.e. some complexes contain overlapping protein pairs), the set was further restricted to complexes containing unique protein components—resulting in 139 effective unique complexes considered. To test whether ERCs within complexes are higher than between complexes, all pairwise ERCs within complexes were compared to the median ρ value for each pair to proteins present in non-redundant CORUM set that are not in complex with either of these proteins. A Wilcoxon matched signed-rank test was performed using the “wilcox.test” function in base R (version 3.6.1; with parameters “paired” and “exact” set to “TRUE”) on the in-complex ρ values and the median out-of-complex ρ values, to test if the in-complex ρ values were significantly greater than the median out-of-complex ρ values. In addition, as there were many complexes with a majority of subcomponents not present in our 1,953 datasets, the likelihood of individual pairs directly interacting within the complex decreases with the increasing number of proteins in a complex. Therefore, an additional Wilcoxon matched signed-rank test was performed on members of protein complexes composed of five or fewer proteins.

### Testing Whether Branch Rates Increase When Extending Branch Time Within Clades

To test whether increasing branch length results in increasing protein evolutionary rate, we selected separate phylogenetic groups (clades) from the full phylogeny (Supplementary Fig. S1) that contain short branch lengths. Protein evolutionary rate was calculated for each protein on the short branch, and then sequentially extended and recalculated after removing adjacent taxa to extend the branch internally (Supplementary Fig. S5). In this way, the protein evolutionary rate was examined as branches are extended internally in independent clades within the tree. Comparing original branches to the 20MY correction resulted in 14 taxa for which time scales would change, there are 12 taxa for which time scales change between 20MY and 30MY corrections, and there are 16 taxa for which time scales change between 0MY and 30MY. Tests on each branch’s rate against the respective adjusted rate were performed using two-tailed Wilcoxon Matched Signed Rank Tests (Base R v3.6.1), first for proteins of interest (e.g. ACE2) and then for the full protein set. Results are described in the Supplementary text.

## Data Availability

Additional text, figures, and tables are available in the Supplementary Text, as are links to large tables and files deposited at FigShare (https://doi.org/10.6084/m9.figshare.14637450). Code for the ERC pipeline and additional R analyses are available on GitHub: https://github.com/austinv11/ERC-Pipeline.

## Acknowledgments

We thank Zhichao Yan for guidance with the mammalian protein database and discussions on ERC methods and independent contrasts. Also thanked are J. Fay, J. Fry, D. Goldfarb, J. Jaenike, A. Kingsley, A. Larracuente, E. Sia, S. Ghaemmaghami for discussions and helpful feedback, and M. Tsuchiya for support and helpful discussions. Special thanks also go to Nathaniel and Helen Wisch for their support.

## Author Contributions

JHW conceived and supervised the project. AAV conducted the computational analyses and developed figures, tables, and pipeline code for the project. AAV, SC, and JHW contributed ideas and analyses during the development of the project. SC conducted aspects relating to phylogenetic analysis and protein evolution. JHW and AAV wrote drafts of the main text and supplement with input from SC.

## Additional Information

### Supplementary Information

Supplementary Text, Figures and Tables and links to large files are available in Supplementary Information

## References

Aase K, Ernkvist M, Ebarasi L, Jakobsson L, Majumdar A, Yi C, Birot O, Ming Y, Kvanta A, Edholm D, Aspenström P, Kissil J, Claesson-Welsh L, Shimono A, Holmgren L. 2007. Angiomotin regulates endothelial cell migration during embryonic angiogenesis. Genes and Development 21:2055–2068. DOI: 10.1101/gad.432007.

Angiolini F, Cavallaro U. 2017. The pleiotropic role of L1CAM in tumor vasculature. International Journal of Molecular Sciences 18:254. DOI: 10.3390/ijms18020254.

Areschoug T, Gordon S. 2009. Scavenger receptors: Role in innate immunity and microbial pathogenesis. Cellular Microbiology 11:1160–1169. DOI: 10.1111/j.1462-5822.2009.01326.x.

Bastolla U. 2020. Mathematical model of SARS-Cov-2 propagation versus ACE2 fits COVID-19 lethality across age and sex and predicts that of SARS, supporting possible therapy. arXiv Preprint.

Bastolla U, Chambers P, Abia D, García-Bermejo M-L, Fresno M. 2021. Is Covid-19 severity associated with ACE2 degradation ? arXiv Preprint.

Beaulieu LM, Freedman JE. 2010. The role of inflammation in regulating platelet production and function: Toll-like receptors in platelets and megakaryocytes. Thrombosis Research 125:205–209. DOI: 10.1016/j.thromres.2009.11.004.

Bernas SN, Baldauf H, Wendler S, Heidenreich F, Lange V, Hofmann JA, Sauter J, Schmidt AH, Schetelig J. 2021. CCR5∆32 mutations do not determine COVID-19 disease course. International Journal of Infectious Diseases 105:653–655. DOI: 10.1016/j.ijid.2021.02.108.

Böhm S, Szakal B, Herken BW, Sullivan MR, Mihalevic MJ, Kabbinavar FF, Branzei D, Clark NL, Bernstein KA. 2016. The budding yeast ubiquitin protease Ubp7 is a novel component involved in s phase progression. Journal of Biological Chemistry 291:4442–4452. DOI: 10.1074/jbc.M115.671057.

Bonaventura A, Vecchié A, Dagna L, Martinod K, Dixon DL, Van Tassell BW, Dentali F, Montecucco F, Massberg S, Levi M, Abbate A. 2021. Endothelial dysfunction and immunothrombosis as key pathogenic mechanisms in COVID-19. Nature Reviews Immunology 21:319–329. DOI: 10.1038/s41577-021-00536-9.

Bouaziz JD, Duong TA, Jachiet M, Velter C, Lestang P, Cassius C, Arsouze A, Domergue Than Trong E, Bagot M, Begon E, Sulimovic L, Rybojad M. 2020. Vascular skin symptoms in COVID-19: a French observational study. Journal of the European Academy of Dermatology and Venereology 34:e451–e452. DOI: 10.1111/jdv.16544.

Bratt A, Birot O, Sinha I, Veitonmäki N, Aase K, Ernkvist M, Holmgren L. 2005. Angiomotin regulates endothelial cell-cell junctions and cell motility. Journal of Biological Chemistry 280:34859–34869. DOI: 10.1074/jbc.M503915200.

Braun M, Sharon E, Unterman I, Miller M, Shtern AM, Benenson S, Vainstein A, Tabach Y. 2020. ACE2 Co-evolutionary Pattern Suggests Targets for Pharmaceutical Intervention in the COVID-19 Pandemic. iScience 23:101384. DOI: 10.1016/j.isci.2020.101384.

Brunette GJ, Jamalruddin MA, Baldock RA, Clark NL, Bernstein KA. 2019. Evolution-based screening enables genome-wide prioritization and discovery of DNA repair genes. Proceedings of the National Academy of Sciences of the United States of America 116:19593–19599. DOI: 10.1073/pnas.1906559116.

Burrell LM, Johnston CI, Tikellis C, Cooper ME. 2004. ACE2, a new regulator of the renin-angiotensin system. Trends in Endocrinology and Metabolism 15:166–169. DOI: 10.1016/j.tem.2004.03.001.

Busino L, Bassermann F, Maiolica A, Lee C, Nolan PM, Godinho SIH, Draetta GF, Pagano M. 2007. SCFFbxl3 controls the oscillation of the circadian clock by directing the degradation of cryptochrome proteins. Science 316:900–904. DOI: 10.1126/science.1141194.

Capella-Gutiérrez S, Silla-Martínez JM, Gabaldón T. 2009. trimAl: A tool for automated alignment trimming in large-scale phylogenetic analyses. Bioinformatics 25:1972–1973. DOI: 10.1093/bioinformatics/btp348.

Caubit X, Lye CM, Martin E, Coré N, Long DA, Vola C, Jenkins D, Garratt AN, Skaer H, Woolf AS, Fasano L. 2008. Teashirt 3 is necessary for ureteral smooth muscle differentiation downstream of SHH and BMP4. Development 135:3301–3310. DOI: 10.1242/dev.022442.

Caubit X, Thoby-Brisson M, Voituron N, Filippi P, Bévengut M, Faralli H, Zanella S, Fortin G, Hilaire G, Fasano L. 2010. Teashirt 3 regulates development of neurons involved in both respiratory rhythm and airflow control. Journal of Neuroscience 30:9465–9476. DOI: 10.1523/JNEUROSCI.1765-10.2010.

Chan YW, West S. 2015. GEN1 promotes Holliday junction resolution by a coordinated nick and counter-nick mechanism. Nucleic Acids Research 43:10882–10892. DOI: 10.1093/nar/gkv1207.

Chen X-L, Qiu L, Wang F, Liu S. 2014. Current understanding of tyrosine kinase BMX in inflammation and its inhibitors. Burns & Trauma 2:121. DOI: 10.4103/2321-3868.135483.

Chen ZG, Wang ZN, Yan Y, Liu J, He TT, Thong KT, Ong YK, Chow VTK, Tan K Sen, Wang DY. 2019. Upregulation of cell-surface mucin MUC15 in human nasal epithelial cells upon influenza A virus infection. BMC Infectious Diseases 19:622. DOI: 10.1186/s12879-019-4213-y.

Chen C, Zhang XR, Ju ZY, He WF. 2020. Advances in the research of mechanism and related immunotherapy on the cytokine storm induced by coronavirus disease 2019. Zhonghua shao shang za zhi = Zhonghua shaoshang zazhi = Chinese journal of burns 36:471–475. DOI: 10.3760/cma.j.cn501120-20200224-00088.

Chen YY, Zhang P, Zhou XM, Liu D, Zhong JC, Zhang CJ, Jin LJ, Yu HM. 2018. Relationship between genetic variants of ACE2 gene and circulating levels of ACE2 and its metabolites. Journal of Clinical Pharmacy and Therapeutics 43:189–195. DOI: 10.1111/jcpt.12625.

Clark NL, Alani E, Aquadro CF. 2012. Evolutionary rate covariation reveals shared functionality and coexpression of genes. Genome Research 22:714–720. DOI: 10.1101/gr.132647.111.

Colgren J, Nichols SA. 2019. Evolution as a guide for experimental cell biology. PLoS Genetics 15:e1007937. DOI: 10.1371/journal.pgen.1007937.

Connors JM, Levy JH. 2020. COVID-19 and its implications for thrombosis and anticoagulation. Blood 135:2033–2040. DOI: 10.1182/BLOOD.2020006000.

Coperchini F, Chiovato L, Croce L, Magri F, Rotondi M. 2020. The cytokine storm in COVID-19: An overview of the involvement of the chemokine/chemokine-receptor system. Cytokine and Growth Factor Reviews 53:25–32. DOI: 10.1016/j.cytogfr.2020.05.003.

Deng Q, Rasool R ur, Russell RM, Natesan R, Asangani IA. 2021. Targeting androgen regulation of TMPRSS2 and ACE2 as a therapeutic strategy to combat COVID-19. iScience 24:102254. DOI: 10.1016/j.isci.2021.102254.

Dolan ME, Hill DP, Mukherjee G, McAndrews MS, Chesler EJ, Blake JA. 2020. Investigation of COVID-19 comorbidities reveals genes and pathways coincident with the SARS-CoV-2 viral disease. Scientific Reports 10:1–11. DOI: 10.1038/s41598-020-77632-8.

Eisenreich A, Orphal M, Böhme K, Kreutz R. 2020. Tmem63c is a potential pro-survival factor in angiotensin II-treated human podocytes. Life Sciences 258. DOI: 10.1016/j.lfs.2020.118175.

Fang Z, Wolf A, Liao Y, McKay A, Fröhlich F, Kimmel J, Xiaohui L, sorrge. 2021. zqfang/GSEApy: gseapy-v0.10.3. DOI: 10.5281/ZENODO.4553090.

Fei Y, Tang N, Liu H, Cao W. 2020. Coagulation dysfunction: A hallmark in COVID-19. Archives of Pathology and Laboratory Medicine 144:1223–1229. DOI: 10.5858/arpa.2020-0324-SA.

Fina L, Molgaard H, Robertson D, Bradley N, Monaghan P, Delia D, Sutherland D, Baker M, Greaves M. 1990. Expression of the CD34 gene in vascular endothelial cells. Blood 75:2417–2426. DOI: 10.1182/blood.v75.12.2417.2417.

Findlay GD, Sitnik JL, Wang W, Aquadro CF, Clark NL, Wolfner MF. 2014. Evolutionary Rate Covariation Identifies New Members of a Protein Network Required for Drosophila melanogaster Female Post-Mating Responses. PLoS Genetics 10:e1004108. DOI: 10.1371/journal.pgen.1004108.

Forsbach A, Samulowitz U, Völp K, Hofmann HP, Noll B, Tluk S, Schmitz C, Wader T, Müller C, Podszuweit A, Lohner A, Curdt R, Uhlmann E, Vollmer J. 2011. Dual or triple activation of TLR7, TLR8, and/or TLR9 by single-stranded oligoribonucleotides. Nucleic Acid Therapeutics 21:423–436. DOI: 10.1089/nat.2011.0323.

Fricke-Galindo I, Falfán-Valencia R. 2021. Genetics Insight for COVID-19 Susceptibility and Severity: A Review. Frontiers in Immunology 12. DOI: 10.3389/fimmu.2021.622176.

García-Sastre A, Biron CA. 2006. Type 1 interferons and the virus-host relationship: A lesson in détente. Science 312:879–882. DOI: 10.1126/science.1125676.

Giurgiu M, Reinhard J, Brauner B, Dunger-Kaltenbach I, Fobo G, Frishman G, Montrone C, Ruepp A. 2019. CORUM: The comprehensive resource of mammalian protein complexes - 2019. Nucleic Acids Research 47:D559–D563. DOI: 10.1093/nar/gky973.

Godin SK, Meslin C, Kabbinavar F, Bratton-Palmer DS, Hornack C, Mihalevic MJ, Yoshida K, Sullivan M, Clark NL, Bernstein KA. 2015. Evolutionary and functional analysis of the invariant SWIM domain in the conserved Shu2/SWS1 protein family from Saccharomyces cerevisiae to Homo sapiens. Genetics 199:1023–1033. DOI: 10.1534/genetics.114.173518.

Gómez J, Cuesta-Llavona E, Albaiceta GM, García-Clemente M, López-Larrea C, Amado-Rodríguez L, López-Alonso I, Hermida T, EnrÍquez AI, Gil H, Alonso B, Iglesias S, Suarez-Alvarez B, Alvarez V, Coto E. 2020. The CCR5-delta32 variant might explain part of the association between COVID-19 and the chemokine-receptor gene cluster. medRxiv:2020.11.02.20224659. DOI: 10.1101/2020.11.02.20224659.

Gotluru C, Roach A, Cherry SH, Runowicz CD. 2021. Sex, Hormones, Immune Functions, and Susceptibility to Coronavirus Disease 2019 (COVID-19)-Related Morbidity. Obstetrics and gynecology 137:423–429. DOI: 10.1097/AOG.0000000000004275.

Gupta A, Madhavan M V., Sehgal K, Nair N, Mahajan S, Sehrawat TS, Bikdeli B, Ahluwalia N, Ausiello JC, Wan EY, Freedberg DE, Kirtane AJ, Parikh SA, Maurer MS, Nordvig AS, Accili D, Bathon JM, Mohan S, Bauer KA, Leon MB, Krumholz HM, Uriel N, Mehra MR, Elkind MSV, Stone GW, Schwartz A, Ho DD, Bilezikian JP, Landry DW. 2020. Extrapulmonary manifestations of COVID-19. Nature Medicine 26:1017–1032. DOI: 10.1038/s41591-020-0968-3.

Handel MA, Schimenti JC. 2010. Genetics of mammalian meiosis: Regulation, dynamics and impact on fertility. Nature Reviews Genetics 11:124–136. DOI: 10.1038/nrg2723.

Hanson A, Cohen H, Wang H, Shekhar N, Shah C, Dhaneshwar A, Harvey BW, Murray R, Harvey CJ. 2020. Impaired ICOS signaling between Tfh and B cells distinguishes hospitalized from ambulatory CoViD-19 patients. medRxiv:2020.12.16.20248343. DOI: 10.1101/2020.12.16.20248343.

Hubacek JA, Dusek L, Majek O, Adamek V, Cervinkova T, Dlouha D, Pavel J, Adamkova V. 2021. CCR5Δ32 Deletion as a Protective Factor in Czech First-Wave COVID-19 Subjects. Physiological Research 70:111–115. DOI: 10.33549/physiolres.934647.

Huerta-Cepas J, Serra F, Bork P. 2016. ETE 3: Reconstruction, Analysis, and Visualization of Phylogenomic Data. Molecular Biology and Evolution 33:1635–1638. DOI: 10.1093/molbev/msw046.

Hutloff A, Dittrich AM, Beier KC, Eljaschewitsch B, Kraft R, Anagnostopoulos I, Kroczek RA. 1999. ICOS is an inducible T-cell co-stimulator structurally and functionally related to CD28. Nature 397:263–266. DOI: 10.1038/16717.

Itakura E, Chiba M, Murata T, Matsuura A. 2020. Heparan sulfate is a clearance receptor for aberrant extracellular proteins. Journal of Cell Biology 219. DOI: 10.1083/JCB.201911126.

Jain R, Ramaswamy S, Harilal D, Uddin M, Loney T, Nowotny N, Alsuwaidi H, Varghese R, Deesi Z, Alkhajeh A, Khansaheb H, Alsheikh-Ali A, Abou Tayoun A. 2021. Host transcriptomic profiling of COVID-19 patients with mild, moderate, and severe clinical outcomes. Computational and Structural Biotechnology Journal 19:153–160. DOI: 10.1016/j.csbj.2020.12.016.

Jeffery CJ. 1999. Moonlighting proteins. Trends in Biochemical Sciences 24:8–11. DOI: 10.1016/S0968-0004(98)01335-8.

Jin HY, Chen LJ, Zhang ZZ, Xu Y Le, Song B, Xu R, Oudit GY, Gao PJ, Zhu DL, Zhong JC. 2015. Deletion of angiotensin-converting enzyme 2 exacerbates renal inflammation and injury in apolipoprotein E-deficient mice through modulation of the nephrin and TNF-alpha-TNFRSF1A signaling. Journal of Translational Medicine 13:1–16. DOI: 10.1186/s12967-015-0616-8.

Jones VG, Mills M, Suarez D, Hogan CA, Yeh D, Bradley Segal J, Nguyen EL, Barsh GR, Maskatia S, Mathew R. 2020. COVID-19 and Kawasaki Disease: Novel Virus and Novel Case. Hospital pediatrics 10:537–540. DOI: 10.1542/hpeds.2020-0123.

De Juan D, Pazos F, Valencia A. 2013. Emerging methods in protein co-evolution. Nature Reviews Genetics 14:249–261. DOI: 10.1038/nrg3414.

Kanehisa M, Goto S. 2000. KEGG: Kyoto Encyclopedia of Genes and Genomes. Nucleic Acids Research 28:27–30. DOI: 10.1093/nar/28.1.27.

Kasela S, Daniloski Z, Jordan TX, Tenoever BR, Sanjana NE, Lappalainen T. 2021. Integrative approach identifies SLC6A20 and CXCR6 as putative causal genes for the COVID-19 GWAS signal in the 3p21.31 locus. medRxiv:2021.04.09.21255184. DOI: 10.1101/2021.04.09.21255184.

Katoh K, Standley DM. 2013. MAFFT multiple sequence alignment software version 7: Improvements in performance and usability. Molecular Biology and Evolution 30:772–780. DOI: 10.1093/molbev/mst010.

Khalili MA, Leisegang K, Majzoub A, Finelli R, Selvam MKP, Henkel R, Mojgan M, Agarwal A. 2020. Male fertility and the COVID-19 pandemic: Systematic review of the literature. World Journal of Men’s Health 38:1–15. DOI: 10.5534/WJMH.200134.

Kowalczyk A, Gbadamosi O, Kolor K, Sosa J, Croix CS, Gibson G, Chikina M, Aizenman E, Clark N, Kiselyov K. 2021. Evolutionary rate covariation identifies SLC30A9 (ZnT9) as a mitochondrial zinc transporter. bioRxiv:2021.04.22.440839 DOI: 10.1101/2021.04.22.440839.

Kriventseva E V., Kuznetsov D, Tegenfeldt F, Manni M, Dias R, Simão FA, Zdobnov EM. 2019. OrthoDB v10: Sampling the diversity of animal, plant, fungal, protist, bacterial and viral genomes for evolutionary and functional annotations of orthologs. Nucleic Acids Research 47:D807–D811. DOI: 10.1093/nar/gky1053.

Kriventseva E V., Tegenfeldt F, Petty TJ, Waterhouse RM, Simão FA, Pozdnyakov IA, Ioannidis P, Zdobnov EM. 2015. OrthoDB v8: Update of the hierarchical catalog of orthologs and the underlying free software. Nucleic Acids Research 43:D250–D256. DOI: 10.1093/nar/gku1220.

Kuba K, Imai Y, Ohto-Nakanishi T, Penninger JM. 2010. Trilogy of ACE2: A peptidase in the renin-angiotensin system, a SARS receptor, and a partner for amino acid transporters. Pharmacology and Therapeutics 128:119–128. DOI: 10.1016/j.pharmthera.2010.06.003.

Kumar S, Stecher G, Suleski M, Hedges SB. 2017. TimeTree: A Resource for Timelines, Timetrees, and Divergence Times. Molecular biology and evolution 34:1812–1819. DOI: 10.1093/molbev/msx116.

Lan J, Ge J, Yu J, Shan S, Zhou H, Fan S, Zhang Q, Shi X, Wang Q, Zhang L, Wang X. 2020. Structure of the SARS-CoV-2 spike receptor-binding domain bound to the ACE2 receptor. Nature 581:215–220. DOI: 10.1038/s41586-020-2180-5.

Lazzaroni MG, Piantoni S, Masneri S, Garrafa E, Martini G, Tincani A, Andreoli L, Franceschini F. 2021. Coagulation dysfunction in COVID-19: The interplay between inflammation, viral infection and the coagulation system. Blood Reviews 46:100745. DOI: 10.1016/j.blre.2020.100745.

Lee IH, Lee JW, Kong SW. 2020. A survey of genetic variants in SARS-CoV-2 interacting domains of ACE2, TMPRSS2 and TLR3/7/8 across populations. Infection, Genetics and Evolution 85:104507. DOI: 10.1016/j.meegid.2020.104507.

Leenaerts D, Loyau S, Mertens JC, Boisseau W, Michel JB, Lambeir AM, Jandrot-Perrus M, Hendriks D. 2018. Carboxypeptidase U (CPU, carboxypeptidase B2, activated thrombin-activatable fibrinolysis inhibitor) inhibition stimulates the fibrinolytic rate in different in vitro models. Journal of Thrombosis and Haemostasis 16:2057–2069. DOI: 10.1111/jth.14249.

Lei Y, Takahama Y. 2012. XCL1 and XCR1 in the immune system. Microbes and Infection 14:262–267. DOI: 10.1016/j.micinf.2011.10.003.

Leung LLK, Morser J. 2018. Carboxypeptidase B2 and carboxypeptidase N in the crosstalk between coagulation, thrombosis, inflammation, and innate immunity. Journal of Thrombosis and Haemostasis 16:1474–1486. DOI: 10.1111/jth.14199.

Levi M, Thachil J, Iba T, Levy JH. 2020. Coagulation abnormalities and thrombosis in patients with COVID-19. The Lancet Haematology 7:e438–e440. DOI: 10.1016/S2352-3026(20)30145-9.

van Lier D, Kox M, Santos K, van der Hoeven H, Pillay J, Pickkers P. 2021. Increased blood angiotensin converting enzyme 2 activity in critically ill COVID-19 patients. ERJ Open Research 7:00848–02020. DOI: 10.1183/23120541.00848-2020.

Liu D, Yang J, Feng B, Lu W, Zhao C, Li L. 2021. Mendelian randomization analysis identified genes pleiotropically associated with the risk and prognosis of COVID-19. Journal of Infection 82:126–132. DOI: 10.1016/j.jinf.2020.11.031.

López-León S, Wegman-Ostrosky T, Perelman C, Sepulveda R, Rebolledo PA, Cuapio A, Villapol S. 2021. More than 50 Long-Term Effects of COVID-19: A Systematic Review and Meta-Analysis. SSRN Electronic Journal:2021.01.27.21250617. DOI: 10.2139/ssrn.3769978.

Louwersheimer E, Cohn-Hokke PE, Pijnenburg YAL, Weiss MM, Sistermans EA, Rozemuller AJ, Hulsman M, Van Swieten JC, Van Duijn CM, Barkhof F, Koenei T, Scheltens P, Van DerFlier WM, Holstege H. 2017. Rare genetic variant in SORL1 may increase penetrance of Alzheimer’s disease in a family with several generations of APOE-ϵ4 homozygosity. Journal of Alzheimer’s Disease 56:63–74. DOI: 10.3233/JAD-160091.

Luo P, Liu Y, Qiu L, Liu X, Liu D, Li J. 2020. Tocilizumab treatment in COVID-19: A single center experience. Journal of Medical Virology 92:814–818. DOI: 10.1002/jmv.25801.

Luo S, Zhang X, Xu H. 2020. Don’t Overlook Digestive Symptoms in Patients With 2019 Novel Coronavirus Disease (COVID-19). Clinical Gastroenterology and Hepatology 18:1636–1637. DOI: 10.1016/j.cgh.2020.03.043.

Magro C, Mulvey JJ, Berlin D, Nuovo G, Salvatore S, Harp J, Baxter-Stoltzfus A, Laurence J. 2020. Complement associated microvascular injury and thrombosis in the pathogenesis of severe COVID-19 infection: A report of five cases. Translational Research 220:1–13. DOI: 10.1016/j.trsl.2020.04.007.

Mamoor S. 2020. The transcription factor ZBTB43 is differentially expressed and transcriptionally induced in models of coronavirus infection. DOI: 10.31219/osf.io/jhnfv.

Mangalmurti N, Hunter CA. 2020. Cytokine Storms: Understanding COVID-19. Immunity 53:19–25. DOI: 10.1016/j.immuni.2020.06.017.

Matsumoto T, Shiina H, Kawano H, Sato T, Kato S. 2008. Androgen receptor functions in male and female physiology. Journal of Steroid Biochemistry and Molecular Biology 109:236–241. DOI: 10.1016/j.jsbmb.2008.03.023.

McKechnie JL, Blish CA. 2020. The Innate Immune System: Fighting on the Front Lines or Fanning the Flames of COVID-19? Cell Host and Microbe 27:863–869. DOI: 10.1016/j.chom.2020.05.009.

Medcalf RL, Keragala CB, Myles PS. 2020. Fibrinolysis and COVID-19: A plasmin paradox. Journal of Thrombosis and Haemostasis 18:2118–2122. DOI: 10.1111/jth.14960.

Medina-Enríquez MM, Lopez-León S, Carlos-Escalante JA, Aponte-Torres Z, Cuapio A, Wegman-Ostrosky T. 2020. ACE2: the molecular doorway to SARS-CoV-2. Cell and Bioscience 10:1–17. DOI: 10.1186/s13578-020-00519-8.

Minh BQ, Schmidt HA, Chernomor O, Schrempf D, Woodhams MD, Von Haeseler A, Lanfear R, Teeling E. 2020. IQ-TREE 2: New Models and Efficient Methods for Phylogenetic Inference in the Genomic Era. Molecular Biology and Evolution 37:1530–1534. DOI: 10.1093/molbev/msaa015.

Misof B, Liu S, Meusemann K, Peters RS, Donath A, Mayer C, Frandsen PB, Ware J, Flouri T, Beutel RG, Niehuis O, Petersen M, Izquierdo-Carrasco F, Wappler T, Rust J, Aberer AJ, Aspöck U, Aspöck H, Bartel D, Blanke A, Berger S, Böhm A, Buckley TR, Calcott B, Chen J, Friedrich F, Fukui M, Fujita M, Greve C, Grobe P, Gu S, Huang Y, Jermiin LS, Kawahara AY, Krogmann L, Kubiak M, Lanfear R, Letsch H, Li Y, Li Z, Li J, Lu H, Machida R, Mashimo Y, Kapli P, McKenna DD, Meng G, Nakagaki Y, Navarrete-Heredia JL, Ott M, Ou Y, Pass G, Podsiadlowski L, Pohl H, Von Reumont BM, Schütte K, Sekiya K, Shimizu S, Slipinski A, Stamatakis A, Song W, Su X, Szucsich NU, Tan M, Tan X, Tang M, Tang J, Timelthaler G, Tomizuka S, Trautwein M, Tong X, Uchifune T, Walzl MG, Wiegmann BM, Wilbrandt J, Wipfler B, Wong TKF, Wu Q, Wu G, Xie Y, Yang S, Yang Q, Yeates DK, Yoshizawa K, Zhang Q, Zhang R, Zhang W, Zhang Y, Zhao J, Zhou C, Zhou L, Ziesmann T, Zou S, Li Y, Xu X, Zhang Y, Yang H, Wang J, Wang J, Kjer KM, Zhou X. 2014. Phylogenomics resolves the timing and pattern of insect evolution. Science 346:763–767. DOI: 10.1126/science.1257570.

Moos M, Tacke R, Scherer H, Teplow D, Früh K, Schachner M. 1988. Neural adhesion molecule L1 as a member of the immunoglobulin superfamily with binding domains similar to fibronectin. Nature 334:701–703. DOI: 10.1038/334701a0.

Morand A, Urbina D, Fabre A. 2020. COVID-19 and Kawasaki like disease : the known-known, the unknown-known and the unknown-unknown. Preprints:2020050160. DOI: 10.20944/PREPRINTS202005.0160.V1.

Morser J, Shao Z, Nishimura T, Zhou Q, Zhao L, Higgins J, Leung LLK. 2018. Carboxypeptidase B2 and N play different roles in regulation of activated complements C3a and C5a in mice. Journal of Thrombosis and Haemostasis 16:991–1002. DOI: 10.1111/jth.13964.

Mosesson MW. 2005. Fibrinogen and fibrin structure and functions. In: Journal of Thrombosis and Haemostasis. John Wiley & Sons, Ltd, 1894–1904. DOI: 10.1111/j.1538-7836.2005.01365.x.

Myllyharju J, Kivirikko KI. 2001. Collagens and collagen-related diseases. Annals of Medicine 33:7–21. DOI: 10.3109/07853890109002055.

Navarro Gonzalez J, Zweig AS, Speir ML, Schmelter D, Rosenbloom KR, Raney BJ, Powell CC, Nassar LR, Maulding ND, Lee CM, Lee BT, Hinrichs AS, Fyfe AC, Fernandes JD, Diekhans M, Clawson H, Casper J, Benet-Pagès A, Barber GP, Haussler D, Kuhn RM, Haeussler M, Kent WJ. 2021. The UCSC genome browser database: 2021 update. Nucleic Acids Research 49:D1046–D1057. DOI: 10.1093/nar/gkaa1070.

Niazkar HR, Zibaee B, Nasimi A, Bahri N. 2020. The neurological manifestations of COVID-19: a review article. Neurological Sciences 41:1667–1671. DOI: 10.1007/s10072-020-04486-3.

Omarova F, Uitte De Willige S, Ariëns RAS, Rosing J, Bertina RM, Castoldi E. 2013. Inhibition of thrombin-mediated factor V activation contributes to the anticoagulant activity of fibrinogen γ′. Journal of Thrombosis and Haemostasis 11:1669–1678. DOI: 10.1111/jth.12354.

Pagadala NS, Syed K, Tuszynski J. 2017. Software for molecular docking: a review. Biophysical Reviews 9:91–102. DOI: 10.1007/s12551-016-0247-1.

Pairo-Castineira E, Clohisey S, Klaric L, Bretherick AD, Rawlik K, Pasko D, Walker S, Parkinson N, Fourman MH, Russell CD, Furniss J, Richmond A, Gountouna E, Wrobel N, Harrison D, Wang B, Wu Y, Meynert A, Griffiths F, Oosthuyzen W, Kousathanas A, Moutsianas L, Yang Z, Zhai R, Zheng C, Grimes G, Beale R, Millar J, Shih B, Keating S, Zechner M, Haley C, Porteous DJ, Hayward C, Yang J, Knight J, Summers C, Shankar-Hari M, Klenerman P, Turtle L, Ho A, Moore SC, Hinds C, Horby P, Nichol A, Maslove D, Ling L, McAuley D, Montgomery H, Walsh T, Pereira AC, Renieri A, Shen X, Ponting CP, Fawkes A, Tenesa A, Caulfield M, Scott R, Rowan K, Murphy L, Openshaw PJM, Semple MG, Law A, Vitart V, Wilson JF, Baillie JK. 2021. Genetic mechanisms of critical illness in COVID-19. Nature 591:92–98. DOI: 10.1038/s41586-020-03065-y.

Pellegrini M, Marcotte EM, Thompson MJ, Eisenberg D, Yeates TO. 1999. Assigning protein functions by comparative genome analysis: Protein phylogenetic profiles. Proceedings of the National Academy of Sciences of the United States of America 96:4285–4288. DOI: 10.1073/pnas.96.8.4285.

Peng X, Xu E, Liang W, Pei X, Chen D, Zheng D, Zhang Y, Zheng C, Wang P, She S, Zhang Y, Ma J, Mo X, Zhang Y, Ma D, Wang Y. 2016. Identification of FAM3D as a newendogenous chemotaxis agonist for the formyl peptide receptors. Journal of Cell Science 129:1831–1842. DOI: 10.1242/jcs.183053.

Priedigkeit N, Wolfe N, Clark NL. 2015. Evolutionary Signatures amongst Disease Genes Permit Novel Methods for Gene Prioritization and Construction of Informative Gene-Based Networks. PLoS Genetics 11:1–17. DOI: 10.1371/journal.pgen.1004967.

Rabb H. 2020. Kidney diseases in the time of COVID-19: Major challenges to patient care. Journal of Clinical Investigation 130:2749–2751. DOI: 10.1172/JCI138871.

Rao VS, Srinivas K, Sujini GN, Kumar GNS. 2014. Protein-Protein Interaction Detection: Methods and Analysis. International Journal of Proteomics 2014:1–12. DOI: 10.1155/2014/147648.

Raza Q, Choi JY, Li Y, O’Dowd RM, Watkins SC, Chikina M, Hong Y, Clark NL, Kwiatkowski A V. 2019. Evolutionary rate covariation analysis of E-cadherin identifies Raskol as a regulator of cell adhesion and actin dynamics in Drosophila. PLoS Genetics 15:e1007720. DOI: 10.1371/journal.pgen.1007720.

Reilly DF, Westgate EJ, FitzGerald GA. 2007. Peripheral circadian clocks in the vasculature. Arteriosclerosis, Thrombosis, and Vascular Biology 27:1694–1705. DOI: 10.1161/ATVBAHA.107.144923.

Rizvanov AA, Kiyasov AP, Gaziziov IM, Yilmaz TS, Kaligin MS, Andreeva DI, Shafigullina AK, Guseva DS, Kiselev SL, Matin K, Palotás A, Islamov RR. 2008. Human umbilical cord blood cells transfected with VEGF and L1CAM do not differentiate into neurons but transform into vascular endothelial cells and secrete neuro-trophic factors to support neuro-genesis-a novel approach in stem cell therapy. Neurochemistry International 53:389–394. DOI: 10.1016/j.neuint.2008.09.011.

Sallard E, Lescure FX, Yazdanpanah Y, Mentre F, Peiffer-Smadja N. 2020. Type 1 interferons as a potential treatment against COVID-19. Antiviral Research 178:104791. DOI: 10.1016/j.antiviral.2020.104791.

Salman AA, Waheed MH, Ali-Abdulsahib AA, Atwan ZW. 2021. Low type i interferon response in covid-19 patients: Interferon response may be a potential treatment for covid-19. Biomedical Reports 14:1–5. DOI: 10.3892/br.2021.1419.

Samatov TR, Wicklein D, Tonevitsky AG. 2016. L1CAM: Cell adhesion and more. Progress in Histochemistry and Cytochemistry 51:25–32. DOI: 10.1016/j.proghi.2016.05.001.

Samavati L, Uhal BD. 2020. ACE2, Much More Than Just a Receptor for SARS-COV-2. Frontiers in Cellular and Infection Microbiology 10:317. DOI: 10.3389/fcimb.2020.00317.

Samuel RM, Majd H, Richter MN, Ghazizadeh Z, Zekavat SM, Navickas A, Ramirez JT, Asgharian H, Simoneau CR, Bonser LR, Koh KD, Garcia-Knight M, Tassetto M, Sunshine S, Farahvashi S, Kalantari A, Liu W, Andino R, Zhao H, Natarajan P, Erle DJ, Ott M, Goodarzi H, Fattahi F. 2020. Androgen Signaling Regulates SARS-CoV-2 Receptor Levels and Is Associated with Severe COVID-19 Symptoms in Men. Cell Stem Cell 27:876–889.e12. DOI: 10.1016/j.stem.2020.11.009.

Sánchez-Martín P, Komatsu M. 2020. Heparan sulfate and clusterin: Cleaning squad for extracellular protein degradation. Journal of Cell Biology 219. DOI: 10.1083/JCB.202001159.

Seabold S, Perktold J. 2010. Statsmodels: Econometric and Statistical Modeling with Python. In: Proceedings of the 9th Python in Science Conference. 92–96. DOI: 10.25080/majora-92bf1922-011.

Seitz R, Schramm W. 2020. DIC in COVID-19: Implications for prognosis and treatment? Journal of Thrombosis and Haemostasis 18:1798–1799. DOI: 10.1111/jth.14878.

Severe Covid-19 GWAS Group. 2020. Genomewide Association Study of Severe Covid-19 with Respiratory Failure. New England Journal of Medicine 383:1522–1534. DOI: 10.1056/nejmoa2020283.

Siddiqi HK, Libby P, Ridker PM. 2021. COVID-19 – A vascular disease. Trends in Cardiovascular Medicine 31:1–5. DOI: 10.1016/j.tcm.2020.10.005.

Singh MK, Mobeen A, Chandra A, Joshi S, Ramachandran S. 2021. A meta-analysis of comorbidities in COVID-19: Which diseases increase the susceptibility of SARS-CoV-2 infection? Computers in Biology and Medicine 130:104219. DOI: 10.1016/j.compbiomed.2021.104219.

Sriram K, Insel PA. 2020. A hypothesis for pathobiology and treatment of COVID-19: The centrality of ACE1/ACE2 imbalance. British Journal of Pharmacology 177:4825–4844. DOI: 10.1111/bph.15082.

Stelzer G, Rosen N, Plaschkes I, Zimmerman S, Twik M, Fishilevich S, Iny Stein T, Nudel R, Lieder I, Mazor Y, Kaplan S, Dahary D, Warshawsky D, Guan-Golan Y, Kohn A, Rappaport N, Safran M, Lancet D. 2016. The GeneCards suite: From gene data mining to disease genome sequence analyses. Current Protocols in Bioinformatics 2016:1.30.1–1.30.33. DOI: 10.1002/cpbi.5.

Tabach Y, Golan T, Hernández-Hernández A, Messer AR, Fukuda T, Kouznetsova A, Liu JG, Lilienthal I, Levy C, Ruvkun G. 2013. Human disease locus discovery and mapping to molecular pathways through phylogenetic profiling. Molecular Systems Biology 9:692. DOI: 10.1038/msb.2013.50.

Tafuri A, Shahinian A, Bladt F, Yoshinaga SK, Jordana M, Wakeham A, Boucher LM, Bouchard D, Chan VSF, Duncan G, Odermatt B, Ho A, Itie A, Horan T, Whoriskey JS, Pawson T, Penninger JM, Ohashi PS, Mak TW. 2001. ICOS is essential for effective T-helper-cell responses. Nature 409:105–109. DOI: 10.1038/35051113.

Takata N, Ishii K aki, Takayama H, Nagashimada M, Kamoshita K, Tanaka T, Kikuchi A, Takeshita Y, Matsumoto Y, Ota T, Yamamoto Y, Yamagoe S, Seki A, Sakai Y, Kaneko S, Takamura T. 2021. LECT2 as a hepatokine links liver steatosis to inflammation via activating tissue macrophages in NASH. Scientific Reports 11. DOI: 10.1038/s41598-020-80689-0.

Taquet M, Geddes JR, Husain M, Luciano S, Harrison PJ. 2021. 6-month neurological and psychiatric outcomes in 236 379 survivors of COVID-19: a retrospective cohort study using electronic health records. The Lancet Psychiatry 8:416–427. DOI: 10.1016/s2215-0366(21)00084-5.

Terpos E, Ntanasis-Stathopoulos I, Elalamy I, Kastritis E, Sergentanis TN, Politou M, Psaltopoulou T, Gerotziafas G, Dimopoulos MA. 2020. Hematological findings and complications of COVID-19. American Journal of Hematology 95:834–847. DOI: 10.1002/ajh.25829.

Thomas C, Moraga I, Levin D, Krutzik PO, Podoplelova Y, Trejo A, Lee C, Yarden G, Vleck SE, Glenn JS, Nolan GP, Piehler J, Schreiber G, Garcia KC. 2011. Structural linkage between ligand discrimination and receptor activation by Type i interferons. Cell 146:621–632. DOI: 10.1016/j.cell.2011.06.048.

Torra R, Tazón-Vega B, Ars E, Ballarín J. 2004. Collagen type IV (α3-α4) nephropathy: From isolated haematuria to renal failure. Nephrology Dialysis Transplantation 19:2429–2432. DOI: 10.1093/ndt/gfh435.

Uhlén M, Fagerberg L, Hallström BM, Lindskog C, Oksvold P, Mardinoglu A, Sivertsson Å, Kampf C, Sjöstedt E, Asplund A, Olsson IM, Edlund K, Lundberg E, Navani S, Szigyarto CAK, Odeberg J, Djureinovic D, Takanen JO, Hober S, Alm T, Edqvist PH, Berling H, Tegel H, Mulder J, Rockberg J, Nilsson P, Schwenk JM, Hamsten M, Von Feilitzen K, Forsberg M, Persson L, Johansson F, Zwahlen M, Von Heijne G, Nielsen J, Pontén F. 2015. Tissue-based map of the human proteome. Science 347:1260419–1260419. DOI: 10.1126/science.1260419.

Verdecchia P, Cavallini C, Spanevello A, Angeli F. 2020. The pivotal link between ACE2 deficiency and SARS-CoV-2 infection. European Journal of Internal Medicine 76:14–20. DOI: 10.1016/j.ejim.2020.04.037.

Virtanen P, Gommers R, Oliphant TE, Haberland M, Reddy T, Cournapeau D, Burovski E, Peterson P, Weckesser W, Bright J, van der Walt SJ, Brett M, Wilson J, Millman KJ, Mayorov N, Nelson ARJ, Jones E, Kern R, Larson E, Carey CJ, Polat İ, Feng Y, Moore EW, VanderPlas J, Laxalde D, Perktold J, Cimrman R, Henriksen I, Quintero EA, Harris CR, Archibald AM, Ribeiro AH, Pedregosa F, van Mulbregt P, Vijaykumar A, Bardelli A Pietro, Rothberg A, Hilboll A, Kloeckner A, Scopatz A, Lee A, Rokem A, Woods CN, Fulton C, Masson C, Häggström C, Fitzgerald C, Nicholson DA, Hagen DR, Pasechnik D V., Olivetti E, Martin E, Wieser E, Silva F, Lenders F, Wilhelm F, Young G, Price GA, Ingold GL, Allen GE, Lee GR, Audren H, Probst I, Dietrich JP, Silterra J, Webber JT, Slavič J, Nothman J, Buchner J, Kulick J, Schönberger JL, de Miranda Cardoso JV, Reimer J, Harrington J, Rodríguez JLC, Nunez-Iglesias J, Kuczynski J, Tritz K, Thoma M, Newville M, Kümmerer M, Bolingbroke M, Tartre M, Pak M, Smith NJ, Nowaczyk N, Shebanov N, Pavlyk O, Brodtkorb PA, Lee P, McGibbon RT, Feldbauer R, Lewis S, Tygier S, Sievert S, Vigna S, Peterson S, More S, Pudlik T, Oshima T, Pingel TJ, Robitaille TP, Spura T, Jones TR, Cera T, Leslie T, Zito T, Krauss T, Upadhyay U, Halchenko YO, Vázquez-Baeza Y. 2020. SciPy 1.0: fundamental algorithms for scientific computing in Python. Nature Methods 17:261–272. DOI: 10.1038/s41592-019-0686-2.

Viveiros A, Rasmuson J, Vu J, Mulvagh SL, Yip CYY, Norris CM, Oudit GY. 2021. Sex differences in COVID-19: Candidate pathways, genetics of ACE2, and sex hormones. American Journal of Physiology - Heart and Circulatory Physiology 320:H296–H304. DOI: 10.1152/AJPHEART.00755.2020.

Wiradjaja F, DiTommaso T, Smyth I. 2010. Basement membranes in development and disease. Birth Defects Research Part C - Embryo Today: Reviews 90:8–31. DOI: 10.1002/bdrc.20172.

Wolfe NW, Clark NL. 2015. ERC analysis: Web-based inference of gene function via evolutionary rate covariation. Bioinformatics 31:3835–3837. DOI: 10.1093/bioinformatics/btv454.

Wright FL, Vogler TO, Moore EE, Moore HB, Wohlauer M V., Urban S, Nydam TL, Moore PK, McIntyre RC. 2020. Fibrinolysis Shutdown Correlation with Thromboembolic Events in Severe COVID-19 Infection. Journal of the American College of Surgeons 231:193–203.e1. DOI: 10.1016/j.jamcollsurg.2020.05.007.

Wu S, Miao L, Zhou Q, Gao C, Liu J, Zhan Q, Guo B, Li F, Wang Y, Xu H, Yan H, Wu R, Zhang S, Zheng J, Yang J, Wang S, Yu W, Niu H, Li F, Yang L, Huang J, Lu X, Chen J, Tong Y, Ma L, Zhou Y, Guo Q. 2020. Suppression of Androgen Receptor (AR)-ACE2/TMPRSS2 Axis by AR Antagonists May Be Therapeutically Beneficial for Male COVID-19 Patients. SSRN Electronic Journal. DOI: 10.2139/ssrn.3580526.

Wyatt AR, Wilson MR. 2010. Identification of human plasma proteins as major clients for the extracellular chaperone clusterin. Journal of Biological Chemistry 285:3532–3539. DOI: 10.1074/jbc.M109.079566.

Xie Z, Bailey A, Kuleshov M V., Clarke DJB, Evangelista JE, Jenkins SL, Lachmann A, Wojciechowicz ML, Kropiwnicki E, Jagodnik KM, Jeon M, Ma’ayan A. 2021. Gene Set Knowledge Discovery with Enrichr. Current Protocols 1:e90. DOI: 10.1002/cpz1.90.

Yamada T, Nakao K, Itoh H, Morii N, Shiono S, Sakamoto M, Sugawara A, Saito Y, Mukoyama M, Arai H, Eigyo M, Matsushita A, Imura H. 1988. Inhibitory action of leumorphin on vasopressin secretion in conscious rats. Endocrinology 122:985–990. DOI: 10.1210/endo-122-3-985.

Yamagoe S, Yamakawa Y, Matsuo Y, Minowada J, Mizuno S, Suzuki K. 1996. Purification and primary amino acid sequence of a novel neutrophil chemotactic factor LECT2. Immunology Letters 52:9–13. DOI: 10.1016/0165-2478(96)02572-2.

Yan Z, Ye G, Werren JH. 2019. Evolutionary Rate Correlation between Mitochondrial-Encoded and Mitochondria-Associated Nuclear-Encoded Proteins in Insects. Molecular Biology and Evolution 36:1022–1036. DOI: 10.1093/molbev/msz036.

Yeung ML, Teng JLL, Jia L, Zhang C, Huang C, Cai JP, Zhou R, Chan KH, Zhao H, Zhu L, Siu KL, Fung SY, Yung S, Chan TM, To KKW, Chan JFW, Cai Z, Lau SKP, Chen Z, Jin DY, Woo PCY, Yuen KY. 2021. Soluble ACE2-mediated cell entry of SARS-CoV-2 via interaction with proteins related to the renin-angiotensin system. Cell 184:2212–2228.e12. DOI: 10.1016/j.cell.2021.02.053.

Zhang Q, Liu Z, Moncada-Velez M, Chen J, Ogishi M, Bigio B, Yang R, Arias AA, Zhou Q, Han JE, Ugurbil AC, Zhang P, Rapaport F, Li J, Spaan AN, Boisson B, Boisson-Dupuis S, Bustamante J, Puel A, Ciancanelli MJ, Zhang SY, Béziat V, Jouanguy E, Abel L, Cobat A, Casanova JL, Bastard P, Korol C, Rosain J, Philippot Q, Chbihi M, Lorenzo L, Bizien L, Neehus AL, Kerner G, Seeleuthner Y, Manry J, Le Voyer T, Boisson B, Boisson-Dupuis S, Bustamante J, Puel A, Zhang SY, Béziat V, Jouanguy E, Abel L, Cobat A, Casanova JL, Bastard P, Rosain J, Philippot Q, Chbihi M, Lorenzo L, Bizien L, Neehus AL, Kerner G, Seeleuthner Y, Manry J, Le Voyer T, Boisson B, Boisson-Dupuis S, Bustamante J, Puel A, Zhang SY, Béziat V, Jouanguy E, Abel L, Cobat A, Casanova JL, Le Pen J, Schneider WM, Razooky BS, Hoffmann HH, Michailidis E, Rice CM, Sabli IKD, Hodeib S, Sancho-Shimizu V, Bilguvar K, Ye J, Maniatis T, Bolze A, Arias AA, Arias AA, Zhang Y, Notarangelo LD, Su HC, Zhang Y, Notarangelo LD, Su HC, Onodi F, Korniotis S, Karpf L, Soumelis V, Bonnet-Madin L, Amara A, Dorgham K, Gorochov G, Smith N, Duffy D, Moens L, Meyts I, Meade P, García-Sastre A, Krammer F, Corneau A, Masson C, Schmitt Y, Schlüter A, Pujol A, Khan T, Marr N, Fellay J, Fellay J, Fellay J, Roussel L, Vinh DC, Shahrooei M, Shahrooei M, Alosaimi MF, Alsohime F, Hasanato R, Mansouri D, Mansouri D, Mansouri D, Al-Saud H, Almourfi F, Al-Mulla F, Al-Muhsen SZ, Al Turki S, Al Turki S, van de Beek D, Biondi A, Bettini LR, D’Angio M, Bonfanti P, Imberti L, Sottini A, Paghera S, Quiros-Roldan E, Rossi C, Oler AJ, Tompkins MF, Alba C, Dalgard CL, Vandernoot I, Smits G, Goffard JC, Migeotte I, Haerynck F, Soler-Palacin P, Martin-Nalda A, Colobran R, Morange PE, Keles S, Çölkesen F, Ozcelik T, Yasar KK, Senoglu S, Karabela ŞN, Rodríguez-Gallego C, Rodríguez-Gallego C, Novelli G, Hraiech S, Tandjaoui-Lambiotte Y, Tandjaoui-Lambiotte Y, Duval X, Laouénan C, Duval X, Laouénan C, Laouénan C, Snow AL, Dalgard CL, Milner JD, Mogensen TH, Mogensen TH, Marr N, Spaan AN, Bustamante J, Ciancanelli MJ, Meyts I, Maniatis T, Soumelis V, Nussenzweig M, Nussenzweig M, Casanova JL, García-Sastre A, García-Sastre A, García-Sastre A, Lifton RP, Lifton RP, Lifton RP, Casanova JL, Foti G, Bellani G, Citerio G, Contro E, Pesci A, Valsecchi MG, Cazzaniga M, Abad J, Blanco I, Rodrigo C, Aguilera-Albesa S, Akcan OM, Darazam IA, Aldave JC, Ramos MA, Nadji SA, Alkan G, Allardet-Servent J, Allende LM, Alsina L, Alyanakian MA, Amador-Borrero B, Mouly S, Sene D, Amoura Z, Mathian A, Antolí A, Blanch GR, Riera JS, Moreno XS, Arslan S, Assant S, Auguet T, Azot A, Bajolle F, Bustamante J, Lévy R, Oualha M, Baldolli A, Ballester M, Feldman HB, Barrou B, Beurton A, Bilbao A, Blanchard-Rohner G, Blandinières A, Rivet N, Blazquez-Gamero D, Bloomfield M, Bolivar-Prados M, Clavé P, Borie R, Bosteels C, Lambrecht BN, van Braeckel E, Bousfiha AA, Bouvattier C, Vincent A, Boyarchuk O, Bueno MRP, Castro M V., Matos LRB, Zatz M, Agra JJC, Calimli S, Capra R, Carrabba M, Fabio G, Casasnovas C, Vélez-Santamaria V, Caseris M, Falck A, Poncelet G, Castelle M, Castelli F, de Vera MC, Catherinot E, Chalumeau M, Toubiana J, Charbit B, Li Z, Pellegrini S, Cheng MP, Clotet B, Codina A, Colkesen F, Çölkesen F, Colobran R, Comarmond C, Dalmau D, Dalmau D, Darley DR, Dauby N, Dauger S, Le Bourgeois F, Levy M, de Pontual L, Dehban A, Delplancq G, Demoule A, Diehl JL, Dobbelaere S, Durand S, Mircher C, Rebillat AS, Vilaire ME, Eldars W, Elgamal M, Elnagdy MH, Emiroglu M, Erdeniz EH, Aytekin SE, Euvrard R, Evcen R, Faivre L, Fartoukh M, Philippot Q, Faure M, Arquero MF, Flores C, Flores C, Flores C, Flores C, Francois B, Fumadó V, Fumadó V, Fumadó V, Fusco F, Ursini MV, Solis BG, de Diego RP, van Den Rym AM, Gaussem P, Gil-Herrera J, Gilardin L, Alarcon MG, Girona-Alarcón M, Goffard JC, Gok F, Yosunkaya A, González-Montelongo R, Íñigo-Campos A, Lorenzo-Salazar JM, Muñoz-Barrera A, Guerder A, Gul Y, Guner SN, Gut M, Hadjadj J, Haerynck F, Halwani R, Hammarström L, Hatipoglu N, Hernandez-Brito E, Heijmans C, Holanda-Peña MS, Horcajada JP, Hoste L, Hoste E, Hraiech S, Humbert L, Mordacq C, Thumerelle C, Vuotto F, Iglesias AD, Jamme M, Arranz MJ, Jordan I, Jorens P, Kanat F, Kapakli H, Kara I, Karbuz A, Yasar KK, Senoglu S, Keles S, Demirkol YK, Klocperk A, Król ZJ, Kuentz P, Kwan YWM, Lagier JC, Lau YL, Leung D, Leo YS, Young BE, Lopez RL, Levin M, Linglart A, Loeys B, Louapre C, Lubetzki C, Luyt CE, Lye DC, Mansouri D, Marjani M, Pereira JM, Martin A, Soler-Palacín P, Pueyo DM, Martinez-Picado J, Marzana I, Matthews G V., Mayaux J, Parizot C, Quentric P, Mège JL, Raoult D, Melki I, Meritet JF, Metin O, Meyts I, Mezidi M, Migeotte I, Taccone F, Millereux M, Mirault T, Mirsaeidi M, Melián AM, Martinez AM, Morange P, Morelle G, Naesens L, Nafati C, Neves JF, Ng LFP, Medina YN, Cuadros EN, Gonzalo Ocejo-Vinyals J, Orbak Z, Özçelik T, Pan-Hammarström Q, Pascreau T, Paz-Artal E, Philippe A, Planas-Serra L, Schluter A, Ploin D, Viel S, Poissy J, Pouletty M, Reisli I, Ricart P, Richard JC, Rivière JG, Rodriguez-Gallego C, Rodriguez-Gallego C, Rodríguez-Palmero A, Romero CS, Rothenbuhler A, Rozenberg F, del Prado MYR, Sanchez O, Sánchez-Ramón S, Schmidt M, Schweitzer CE, Scolari F, Sediva A, Seijo LM, Seppänen MRJ, Ilovich AS, Slabbynck H, Smadja DM, Sobh A, Solé-Violán J, Soler C, Stepanovskiy Y, Stoclin A, Tandjaoui-Lambiotte Y, Taupin JL, Tavernier SJ, Terrier B, Tomasoni G, Alvarez JT, Trouillet-Assant S, Troya J, Tucci A, Uzunhan Y, Vabres P, Valencia-Ramos J, van de Velde S, van Praet J, Vandernoot I, Vatansev H, Vilain C, Voiriot G, Yucel F, Zannad F, Belot A, Bole-Feysot C, Lyonnet S, Masson C, Nitschke P, Pouliet A, Schmitt Y, Tores F, Zarhrate M, Shahrooei M, Abel L, Andrejak C, Angoulvant F, Bachelet D, Bhavsar K, Bouadma L, Chair A, Couffignal C, Silveira C Da, Debray MP, Duval X, Eloy P, Esposito-Farese M, Ettalhaoui N, Gault N, Ghosn J, Gorenne I, Hoffmann I, Kafif O, Kali S, Khalil A, Laouénan C, Laribi S, Le M, Le Hingrat Q, Lescure FX, Lucet JC, Mentré F, Mullaert J, Peiffer-Smadja N, Peytavin G, Roy C, Schneider M, Mohammed NS, Tagherset L, Tardivon C, Tellier MC, Timsit JF, Trioux T, Tubiana S, Basmaci R, Behillil S, Beluze M, Benkerrou D, Dorival C, Meziane A, Téoulé F, Bompart F, Bouscambert M, Gaymard A, Lina B, Rosa-Calatrava M, Terrier O, Caralp M, Cervantes-Gonzalez M, D’Ortenzio E, Puéchal O, Semaille C, Coelho A, Diouf A, Hoctin A, Mambert M, Couffin-Cadiergues S, Deplanque D, Descamps D, Visseaux B, Desvallées M, Khan C, Diallo A, Mercier N, Paul C, Petrov-Sanchez V, Dubos F, Enouf VVE, Mouquet H, Esperou H, Jaafoura S, Papadopoulos A, Etienne M, Gigante T, Rossignol B, Guedj J, Le Nagard H, Lingas G, Neant N, Kaguelidou F, Lévy Y, Wiedemann A, Lévy Y, Wiedemann A, Levy-Marchal C, Malvy D, Noret M, Pages J, Picone O, Rossignol P, Tual C, Veislinger A, van der Werf S, Vanel N, Yazdanpanah Y, Alavoine L, Costa Y, Duval X, Ecobichon JL, Frezouls W, Ilic-Habensus E, Leclercq A, Lehacaut J, Letrou S, Mandic M, Nouroudine M, Quintin C, Rexach J, Tubiana S, Vignali V, Amat KKA, Behillil S, Enouf V, van der Werf S, Bielicki J, Bruijning P, Burdet C, Burdet C, Caumes E, Charpentier C, Damond F, Descamps D, Le Hingrat Q, Visseaux B, Coignard B, Couffin-Cadiergues S, Delmas C, Espérou H, Roufai L, Dechanet A, Houhou N, Kafif O, Kikoine J, Manchon P, Piquard V, Postolache A, Terzian Z, Lebeaux D, Lina B, Lucet JC, Malvy D, Meghadecha M, Motiejunaite J, Thy M, van Agtmael M, Bomers M, Chouchane O, Geerlings S, Goorhuis B, Grobusch MP, Harris V, Hermans SM, Hovius JW, Nellen J, Peters E, van der Poll T, Prins JM, Reijnders T, Schinkel M, Sigaloff K, Stijnis CS, van der Valk M, van Vugt M, Joost Wiersinga W, Algera AG, van Baarle F, Bos L, Botta M, de Bruin S, Bulle E, Elbers P, Fleuren L, Girbes A, Hagens L, Heunks L, Horn J, van Mourik N, Paulus F, Raasveld J, Schultz MJ, Smit M, Stilma W, Thoral P, Tsonas A, de Vries H, Bax D, Cloherty A, Beudel M, Brouwer MC, Koning R, van de Beek D, Bogaard HJ, de Brabander J, de Bree G, Bugiani M, Geerts B, Hollmann MW, Preckel B, Veelo D, Geijtenbeek T, Hafkamp F, Hamann J, Hemke R, de Jong MD, Schuurman A, Teunissen C, Vlaar APJ, Wouters D, Zwinderman AH, Abel L, Aiuti A, Muhsen S Al, Al-Mulla F, Anderson MS, Arias AA, Feldman HB, Bogunovic D, Itan Y, Bolze A, Cirulli E, Barrett KS, Washington N, Bondarenko A, Bousfiha AA, Brodin P, Bryceson Y, Bustamante CD, Butte M, Casari G, Chakravorty S, Christodoulou J, Le Mestre S, Condino-Neto A, Cooper MA, Dalgard CL, David A, DeRisi JL, DeRisi JL, Desai M, Drolet BA, Espinosa S, Fellay J, Flores C, Franco JL, Gregersen PK, Haerynck F, Hagin D, Halwani R, Heath J, Henrickson SE, Hsieh E, Imai K, Karamitros T, Kisand K, Ku CL, Lau YL, Ling Y, Lucas CL, Maniatis T, Mansouri D, Marodi L, Meyts I, Milner J, Mironska K, Mogensen T, Morio T, Ng LFP, Notarangelo LD, Su HC, Novelli A, Novelli G, O’Farrelly C, Okada S, Ozcelik T, de Diego RP, Planas AM, Prando C, Pujol A, Quintana-Murci L, Renia L, Renieri A, Rodríguez-Gallego C, Sancho-Shimizu V, Sankaran V, Shahrooei M, Snow A, Soler-Palacín P, Spaan AN, Tangye S, Turvey S, Uddin F, Uddin MJ, Uddin MJ, van de Beek D, Vazquez SE, Vinh DC, von Bernuth H, Zawadzki P, Casanova JL, Jing H, Tung W, Meguro K, Shaw E, Jing H, Tung W, Shafer S, Zheng L, Zhang Z, Kubo S, Chauvin SD, Meguro K, Shaw E, Lenardo M, Luthers CR, Bauman BM, Shafer S, Zheng L, Zhang Z, Kubo S, Chauvin SD, Lenardo M, Lack J, Karlins E, Hupalo DM, Rosenberger J, Sukumar G, Wilkerson MD, Zhang X. 2020. Inborn errors of type I IFN immunity in patients with life-threatening COVID-19. Science 370. DOI: 10.1126/science.abd4570.

Zhang Z, Zeng H, Lin J, Hu Y, Yang R, Sun J, Chen R, Chen H. 2018. Circulating LECT2 levels in newly diagnosed type 2 diabetes mellitus and their association with metabolic parameters. Medicine (United States) 97. DOI: 10.1097/MD.0000000000010354.

Van Zundert GCP, Rodrigues JPGLM, Trellet M, Schmitz C, Kastritis PL, Karaca E, Melquiond ASJ, Van Dijk M, De Vries SJ, Bonvin AMJJ. 2016. The HADDOCK2.2 Web Server: User-Friendly Integrative Modeling of Biomolecular Complexes. Journal of Molecular Biology 428:720–725. DOI: 10.1016/j.jmb.2015.09.014.

