## Supplementary Information for "Evolutionary Inference Predicts Novel ACE2 Protein Interactions Relevant to COVID-19 Pathologies"

Supplementary Information is composed of three parts: (1) Large supplementary files deposited at FigShare (https://doi.org/10.6084/m9.figshare.14637450), (2) Python and R code for ERC pipelines and additional analyses deposited in GitHub (https://github.com/austinv11/ERC-Pipeline), and (3) Supplementary Text with embedded associated figures and tables.

**1. FigShare Collection: The following files are available at https://doi.org/10.6084/m9.figshare.14637450.**

**File S1**: Select proteins' 30MY ERC lists, contains multiple-test corrected p-values.
**File S2**: Pairwise ρ and unadjusted p-value 30MY ERC matrices for all proteins.
**File S3**: Enrichment results for select top ERC protein sets.
**File S4**: Zip file containing the mammalian time-scaled phylogeny and maximum likelihood protein trees in newick format.
**File S5**: Table depicting the total number of taxa present for each protein's sequence data, along with the number of taxa for which there are paralogy in the uncorrected and 30MY corrected data.
**File S6**: Branch time to terminal branch rate correlation results for the protein set.
**File S7**: Chi-squared test results for all proteins testing for whether there is an overrepresentation of rates below the regression line for short branches (<30MY).
**File S8**: Branch time vs terminal branch rate residuals to branch time correlation results for the protein set.
**File S9**: Wilcoxon matched signed-rank test significance values testing for branch adjustments following 20MY and 30MY adjustments.
**File S10**: Coefficients for the select proteins used for the linear models containing ACE2 rate rank, Btime rate rank, and taxonomic orders as independent variables.
**File S11**: 30MY-adjusted ERC comparisons within and between CORUM complex members.

**2. Code Repository:** https://github.com/austinv11/ERC-Pipeline

**3. Supplementary Text with Embedded Figures and Tables:** Below is the supplementary text with associated figures and tables

#### SUPPLEMENTARY TEXT

#### Index

A. Mammalian Data Set 4
B. ERCs on The Original Phylogeny with Short Branches 9
C. Branch Time to Protein-Rate Correlation Problem 10
D. Partial Correlation to Address BT-PR Correlation 12
E. Removing Short Branches to Remove BT-Rate Confounding Effects 13
F. Testing Whether Branch Rate Increases with Evolutionary Time 17
G. Testing for Taxonomic Order Effects 21
H. Additional ACE2 Interactor Descriptions 23
I. Supplementary Text References 32

**Supplementary Tables**

**Table S1**: Disambiguated proteins from OrthoDB groups with ancient family expansions.
**Table S2**: List of taxa considered in the mammalian phylogeny with no adjustment, 20MY adjustment, and 30MY adjustment.
**Table S3**: The top 40 ERCs for ACE2 based on the original ERC method, the BTime-adjusted partial ERC method, and the 30MY adjusted ERC method.
**Table S4**: Select Spearman's correlations for protein rates against BTime using the original rate data.
**Table S5**: Regression and ANOVA results for linear models with ACE2 and BTime as independent variables.
**Table S6**: Select Spearman's correlation test results for rate-to-BTime residual models against BTime.
**Table S7**: Select Spearman's correlation test results for rates against BTime using the original rate data, 20MY-adjusted rate data, and 30MY-adjusted rate data.
**Table S8**: Comparison of the FGA-FGB ERC using the original ERC method, the BTime-adjusted partial ERC method, and the 30MY adjusted ERC method.
**Table S9**: Comparison of the FGA-FGG ERC using the original ERC method, the BTime-adjusted partial ERC method, and the 30MY adjusted ERC method.
**Table S10**: Comparison of the FGB-FGG ERC using the original ERC method, the BTime-adjusted partial ERC method, and the 30MY adjusted ERC method.
**Table S11**: Comparison of the IFNAR1-IFNAR2 ERC using the original ERC method, the BTime-adjusted partial ERC method, and the 30MY adjusted ERC method.
**Table S12**: Comparison of the Collagen Type IV interacting pairs using the original ERC method, the BTime-adjusted partial ERC method, and the 30MY adjusted ERC method.
**Table S13**: Select Wilcoxon matched signed-rank test results testing for whether the rates of branches increased as time scales are increased.
**Table S14**: Linear regression fit statistics for select proteins using models that used ACE2 rate ranks, BTime ranks, and taxonomic order as independent variables.
**Table S15**: Select ANCOVA results for linear models that used ACE2 rate ranks, BTime ranks, and taxonomic order as independent variables.
**Table S16**: Spearman independent contrasts tests for select proteins against ACE2.

**Supplementary Figures**

**Figure S1**: Full time-scaled mammalian phylogeny.
**Figure S2**: 30MY-adjusted time-scaled mammalian phylogeny.

**Figure S3**: Scatterplots depicting the rates of ACE2 to some proteins of interest using the original rate data.
**Figure S4**: Scatterplots depicting the rates of ACE2 to some proteins of interest using the original rate data and 30MY-adjusted rate data side-by-side.
**Figure S5**: Cartoon illustrating how branches were extended to test for rate changes as time scales were increased.
**Figure S6**: Box plots depicting the rate shifts following 20MY and 30MY adjustments for select proteins.
**Figure S7**: The distribution of p-values of Wilcoxon matched signed-rank tests of all proteins comparing the rate shifts following the 20MY and 30MY adjustments.

#### A. Mammalian Data Set

As described in the methods section, the data set is primarily based on the orthologous protein groups available on OrthoDB (Kriventseva et al., 2019) based on the “mammalia” taxonomic level. We selected protein groups that are single-copy in all species with greater than 90 taxa represented. An additional 156 proteins, which did not meet the initial single copy in all taxa requirement, were added to extend the analysis in pathways of interest (e.g. coagulation cascade, sphingolipid signaling, renin-angiotensin system). Of these proteins, 47 were added due to literature suggesting an association with COVID-19, to evaluate their ERCs to ACE2, such as IFNAR2 and XCR1 (Severe Covid-19 GWAS Group, 2020; Pairo-Castineira et al., 2021; Fricke-Galindo & Falfán-Valencia, 2021). Only proteins with relatively minor paralogy issues were added by this method (Supplementary File S5). The rationale for this approach is that it would be very difficult to determine which paralog to choose for the analysis in terminal branches with multiple paralogs for a particular protein. The final set contains a total of 1,953 proteins, including ACE2.

In 23 cases (Table S1), OrthoDB orthology groups contain multiple distinct protein groups resulting from ancient gene duplications. In some cases, we examined the phylogeny of the orthology group and, where appropriate, divided and added them to our protein set. In most cases, the division was supported by protein annotation names within the orthology group, and the protein sequences were split based on reference annotations given by OrthoDB and sequence similarity. For example, coagulation factor IX (F9) and X (F10) were within the same orthology group (OrthoDB ID: 91794at40674).

| OrthoDB ID | Distinct Proteins Added |
|---|---|
| 10776at40674 | IGF1R,INSR |
| 15742at40674 | ABCC1,ABCC3,ABCC6 |
| 25854at40674 | DPP8,DPP9 |
| 32671at40674 | MAP3K5,MAP3K15 |
| 46864at40674 | LIFR,OSMR |
| 55743at40674 | PRKCI,PRKCZ |
| 66003at40674 | BMX,BTK |
| 68344at40674 | SPTLC2,SPTLC3 |
| 79978at40674 | PPP2R5D |
| 85041at40674 | TMPRSS2,TMPRSS3 |
| 91794at40674 | F9,F10 |
| 94914at40674 | PPP2R2A,PPP2R2B,PPP2R2C |
| 95740at40674 | MAPK8,MAPK10 |
| 103747at40674 | GLA,NAGA |
| 111203at40674 | MAPK12,MAPK13 |
| 114138at40674 | CERS5,CERS6 |
| 123408at40674 | DEGS1,DEGS2 |
| 123688at40674 | SGPP1,SGPP2 |
| 123726at40674 | PPP2CA,PPP2CB |
| 129864at40674 | MAPK11,MAPK14 |
| 132357at40674 | SPHK1,SPHK2 |
| 138259at40674 | CCR2,CCR5 |
| 166274at40674 | ACER1,ACER2 |

Table S1: The OrthoDB groups that were added to the dataset for which there were multiple distinct proteins reported as a single orthology group. The proteins listed on the right column were all the disambiguated proteins added to the 30MY dataset (so they had to have met our requirement of having at least 50 of the selected taxa).

A well-resolved time-scaled mammalian phylogeny available from TimeTree (Kumar et al., 2017) was used that includes the taxa that were in our orthologous protein sets. This tree contained 108 mammals (Fig. S1, Table S2) in the original uncorrected data set. Later, in order to correct a terminal branch time (BT) to protein rate correlation found for most proteins due to short branches (see below), we removed taxa from oversampled clades with short terminal branches. We found that a 30MY threshold for terminal branches eliminated the terminal branch time to protein rate for 87.5% of proteins (described in Section E), resulting in 50-60 taxa per protein (Table S2). These data were used for the ERC analysis reported in the main text.

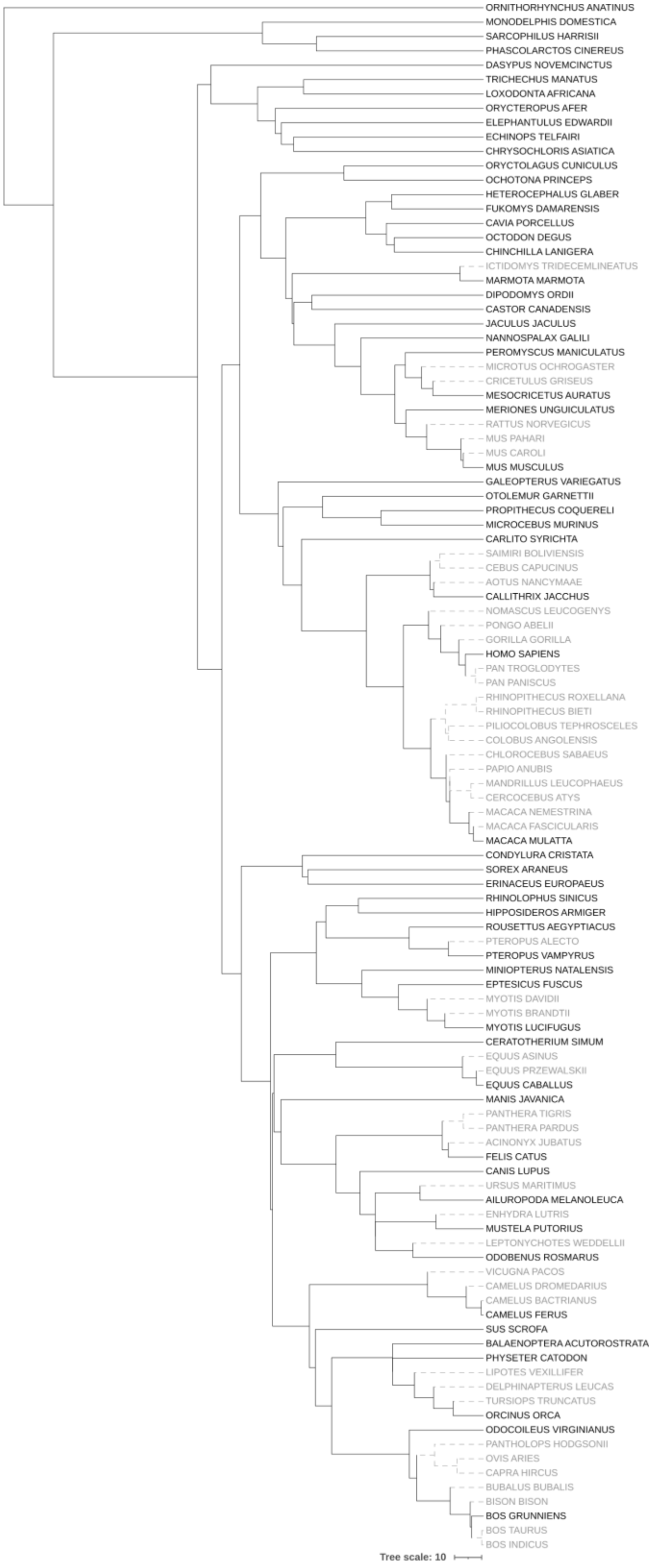


Figure S1: Full original phylogeny topology with branches scaled to time (in millions of years) based on TimeTree (Kumar et al., 2017). Branches highlighted in grey are removed following a 30MY branch length threshold correction. The tree illustration is created using iTOL (Letunic & Bork, 2021).

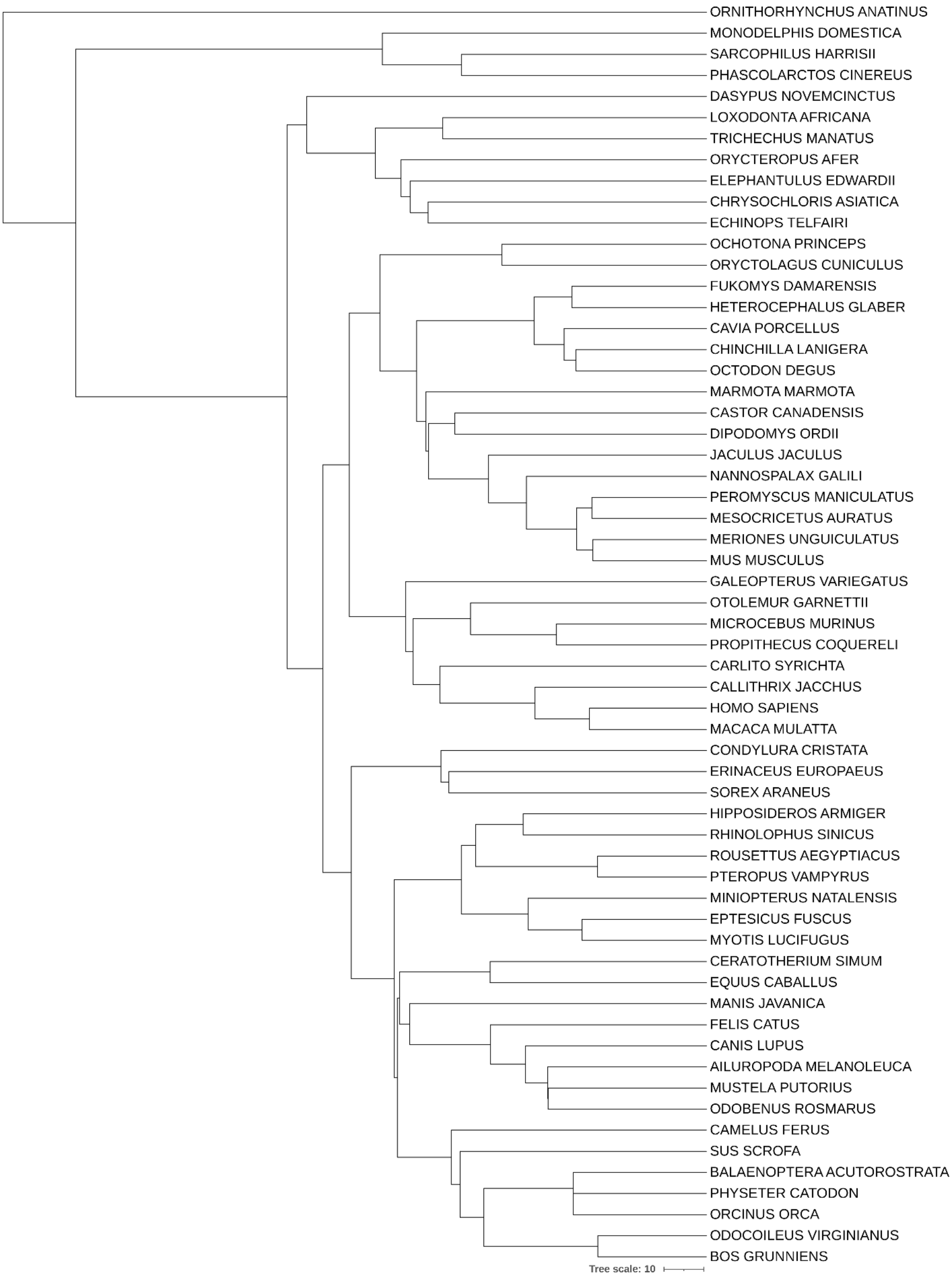


Figure S2: Time-scaled phylogeny only containing the 60 selected taxa following a 30MY threshold correction. The tree illustration is created using iTOL (Letunic & Bork, 2021).

| Full Taxa List | 20MY Taxa List | 30MY Taxa List |
|---|---|---|
| Ornithorhynchus anatinus | Ornithorhynchus anatinus | Ornithorhynchus anatinus |
| Dasypus novemcinctus | Monodelphis domestica | Monodelphis domestica |
| Orycteropus afer | Phascolarctos cinereus | Phascolarctos cinereus |
| Elephantulus edwardii | Sarcophilus harrisii | Sarcophilus harrisii |
| Echinops telfairi | Dasypus novemcinctus | Dasypus novemcinctus |
| Chrysochloris asiatica | Loxodonta africana | Loxodonta africana |
| Trichechus manatus | Trichechus manatus | Trichechus manatus |
| Loxodonta africana | Orycteropus afer | Orycteropus afer |
| Galeopterus variegatus | Elephantulus edwardii | Elephantulus edwardii |
| Carlito syrichta | Chrysochloris asiatica | Chrysochloris asiatica |
| Nomascus leucogenys | Echinops telfairi | Echinops telfairi |
| Pongo abelii | Ochotona princeps | Ochotona princeps |
| Gorilla gorilla | Oryctolagus cuniculus | Oryctolagus cuniculus |
| Homo sapiens | Fukomys damarensis | Fukomys damarensis |
| Pan troglodytes | Heterocephalus glaber | Heterocephalus glaber |
| Pan paniscus | Cavia porcellus | Cavia porcellus |
| Piliocolobus tephrosceles | Chinchilla lanigera | Chinchilla lanigera |
| Colobus angolensis | Octodon degus | Octodon degus |
| Rhinopithecus roxellana | Marmota marmota | Marmota marmota |
| Rhinopithecus bieti | Castor canadensis | Castor canadensis |
| Chlorocebus sabaeus | Dipodomys ordii | Dipodomys ordii |
| Macaca nemestrina | Jaculus jaculus | Jaculus jaculus |
| Macaca mulatta | Nannospalax galili | Nannospalax galili |
| Macaca fascicularis | Peromyscus maniculatus | Peromyscus maniculatus |
| Papio anubis | Microtus ochrogaster | Mesocricetus auratus |
| Mandrillus leucophaeus | Cricetulus griseus | Meriones unguiculatus |
| Cercocebus atys | Mesocricetus auratus | Mus musculus |
| Saimiri boliviensis | Meriones unguiculatus | Galeopterus variegatus |
| Cebus capucinus | Rattus norvegicus | Otolemur garnettii |
| Callithrix jacchus | Mus musculus | Microcebus murinus |
| Aotus nancymaae | Galeopterus variegatus | Propithecus coquereli |
| Otolemur garnettii | Otolemur garnettii | Carlito syrichta |
| Propithecus coquereli | Microcebus murinus | Callithrix jacchus |
| Microcebus murinus | Propithecus coquereli | Homo sapiens |
| Marmota marmota | Carlito syrichta | Macaca mulatta |
| Ictidomys tridecemlineatus | Aotus nancymaae | Condylura cristata |
| Dipodomys ordii | Callithrix jacchus | Erinaceus europaeus |
| Castor canadensis | Saimiri boliviensis | Sorex araneus |
| Jaculus jaculus | Nomascus leucogenys | Hipposideros armiger |
| Nannospalax galili | Homo sapiens | Rhinolophus sinicus |
| Peromyscus maniculatus | Rhinopithecus bieti | Rousettus aegyptiacus |
| Microtus ochrogaster | Macaca mulatta | Pteropus vampyrus |
| Mesocricetus auratus | Condylura cristata | Miniopterus natalensis |
| Cricetulus griseus | Erinaceus europaeus | Eptesicus fuscus |
| Meriones unguiculatus | Sorex araneus | Myotis lucifugus |
| Rattus norvegicus | Hipposideros armiger | Ceratotherium simum |
| Mus pahari | Rhinolophus sinicus | Equus caballus |
| Mus musculus | Rousettus aegyptiacus | Manis javanica |
| Mus caroli | Pteropus vampyrus | Felis catus |
| Cavia porcellus | Miniopterus natalensis | Canis lupus |
| Octodon degus | Eptesicus fuscus | Ailuropoda melanoleuca |
| Chinchilla lanigera | Myotis davidii | Mustela putorius |
| Heterocephalus glaber | Myotis lucifugus | Odobenus rosmarus |
| Fukomys damarensis | Ceratotherium simum | Camelus ferus |
| Oryctolagus cuniculus | Equus caballus | Sus scrofa |
| Ochotona princeps | Manis javanica | Balaenoptera acutorostrata |
| Miniopterus natalensis | Felis catus | Physeter catodon |
| Eptesicus fuscus | Canis lupus | Orcinus orca |
| Myotis davidii | Ailuropoda melanoleuca | Odocoileus virginianus |
| Myotis lucifugus | Ursus maritimus | Bos grunniens |
| Myotis brandtii | Enhydra lutris | |
| Rousettus aegyptiacus | Mustela putorius | |
| Pteropus vampyrus | Leptonychotes weddellii | |
| Pteropus alecto | Odobenus rosmarus | |
| Rhinolophus sinicus | Vicugna pacos | |
| Hipposideros armiger | Camelus ferus | |
| Sus scrofa | Sus scrofa | |
| Odocoileus virginianus | Balaenoptera acutorostrata | |
| Pantholops hodgsonii | Physeter catodon | |
| Ovis aries | Lipotes vexillifer | |
| Capra hircus | Delphinapterus leucas | |
| Bubalus bubalis | Orcinus orca | |
| Bison bison | Odocoileus virginianus | |
| Bos grunniens | Pantholops hodgsonii | |
| Bos taurus | Ovis aries | |
| Bos indicus | Bos grunniens | |
| Balaenoptera acutorostrata | | |
| Physeter catodon | | |
| Lipotes vexillifer | | |
| Delphinapterus leucas | | |
| Tursiops truncatus | | |
| Orcinus orca | | |
| Vicugna pacos | | |
| Camelus dromedarius | | |
| Camelus ferus | | |
| Camelus bactrianus | | |
| Ceratotherium simum | | |
| Equus asinus | | |
| Equus przewalskii | | |
| Equus caballus | | |
| Manis javanica | | |
| Canis lupus | | |
| Ursus maritimus | | |
| Ailuropoda melanoleuca | | |
| Odobenus rosmarus | | |
| Leptonychotes weddellii | | |
| Mustela putorius | | |
| Enhydra lutris | | |
| Felis catus | | |
| Acinonyx jubatus | | |
| Panthera tigris | | |
| Panthera pardus | | |
| Condylura cristata | | |
| Sorex araneus | | |
| Erinaceus europaeus | | |
| Monodelphis domestica | | |
| Sarcophilus harrisii | | |
| Phascolarctos cinereus | | |

Table S2: List of taxa that are in the original phylogeny (left column), the taxa that are chosen following a 20MY correction (center column), and the taxa which are chosen following the 30MY threshold correction (right column).

The final data set is composed of 1,953 orthologous protein groups with each individual protein containing 50 to 60 taxa total.

#### B. ERCs on The Original Phylogeny with Short Branches

ERCs were initially calculated for the 1,953 proteins using the complete mammalian phylogeny (Fig. S1) using the same scheme as defined in Methods section of the main text. The top 40 ERCs for ACE2 using this initial method are shown in Table S3. However, these ERCs could be driven (in part) by a spurious correlation to branch time (Section C) An initial attempt to remove the correlation was conducted using partial correlations(Kim, 2015) (Section D). The top 40 ACE2 ERCs for this treatment are also presented in Table S3, along with the final, 30MY threshold corrected ERCs. There are 7 proteins (TNFSF18, IFNAR2, GPR141, CLU, F5, SERPINA5, and SLC10A6) that are shared among all three top 40 ACE2 ERCs. Nine proteins are shared between the top 40 original ACE2 ERCs (TSGA13, CLU, F5, GPR141, PLA2G7, SLC10A6, IFNAR2, TNFSF18, and SERPINA5) and the 30MY ERCs, with 8 proteins that are shared between the top 40 ACE2 branch time-corrected ERC and 30MY ERC sets (CLU, F5, COL4A4, GPR141, SLC10A6, IFNAR2, TNFSF18, and SERPINA5).

| | Original ERCs | | | | BT-Corrected ERCs | | | | 30MY-Adjusted ERCs | | | |
|---|---|---|---|---|---|---|---|---|---|---|---|---|
| Rank | Protein | $\rho$ | P | FDR P | Protein | $\rho$ | P | FDR P | Protein | $\rho$ | P | FDR P |
| 1 | IFNAR2 | 0.711 | 2.0E-15 | 8.8E-13 | PLA2R1 | 0.567 | 6.2E-10 | 1.2E-06 | GEN1 | 0.669 | 4.3E-08 | 4.2E-05 |
| 2 | APOB | 0.709 | 5.5E-17 | 1.1E-13 | APOB | 0.543 | 3.8E-09 | 3.7E-06 | XCR1 | 0.669 | 3.2E-08 | 4.2E-05 |
| 3 | TNFSF18 | 0.700 | 9.9E-16 | 6.5E-13 | CERS3 | 0.537 | 6.0E-09 | 3.9E-06 | CLU | 0.631 | 3.1E-07 | 1.5E-04 |
| 4 | OSMR | 0.699 | 5.8E-16 | 5.6E-13 | IFNAR2 | 0.521 | 1.2E-07 | 3.3E-05 | TMEM63C | 0.630 | 2.0E-07 | 1.3E-04 |
| 5 | CERS3 | 0.682 | 2.3E-15 | 8.8E-13 | WRN | 0.500 | 8.5E-08 | 3.3E-05 | IFNAR2 | 0.616 | 2.5E-06 | 6.1E-04 |
| 6 | SERPINA5 | 0.661 | 1.3E-13 | 2.1E-11 | RMI1 | 0.499 | 1.1E-07 | 3.3E-05 | KIF3B | 0.599 | 1.7E-06 | 4.9E-04 |
| 7 | PLA2G7 | 0.658 | 7.5E-14 | 1.8E-11 | SPHKAP | 0.498 | 1.0E-07 | 3.3E-05 | ITPRIPL2 | 0.590 | 1.7E-06 | 4.9E-04 |
| 8 | LIFR | 0.657 | 1.1E-13 | 2.1E-11 | GPR183 | 0.495 | 1.4E-07 | 3.4E-05 | FAM227A | 0.589 | 1.8E-06 | 4.9E-04 |
| 9 | SLC51B | 0.656 | 6.9E-13 | 5.4E-11 | OSMR | 0.486 | 3.5E-07 | 6.0E-05 | TLR8 | 0.583 | 3.7E-06 | 7.2E-04 |
| 10 | GPR183 | 0.655 | 8.5E-14 | 1.8E-11 | TNFSF18 | 0.483 | 5.5E-07 | 7.6E-05 | COL4A4 | 0.579 | 3.7E-06 | 7.2E-04 |
| 11 | FGB | 0.655 | 6.5E-14 | 1.8E-11 | COL4A4 | 0.483 | 3.7E-07 | 6.0E-05 | FAM3D | 0.574 | 5.8E-06 | 8.4E-04 |
| 12 | F5 | 0.654 | 7.2E-14 | 1.8E-11 | SLC10A6 | 0.482 | 3.3E-07 | 6.0E-05 | F5 | 0.572 | 4.1E-06 | 7.2E-04 |
| 13 | RMI1 | 0.651 | 1.2E-13 | 2.1E-11 | F5 | 0.480 | 3.3E-07 | 6.0E-05 | AR | 0.572 | 7.7E-06 | 8.8E-04 |
| 14 | CPB2 | 0.649 | 3.6E-13 | 3.9E-11 | EPB42 | 0.479 | 4.0E-07 | 6.1E-05 | TSGA13 | 0.569 | 7.1E-06 | 8.8E-04 |
| 15 | CLU | 0.649 | 4.9E-13 | 4.2E-11 | ZNF830 | 0.478 | 1.1E-06 | 1.0E-04 | PLA2G7 | 0.568 | 6.0E-06 | 8.4E-04 |
| 16 | TSGA13 | 0.649 | 2.9E-13 | 3.5E-11 | SERPINA5 | 0.473 | 1.0E-06 | 1.0E-04 | MMS19 | 0.564 | 5.9E-06 | 8.4E-04 |
| 17 | PROCR | 0.647 | 1.9E-13 | 2.9E-11 | CIP2A | 0.470 | 6.2E-07 | 8.1E-05 | AMOT | 0.562 | 8.1E-06 | 8.8E-04 |
| 18 | ZNF830 | 0.646 | 1.6E-12 | 9.4E-11 | C5orf34 | 0.469 | 7.6E-07 | 8.7E-05 | L1CAM | 0.560 | 8.6E-06 | 8.8E-04 |
| 19 | C5orf34 | 0.646 | 2.4E-13 | 3.2E-11 | CAT | 0.468 | 7.1E-07 | 8.7E-05 | PDYN | 0.560 | 7.3E-06 | 8.8E-04 |
| 20 | GPR141 | 0.645 | 3.2E-13 | 3.7E-11 | MUC15 | 0.465 | 1.3E-06 | 1.1E-04 | IQCD | 0.559 | 9.2E-06 | 8.9E-04 |
| 21 | CXCL10 | 0.645 | 2.5E-13 | 3.2E-11 | VTN | 0.464 | 1.2E-06 | 1.0E-04 | SERPINA5 | 0.557 | 2.2E-05 | 1.4E-03 |
| 22 | PLA2R1 | 0.640 | 4.5E-13 | 4.2E-11 | SELP | 0.461 | 1.1E-06 | 1.0E-04 | CERS4 | 0.555 | 2.9E-05 | 1.5E-03 |
| 23 | EPB42 | 0.639 | 4.8E-13 | 4.2E-11 | PPP1R3A | 0.460 | 1.1E-06 | 1.0E-04 | CC2D1B | 0.555 | 1.1E-05 | 1.0E-03 |
| 24 | WRN | 0.637 | 4.7E-13 | 4.2E-11 | PLG | 0.459 | 1.1E-05 | 3.8E-04 | GPR141 | 0.552 | 1.5E-05 | 1.2E-03 |
| 25 | IFIH1 | 0.637 | 4.7E-13 | 4.2E-11 | KITLG | 0.455 | 1.8E-06 | 1.4E-04 | FSCB | 0.551 | 2.8E-05 | 1.5E-03 |
| 26 | BDKRB2 | 0.635 | 1.3E-12 | 8.5E-11 | FGB | 0.454 | 1.6E-06 | 1.3E-04 | RGR | 0.549 | 3.0E-05 | 1.5E-03 |
| 27 | IL1B | 0.635 | 1.3E-12 | 8.6E-11 | FAM237A | 0.454 | 3.0E-06 | 2.0E-04 | COL4A5 | 0.549 | 2.1E-05 | 1.4E-03 |
| 28 | BVES | 0.634 | 6.3E-13 | 5.2E-11 | GPR141 | 0.452 | 2.3E-06 | 1.7E-04 | TNFSF8 | 0.548 | 1.2E-05 | 1.1E-03 |
| 29 | COL1A2 | 0.633 | 1.3E-12 | 8.5E-11 | APOBR | 0.451 | 5.0E-06 | 2.4E-04 | CCDC36 | 0.548 | 1.5E-05 | 1.2E-03 |
| 30 | CAT | 0.632 | 7.9E-13 | 6.0E-11 | CXCL10 | 0.450 | 2.3E-06 | 1.7E-04 | MRC1 | 0.548 | 1.3E-05 | 1.1E-03 |
| 31 | TLR7 | 0.632 | 1.4E-12 | 8.7E-11 | CLU | 0.449 | 4.1E-06 | 2.1E-04 | CD27 | 0.545 | 3.0E-05 | 1.5E-03 |
| 32 | PPP1R3A | 0.632 | 8.5E-13 | 6.1E-11 | ATP10D | 0.448 | 3.0E-06 | 2.0E-04 | ADCK4 | 0.545 | 2.1E-05 | 1.4E-03 |
| 33 | HK3 | 0.629 | 1.8E-12 | 1.0E-10 | LIFR | 0.446 | 3.7E-06 | 2.0E-04 | SOWAHA | 0.543 | 2.2E-05 | 1.4E-03 |
| 34 | SLC10A6 | 0.629 | 1.4E-12 | 8.7E-11 | FER1L5 | 0.446 | 3.3E-06 | 2.0E-04 | F2RL2 | 0.539 | 3.7E-05 | 1.7E-03 |
| 35 | VTN | 0.625 | 2.9E-12 | 1.5E-10 | SERTAD4 | 0.446 | 2.6E-06 | 1.8E-04 | WDR66 | 0.536 | 2.1E-05 | 1.4E-03 |
| 36 | BCLAF3 | 0.624 | 2.4E-12 | 1.3E-10 | PPP2R3A | 0.443 | 3.6E-06 | 2.0E-04 | TRADD | 0.535 | 2.6E-05 | 1.5E-03 |
| 37 | REL | 0.623 | 2.0E-12 | 1.1E-10 | IL5RA | 0.442 | 4.6E-06 | 2.2E-04 | RELA | 0.533 | 2.8E-05 | 1.5E-03 |
| 38 | RNASEL | 0.621 | 3.2E-12 | 1.7E-10 | PIGV | 0.442 | 3.7E-06 | 2.0E-04 | SLC10A6 | 0.531 | 3.0E-05 | 1.5E-03 |
| 39 | APOBR | 0.621 | 2.0E-11 | 7.1E-10 | ZDHHC4 | 0.442 | 3.7E-06 | 2.0E-04 | IL23A | 0.529 | 4.7E-05 | 1.7E-03 |
| 40 | SELENOP | 0.618 | 5.6E-12 | 2.7E-10 | CPED1 | 0.441 | 3.5E-06 | 2.0E-04 | TNFSF18 | 0.528 | 5.8E-05 | 1.8E-03 |

Table S3: The top 40 ERCs for ACE2 based on the original ERC method (left), BT-Corrected partial correlation ERC method (center), and the standard 30MY-adjusted ERC method (right). FDR corrections are based on the full ERC dataset for each respective ERC method.

#### C. Branch Time to Protein-Rate Correlation Problem

In examining the terminal branch rate correlation data for ACE2, we found that its rate of evolution was correlated with the terminal branch time (BT) (illustrated in Fig. S3). We suspect that this correlation may be due to episodic selection over the course of its evolution (possibly driven in part by evolution in its partners). As a result, BT could be a confounding correlate in ERC. Examination of the proteins in our set indicated a significant BT correlation to evolutionary rate for 1,559 out of 1,953 proteins ($p < 0.05$; Supplementary File S6). Notably, many of the strongest original ERCs to ACE2 (such as IFNAR2 and APOB), have very significant correlations to BT with ρ values greater than 0.5 (Table S4). To directly test the effects of time on predicted ERC interactions, multiple linear regressions were performed on the rank-transformed rate data from protein relationships of interest, with time as a covariate (equations in the form: $Protein_{RateRank} = \beta_2 ACE2_{RateRank} + \beta_1 BranchTime_{Rank} + \beta_0$). Many of the proteins with strong ACE2 ERCs resulted in models with the time variable being a significant factor (Table S5). These results additionally hold using similar models under an ANOVA test (Table S5). Examining scatterplots of protein evolutionary rates indicate that the pattern may be driven by short branches with respect to BT (examples in Fig. S3). As expected by this interpretation, the vast majority of proteins (all but 37 of 1953; Supplementary File S7) show significantly more points below the regression line for short branches (<30MY). The short branches occur in relatively oversampled taxonomic orders, as oversampling of closely related species shortens terminal branch times. Since BT is a significant covariate in the original ERC data, the significant ERCs could be due, in part, to a confounding covariance to BT. We therefore examined different approaches to remove this confounding variable (below).

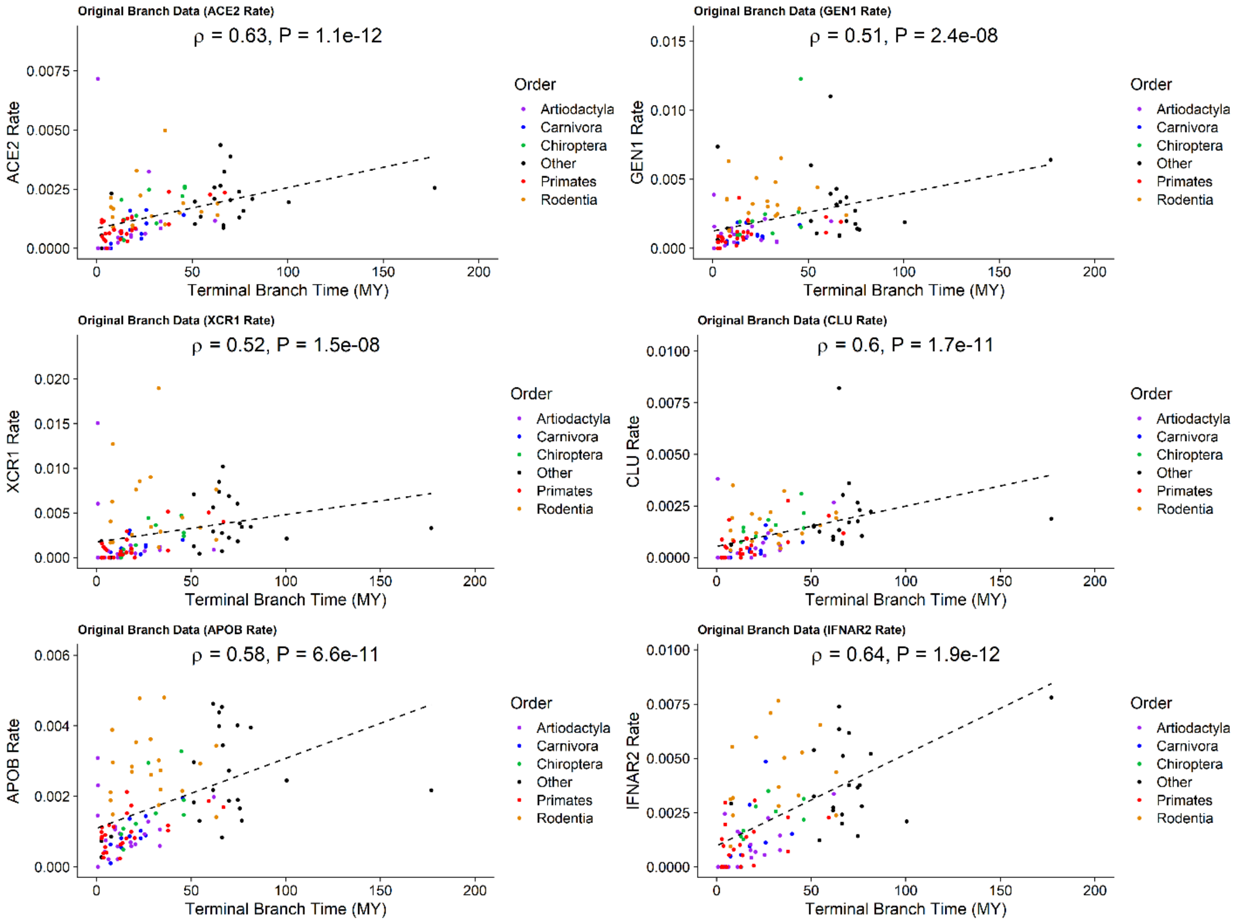


Figure S3: A set of scatterplots depicting the rates of evolution of several proteins of interest plotted against terminal branch time for the original data, with highly sampled clades colored. Also depicted is the linear regression line to emphasize the positive association and Spearman's rank correlation test results (ρ and p-value). In each case, the rate data shows a significant correlation with BT. For each protein, there are significantly more points below the regression line for terminal branches <30MY, indicating lower rates for short branches.

| Protein | $\rho$ | P-Value |
|---|---|---|
| ACE2 | 0.629 | 1.10E-12 |
| GEN1 | 0.512 | 2.38E-08 |
| XCR1 | 0.521 | 1.45E-08 |
| CLU | 0.605 | 1.71E-11 |
| TMEM63C | 0.440 | 1.87E-06 |
| IFNAR2 | 0.641 | 1.92E-12 |
| KIF3B | 0.117 | 2.32E-01 |
| ITPRIPL2 | 0.534 | 2.67E-09 |
| FAM227A | 0.540 | 1.57E-09 |
| TLR8 | 0.459 | 1.07E-06 |
| COL4A4 | 0.384 | 5.23E-05 |
| APOB | 0.576 | 6.64E-11 |
| PLA2R1 | 0.375 | 6.87E-05 |
| CAT | 0.515 | 1.20E-08 |
| CERS3 | 0.498 | 4.30E-08 |

Table S4: Spearman's rank correlation tests on the terminal branch rates against BT for branch time uncorrected data to proteins of interest (strong ERCs in the original or 30MY threshold ERCs). In all cases shown, the proteins have a strong correlation between their terminal branch rates and time prior to correction for short branches.

| Protein | $R^2_{adj}$ | Model P | Intercept Term | Intercept P | ACE2 Term | ACE2 P | Time Term | Time P | ANOVA ACE2 P | ANOVA Time P |
|---|---|---|---|---|---|---|---|---|---|---|
| GEN1 | 0.382 | 2.68E-11 | 15.672 | 2.18E-03 | 0.449 | 2.48E-05 | 0.241 | 1.92E-02 | 2.00E-11 | 1.92E-02 |
| XCR1 | 0.341 | 6.02E-10 | 17.340 | 1.05E-03 | 0.384 | 4.37E-04 | 0.272 | 1.13E-02 | 7.58E-10 | 1.13E-02 |
| CLU | 0.476 | 1.72E-14 | 11.114 | 1.66E-02 | 0.453 | 4.07E-06 | 0.323 | 7.49E-04 | 5.19E-14 | 7.49E-04 |
| TMEM63C | 0.222 | 1.33E-06 | 24.124 | 4.63E-05 | 0.204 | 7.20E-02 | 0.332 | 3.94E-03 | 7.41E-06 | 3.94E-03 |
| IFNAR2 | 0.556 | 7.33E-17 | 7.939 | 5.32E-02 | 0.515 | 1.18E-07 | 0.314 | 6.92E-04 | 1.33E-16 | 6.92E-04 |
| KIF3B | 0.125 | 5.21E-04 | 36.182 | 2.42E-08 | 0.458 | 2.12E-04 | -0.168 | 1.62E-01 | 2.60E-04 | 1.62E-01 |
| ITPRIPL2 | 0.292 | 1.21E-08 | 20.694 | 2.24E-04 | 0.183 | 9.12E-02 | 0.419 | 1.69E-04 | 5.37E-07 | 1.69E-04 |
| FAM227A | 0.339 | 3.94E-10 | 17.841 | 9.16E-04 | 0.330 | 1.92E-03 | 0.327 | 2.14E-03 | 1.55E-09 | 2.14E-03 |
| TLR8 | 0.322 | 2.93E-09 | 18.343 | 6.31E-04 | 0.433 | 9.81E-05 | 0.200 | 6.37E-02 | 1.35E-09 | 6.37E-02 |
| COL4A4 | 0.327 | 1.35E-09 | 21.071 | 1.04E-04 | 0.578 | 3.68E-07 | 0.008 | 9.37E-01 | 1.83E-10 | 9.37E-01 |
| APOB | 0.523 | 3.15E-17 | 10.849 | 1.62E-02 | 0.569 | 3.79E-09 | 0.223 | 1.29E-02 | 1.58E-17 | 1.29E-02 |
| PLA2R1 | 0.400 | 3.98E-12 | 19.693 | 1.34E-04 | 0.677 | 6.17E-10 | -0.059 | 5.49E-01 | 5.45E-13 | 5.49E-01 |
| CAT | 0.408 | 1.58E-12 | 15.654 | 2.03E-03 | 0.519 | 7.13E-07 | 0.180 | 6.94E-02 | 5.22E-13 | 6.94E-02 |
| CERS3 | 0.467 | 8.43E-15 | 13.804 | 4.03E-03 | 0.592 | 6.02E-09 | 0.143 | 1.29E-01 | 1.79E-15 | 1.29E-01 |

Table S5: Linear model fit using the original data set to test for branch time and ACE2 effects, using the form: $Protein_{RateRank} = \beta_2 ACE2_{RateRank} + \beta_1 BranchTime_{Rank} + \beta_0$ . Selected proteins of interest are shown from top ACE2 ERCs of the original and 30MY data sets. In all cases, except for TMEM63C and ITPRIPL2, the model has a strongly significant reported P-value, indicating that ACE2 is significantly predictive. For 8 of 14 proteins branch time is also significantly predictive. For ANOVA, all 14 proteins show a significant ACE2 effect, and 8 of 14 have a significant Branch time effect. This indicates that branch time is a confounding factor for many ACE2's ERCs in the original data, which contains short terminal branches.

#### D. Partial Correlation to Address BT-PR Correlation

As time is a significant confounding effect on the protein rate, ERCs values may be distorted by the branch time covariate. We, therefore, investigated the use of "partial correlations" to control for the confounding effect of time on our correlation calculations (Kim, 2015). Partial correlation-based ERCs were generated utilizing the "ppcor" R package (Kim, 2015) to produce Spearman's rank partial correlation tests while controlling for the effects of terminal branch time. The partial correlations are based on fitting a linear model to the variable(s) being controlled for and then performing a Spearman's rank correlation test on the residuals of the two models. These residuals represent the variance in the data that are unexplained by the variable(s) being controlled for. In particular, terminal branch time was controlled to account for the observed correlation to BT. Even following the partial correlation controlling for BT, ACE2 still had strong ERCs to immune system-related proteins such as IFNAR2 (Table S3). However, partial correlations are not robust to assumption violations. As partial correlations are based on performing a rank correlation test on the residuals of linear models of rates trained against time, we examined the data to assess the possibility of these violations. Several problems were noted upon examining residuals of individually trained models. The most important of which is that rate vs BT residuals were still correlated with BT. Since these residuals should capture variance that is not explained by terminal branch time, it is unexpected for these residuals to still have a strong association to BT. However, 1,529 of 1,953 proteins have residuals that still have a significant Spearman's correlation to time ($p < 0.05$ Supplementary File S8, select proteins are displayed in Table S6). Key proteins such as ACE2 are among the set of proteins with residuals that still correlate significantly to BT (Table S6). The previous analysis showed that short branch rates are overrepresented below the protein rate to branch time regression line for the vast many proteins, which likely explains why the partial regression fails to remove the branch time correlation in many cases.

| Protein | Residuals vs Time $\rho$ | Residuals vs BTime P |
|---|---|---|
| ACE2 | 0.197 | 4.60E-02 |
| GEN1 | -0.006 | 9.51E-01 |
| XCR1 | 0.133 | 1.77E-01 |
| CLU | 0.092 | 3.55E-01 |
| TMEM63C | 0.374 | 6.55E-05 |
| IFNAR2 | 0.081 | 4.35E-01 |
| KIF3B | 0.519 | 1.17E-08 |
| ITPRIPL2 | 0.365 | 1.03E-04 |
| FAM227A | 0.284 | 2.86E-03 |
| TLR8 | 0.200 | 4.23E-02 |
| COL4A4 | 0.546 | 1.75E-09 |
| APOB | 0.114 | 2.41E-01 |
| PLA2R1 | 0.327 | 5.75E-04 |
| CAT | 0.408 | 1.14E-05 |
| CERS3 | 0.299 | 1.67E-03 |

Table S6: Spearman's rank correlation tests of the residuals of linear models trained on a protein's rates against time. Ten of the 15 proteins depicted (including ACE2) retain a significant association with time after accounting for time. Full table available in Supplementary File S8.

As we noted that short branches appear to drive the rate to BT correlation (Fig. S3), we therefore decided to control for confounding branch time effects by removing short branches and recalculating ERC rates.

#### E. Removing Short Branches to Remove the Confounding BT-Rate Factor

As we observed that terminal branch time is a confounding factor in our ERC analysis (Section C), we examined short branches as a likely driver for the association. Therefore, we identified sister taxa with short branches and selectively remove one or more, to remove short branches and extend branches in the remaining sister taxa (Fig. S2, Table S2). The procedure was applied to produce clades with branch lengths with a 20MY BT threshold or a 30MY BT threshold. Note that we allowed around a 3 MY buffer (e.g. 30-27 MY threshold) so as to not restrict the taxonomic sample sizes too heavily. The specific representative taxa were picked arbitrarily, but generally were chosen to allow for the greatest number of internal nodes to be merged into a single branch (Table S2), with the main exception to the rule being that *Homo sapiens* was selected as the representative of its clade, due to its relevance to the COVID-19. The taxa selections at the 20MY time scale resulted in the removal of 32 taxa from the original phylogeny and the taxa selections at the 30MY time scale resulted in the removal of 48 taxa. Both adjustments to the data strongly reduced the number of proteins displaying a significant association between rate and BT. Specifically, while the original data set had 1,559 out of 1,953 proteins which displayed a significant correlation between BT and rate ($p < 0.05$), the 20MY adjustment reduced this number to 1,065 proteins, and the 30MY adjustment reduced the number of proteins with a significant rate to BT correlation to 245 (select proteins in Table S7, complete set in Supplementary File S6), or 12.5% of proteins. Therefore, the 30MY terminal branch length threshold most effectively removed branch time as a confounding factor. After the 30MY correction, there is no longer a significant correlation between branch time and branch rate for most proteins, as illustrated in Table S7 and Figure S4.

| | Full Rate-Btime | | 20MY Rate-Btime | | 30MY Rate-Btime | |
|---|---|---|---|---|---|---|
| **Protein** | $\rho$ | **P-Value** | $\rho$ | **P-Value** | $\rho$ | **P-Value** |
| ACE2 | 0.629 | 1.10E-12 | 0.484 | 1.86E-05 | 0.208 | 1.24E-01 |
| GEN1 | 0.512 | 2.38E-08 | 0.381 | 8.73E-04 | 0.075 | 5.78E-01 |
| XCR1 | 0.521 | 1.45E-08 | 0.436 | 1.32E-04 | 0.116 | 3.86E-01 |
| CLU | 0.605 | 1.71E-11 | 0.553 | 3.91E-07 | 0.297 | 2.36E-02 |
| TMEM63C | 0.440 | 1.87E-06 | 0.248 | 3.07E-02 | 0.189 | 1.49E-01 |
| IFNAR2 | 0.641 | 1.92E-12 | 0.377 | 1.94E-03 | 0.263 | 5.67E-02 |
| KIF3B | 0.117 | 2.32E-01 | 0.010 | 9.35E-01 | 0.164 | 2.17E-01 |
| ITPRIPL2 | 0.534 | 2.67E-09 | 0.498 | 4.60E-06 | 0.221 | 8.94E-02 |
| FAM227A | 0.540 | 1.57E-09 | 0.293 | 1.03E-02 | 0.192 | 1.42E-01 |
| TLR8 | 0.459 | 1.07E-06 | 0.462 | 4.33E-05 | 0.241 | 7.05E-02 |
| COL4A4 | 0.384 | 5.23E-05 | 0.397 | 5.05E-04 | 0.132 | 3.20E-01 |
| APOB | 0.576 | 6.64E-11 | 0.488 | 7.69E-06 | 0.207 | 1.12E-01 |
| PLA2R1 | 0.375 | 6.87E-05 | 0.309 | 6.93E-03 | 0.014 | 9.14E-01 |
| CAT | 0.515 | 1.20E-08 | 0.246 | 3.25E-02 | 0.046 | 7.27E-01 |
| CERS3 | 0.498 | 4.30E-08 | 0.316 | 5.48E-03 | -0.048 | 7.15E-01 |

Table S7: Spearman's rank correlation tests on the terminal branch rates versus branch time for proteins of interest for the three different time threshold treatments: No cutoff, 20MY cutoff, 30MY cutoff. In all cases, the correlation with rate and time decrease—to the point where unadjusted p-values are insignificant at $p < 0.05$ level for all but one protein at the 30MY cutoff.

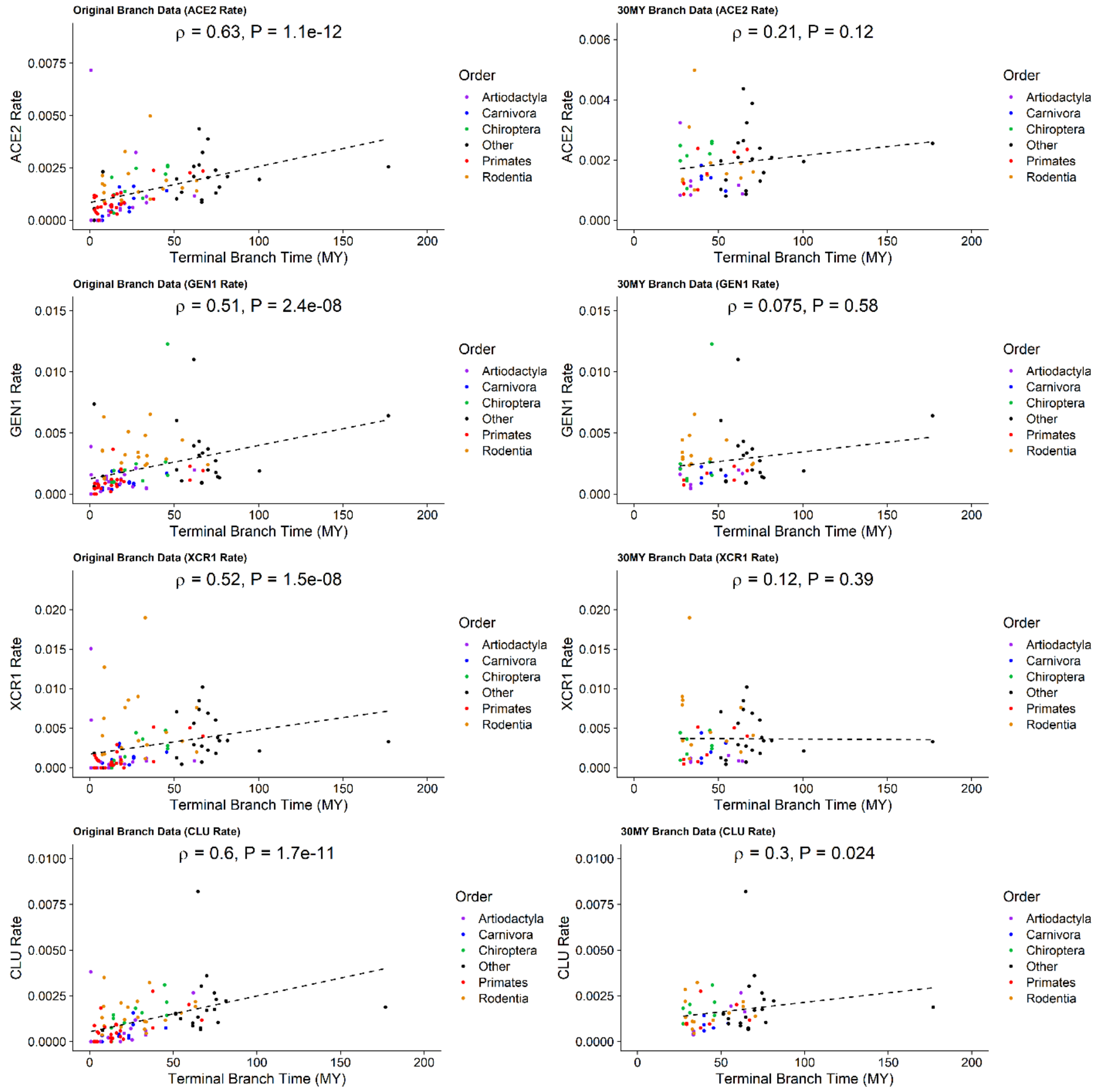


Figure S4: A set of scatterplots depicting the rate of evolution of several proteins of interest plotted against terminal branch time with highly sampled clades colored. The left column of plots depicts the original rate data and the right column depicts the corresponding rate data following a 30MY adjustment. Also depicted is the regression line to emphasize the positive association and the statistics of Spearman's rank correlation test results (ρ and p-value). In each case, the original data shows a significant correlation with BT while the 30MY adjusted data shows that the association is no longer significant.

Results from the 30MY adjustment also reveal strong reciprocal ERCs among proteins known to occur in complex with each other that were not apparent in the uncorrected ERC analysis. For instance, the three fibrinogen subunits FGA, FGB, and FGG form a well-known fibrinogen complex (Mosesson, 2005), and have strong reciprocal rank ERCs in the 30MY data, but do not in the original treatment (Tables S8-S10). Similar empirical observations were noted among several other interacting proteins such as the weak relationship between IFNAR1 and IFNAR2 in the uncorrected data but the much stronger relationship in the 30MY data (Table S11), despite their being known to complex (Thomas et al., 2011). We also note weak relationships between several of the Collagen Type IV subunits in the uncorrected ERC data, but the relationships were again strengthened following the 30MY adjustment (Table S12) which are known to physically interact (Casino et al., 2018), and found to form strong reciprocal rank ERCs in the corrected data set.

| FGA and FGB | | | | |
|---|---|---|---|---|
| **ERC** | $\rho$ | **Raw P** | **Rank of FGB for FGA** | **Rank of FGA for FGB** |
| Original | 0.703 | 8.5E-17 | 48 | 6 |
| Time-Corrected | 0.575 | 2.1E-10 | 62 | 5 |
| 30MY-Corrected | 0.696 | 9.6E-10 | 12 | 1 |

Table S8: The ERC results between the expected interacting proteins FGA and FGB under the original ERC method, the time-corrected partial correlation-based ERC, and the final 30MY-corrected ERC. This interaction does not meet our reciprocal rank 20 criteria until we use the 30MY-corrected ERCs.

| FGA and FGG | | | | |
|---|---|---|---|---|
| **ERC** | $\rho$ | **Raw P** | **Rank of FGG for FGA** | **Rank of FGA for FGG** |
| Original | 0.661 | 3.8E-14 | 109 | 5 |
| Time-Corrected | 0.563 | 9.2E-10 | 77 | 5 |
| 30MY-Corrected | 0.722 | 1.6E-10 | 2 | 1 |

Table S9: The ERC results between the expected interacting proteins FGA and FGG under the original ERC method, the time-corrected partial correlation-based ERC, and the final 30MY-corrected ERC. This interaction does not meet our reciprocal rank 20 criteria until we use the 30MY-corrected ERCs, additionally, the 30MY ERC value itself is strongest after the 30MY correction.

| FGB and FGG | | | | |
|---|---|---|---|---|
| **ERC** | $\rho$ | **Raw P** | **Rank of FGG for FGB** | **Rank of FGB for FGG** |
| Original | 0.670 | 5.4E-15 | 23 | 4 |
| Time-Corrected | 0.568 | 3.3E-10 | 11 | 4 |
| 30MY-Corrected | 0.603 | 4.3E-07 | 4 | 7 |

Table S10: The ERC results between the expected interacting proteins FGB and FGG under the original ERC method, the time-corrected partial correlation-based ERC, and the final 30MY-corrected ERC. This interaction does not meet our reciprocal rank 20 criteria using the original ERC calculation. It does meet the reciprocal rank 20 criteria after time correction, but this reciprocal rank interaction gets even stronger after the 30MY correction.

| IFNAR1 and IFNAR2 | | | | |
|---|---|---|---|---|
| **ERC** | $\rho$ | **Raw P** | **Rank of IFNAR2 for IFNAR1** | **Rank of IFNAR1 for IFNAR2** |
| Original | 0.635 | 1.8E-11 | 52 | 215 |
| Time-Corrected | 0.497 | 7.1E-07 | 131 | 167 |
| 30MY-Corrected | 0.786 | 2.1E-11 | 2 | 18 |

Table S11: The ERC results between the expected interacting proteins IFNAR1 and IFNAR2 under the original ERC method, the time-corrected partial correlation-based ERC, and the final 30MY-corrected ERC. Notably, the interaction does not meet our reciprocal rank 20 criteria until our 30MY correction. We also note that the 30MY ERC is stronger than all other attempts.

| Interacting Collagen Type IV Pairs | | | |
|---|---|---|---|
| **Protein Pairs** | **Original ERC Ranks** | **Time-Corrected ERC Ranks** | **30MY-Corrected ERC Ranks** |
| COL4A1 - COL4A2 | 134 - 70 | 157 - 54 | 7 - 22 |
| COL4A3 - COL4A4 | 48 - 57 | 40 - 38 | 1 - 1 |
| COL4A3 - COL4A5 | 414 - 262 | 423 - 303 | 115 - 84 |
| COL4A4 - COL4A5 | 228 - 123 | 189 - 115 | 105 - 74 |
| COL4A5 - COL4A6 | 7 - 33 | 4 - 23 | 7 - 18 |

Table S12: The ERC ranks of protein pairs of interacting Collagen Type IV subunits according to Casino et al. (2018) under different ERC corrections. P and p-values are omitted for clarity but in all, instances, the ρ values were increased under the 30MY correction when compared to either the time-corrected or original ERCs.

#### F. Testing Whether Branch Rate Increases with Evolutionary Time

There is a positive association between terminal branch time and the rate of evolution for many proteins (Section C). The question, therefore, arises as to whether there is actually an increase in evolutionary rate over time for these proteins. To test this question, we conducted an "experiment" to extend branches along independent clades, in order to test whether increasing branch time increases protein evolutionary rate. This was accomplished by extending branch lengths along taxonomic branches in different clades by trimming adjacent taxa and comparing the protein rates as branches are extended. (Fig. S5). Based on the TimeTree phylogeny (Kumar et al., 2017), we selected individual clades containing short branches that would have their time scales extended following a 20MY and 30MY adjustment (Fig. S5, Table S2).

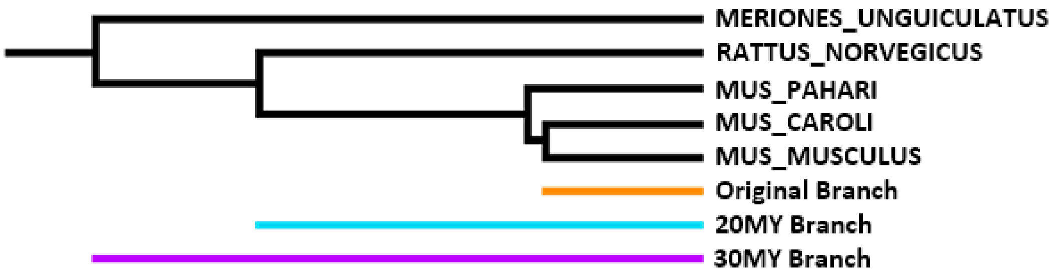


Figure S5: Cartoon illustrating the branches being compared when testing whether branch rates change upon an increase in time scale. In this instance, the taxon "Mus musculus" is selected from the Rattus and Mus clades. The original short branch (orange), 20MY branch (cyan), and 30MY branch (purple) are each used to calculate rates, and these are the paired data that is compared to test for changes in rates.

Since we suspected that rates scale as time increases, we specifically tested whether there is a significant difference in rate for each of these branches before and after 20MY and 30MY adjustments, as described in Section E (14 selected taxa for comparing original vs 20MY, 12 selected taxa for comparing 20MY vs 30MY, 16 selected taxa for comparing original vs 30MY). Tests on each branch's rate against the respective adjusted rate were performed using two-tailed Wilcoxon Matched Signed Rank Tests (results for all proteins are reported in Supplementary File S9), to test whether these rates significantly differed. We note that many proteins show significant changes in rate under each adjustment, but this pattern is most prominent in the shift from short branch rates to 30MY rates (longer branches). Examples are shown in Table S13 and Figure S6, and the complete data are present in Supplementary File S9. Notably, out of our set of 1,953 proteins using a significance cutoff of $p < 0.05$, 261 proteins show significant rate changes (238 of which have a median increase in rate) in the Short-to-20MY treatment, 456 show significant rate changes (442 of which have a median increase in rate) in the 20MY-to-30MY treatment, and 551 show significant rate changes (545 of which have a median increase in rate) in the Short-to-30MY treatment (Fig. S7).

| Protein | Short vs 20MY P | 20MY vs 30MY P | Short vs 30MY P |
|---|---|---|---|
| ACE2 | 3.58E-01 | 2.44E-03 | 2.14E-04 |
| GEN1 | 2.45E-02 | 2.10E-02 | 7.63E-04 |
| XCR1 | 1.04E-01 | 3.42E-03 | 1.82E-02 |
| CLU | 1.05E-02 | 6.84E-03 | 1.68E-03 |
| TMEM63C | 4.63E-01 | 2.33E-01 | 1.93E-01 |
| IFNAR2 | 2.44E-04 | 2.50E-01 | 2.44E-03 |
| KIF3B | 9.52E-01 | 7.91E-01 | 8.60E-01 |
| ITPRIPL2 | 1.00E+00 | 5.22E-02 | 4.64E-01 |
| FAM227A | 9.52E-01 | 6.77E-01 | 9.00E-01 |
| TLR8 | 3.53E-02 | 9.77E-04 | 4.27E-03 |
| COL4A4 | 5.83E-01 | 3.22E-02 | 7.39E-02 |
| APOB | 1.66E-02 | 9.77E-04 | 9.16E-05 |
| PLA2R1 | 5.83E-01 | 9.77E-04 | 5.77E-02 |
| CAT | 8.54E-03 | 1.10E-01 | 1.68E-03 |
| CERS3 | 1.94E-01 | 4.88E-04 | 9.19E-03 |

Table S13: Unadjusted P-values for two-tailed paired Wilcoxon signed-rank tests comparing the rate of evolution of selected branches after various adjustments for selected proteins of interest. Most proteins show significant differences in rate, and all but PLA2R1 has a significant difference in rates from the original rate data and 30MY rate data.

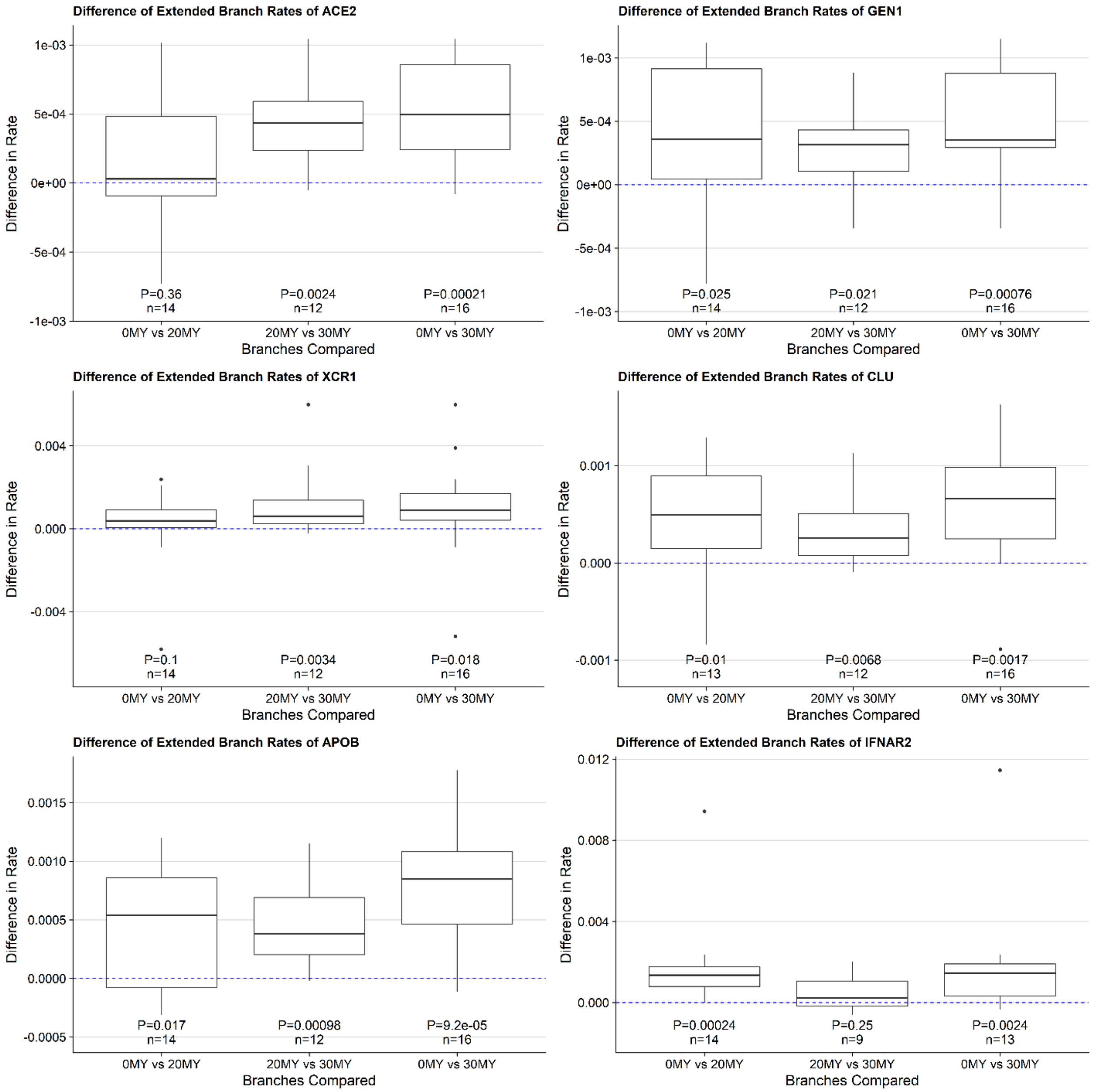


Figure S6: Boxplots of the differences in the rate of evolution of selected branches after various adjustments for selected proteins of interest. A dashed blue line indicates a difference of zero. Sample size and two-tailed paired Wilcoxon signed-rank test p-values are indicated underneath each respective box indicating if there was a significant change in rates.

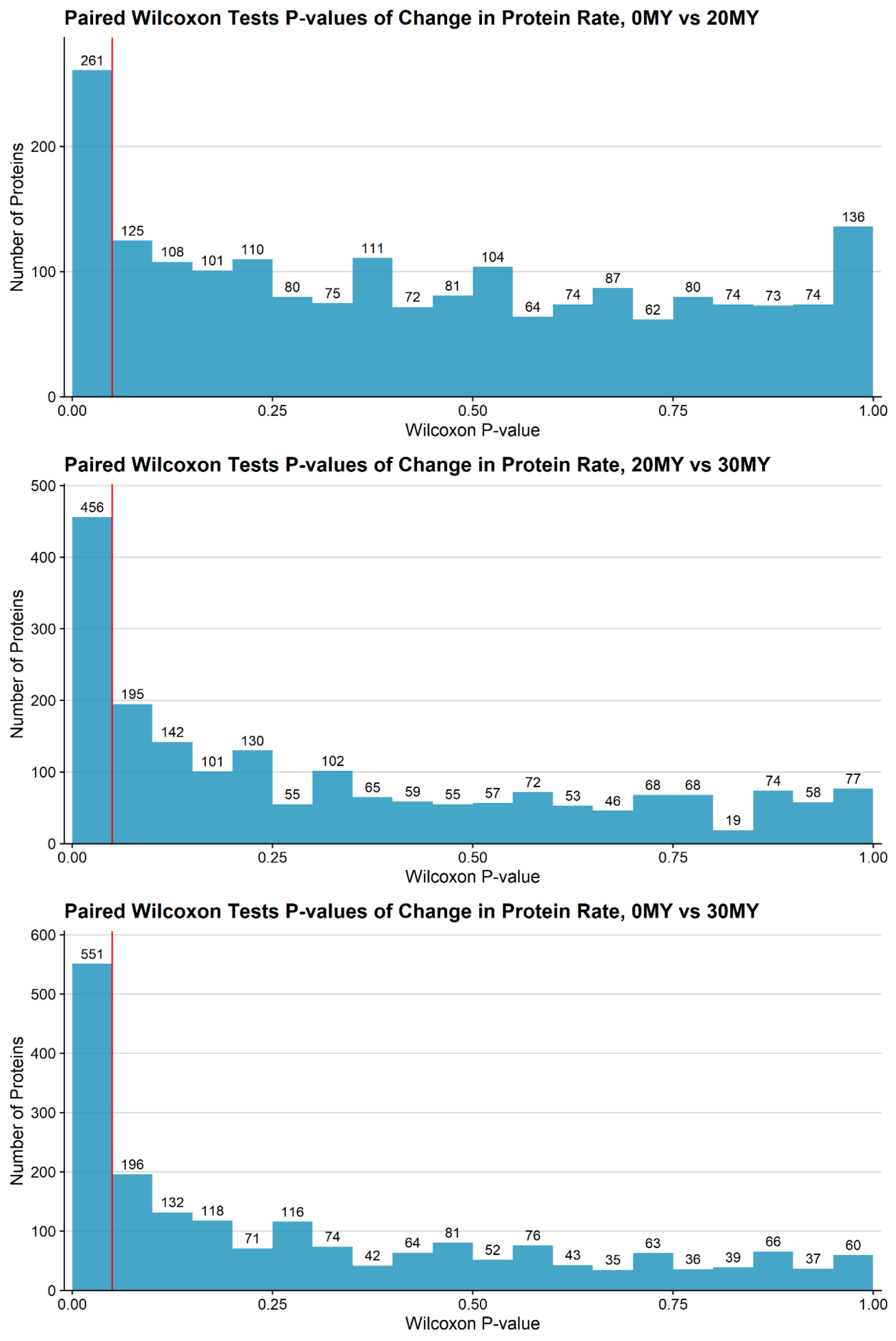


Figure S7: The distributions of p-values of the two-tailed Wilcoxon matched signed-rank tests comparing whether there is a significant difference in the rates of difference in selected branches when time scales were increased. Additionally, the vertical red line indicates a $p < 0.05$ threshold for significance, such that all bins to the right of it represent insignificant tests.

We hypothesize that these shifts in rate may be due to increased evolutionary time scales being able to capture episodic evolutionary events that would otherwise be missed in the short branches of the original phylogeny. As longer time scales are considered, there could be a larger chance that these episodic events would be captured, explaining the pattern.

#### G. Testing for Taxonomic Order Effects

We use three methods to test for taxonomic order effects on the calculated 30MY ERCs, (1) multiple linear regression, (2) analysis of covariance (ANCOVA), and (3) non-parametric independent contrasts. For the regression and ANCOVA approaches, 30MY rate data is grouped by mammalian taxonomic orders accessed via ETE3 (Huerta-Cepas, Serra & Bork, 2016) and treated as an independent variable. The independent contrasts test uses the mammalian topology previously created with TimeTree (Kumar et al., 2017) to generate independent contrasts within the phylogeny. Statistical tests for each method are performed using base R (version 3.6.1).

Linear regression models using mammalian order as a variable were tested in the following general equation format: $Protein_{RateRank} = \beta_3 ACE2_{RateRank} + \beta_2 BranchTime_{Rank} + \beta_1 Order + \beta_0$ on the 30MY adjusted terminal branch time data. Since taxonomic order is a categorical variable, R implicitly converts the variable to become a one-hot encoded "contrast" matrix. One can then examine the reported model metrics to see if any of the encoded taxonomic order variables have a statistically significant contribution to the resultant model. We focus our analysis on the top 5 proteins showing high 30MY ERCs with ACE2 (GEN1, XCR1, CLU, TMEM63C, and IFNAR2). All the examined models have a strong fit (Table S14). In most cases, none of the orders provide a significant contribution to the model (Supplementary File S10). There are a few notable exceptions. The model for GEN1 displays a near-significant contribution of Rodentia, but removing Rodentia still results in a significant ERC to ACE2 ($\rho = 0.60$, unadjusted $p = 4.5E-11$) so the ERC is not an artifact of the effect of Rodentia. Additionally, the model for CLU displays a significant contribution of Dasyuromorphia (Supplementary File S10), however, there is only one taxon within the order in the data and there is still a strong ERC when this taxon is removed ($\rho = 0.67$, unadjusted $p = 3.2E-08$). So, we do not consider this an important contributor to the ACE2-CLU relationship, and it is more likely to be due to model overfitting. We also note that IFNAR2's model shows a significant contribution of the Carnivora, Cingulata, Perissodactyla, Pholidota, and Primates. (Supplementary File S9). But the ERC between ACE2 and IFNAR2 is still strong after removing these orders from the 30MY rate data ($\rho = 0.56$, unadjusted $p = 3.7E-04$). Importantly, all the models calculated show a significant contribution of ACE2, even in the presence of these order effects (p-values range from 2.04E-02 to 4.1E-04; Supplementary File S9). Furthermore, the linear models for each of these proteins of interest show an insignificant contribution of branch time using the 30MY-based rate data, further validating the removal of the rate-time correlation (Supplementary File S10).

| Protein | $R^2_{Adj}$ | Model P-Value |
|---|---|---|
| GEN1 | 0.678 | 1.04E-06 |
| XCR1 | 0.520 | 2.93E-04 |
| CLU | 0.416 | 2.63E-03 |
| TMEM63C | 0.456 | 9.68E-04 |
| IFNAR2 | 0.724 | 1.51E-06 |

Table S14: The adjusted $R^2$ and overall model significance values for each of the linear models representing ACE2's top 5 ERCs to test for the effects of taxonomic order. In all cases, the model is significant at $p < 0.05$ and has strong fits reported by the $R^2$ values, confirming the relationships identified with the 30MY ERCs between ACE2 and these proteins.

As an alternate method to test for the effects of taxonomic order, we used ANCOVA. ANCOVA is a parametric test that allows for the inclusion of categorical data. Since ANCOVA has a similar model structure as linear modeling, the same model structure described above is once again utilized for statistical testing. ACE2's top 5 ERC partners in the 30MY set have no significant effect of taxonomic order except for GEN1 (p = 1.6E-03; Table S15) and IFNAR2 (p = 2.5E-04; Table S15). However, ACE2 has a much more significant contribution to each of these models than does Order (p = 7.9E-10 for GEN1 and p = 7.8E-09 for IFNAR2; Table S15). Removing the orders identified above in the regression analysis eliminates the significant order effect detected by ANCOVA for GEN1 (p = 2.2E-01) and reduces the effect for IFNAR2 (p = 8.6E-03). But as discussed above, the ERCs for ACE2 to GEN1 and to IFNAR2 are still strong and significant after removing the taxa identified in the regression analysis. We also note again, that under 30MY adjustment, terminal branch time is not a significant covariate in all cases examined (Table S15).

| Protein | ACE2 P | Order P | BTime P |
|---|---|---|---|
| GEN1 | 7.9E-10 | 1.6E-03 | 7.0E-01 |
| XCR1 | 4.8E-08 | 1.3E-01 | 9.9E-01 |
| CLU | 8.2E-07 | 4.9E-01 | 6.4E-02 |
| TMEM63C | 2.9E-07 | 1.8E-01 | 5.8E-01 |
| IFNAR2 | 7.8E-09 | 2.5E-04 | 3.5E-01 |

Table S15: Table showing the p-values for the covariates of ANCOVA tests run on linear models considering the rates of proteins of interest against ACE2 with taxonomic order and terminal branch time.

A Spearman non-parametric independent contrasts test (Garland, Harvey & Ives, 1992) was also used to check for taxonomic effects in the 30MY adjusted rate data. The independent contrasts test is used to examine if there is a significant relationship between ACE2 rates and its top 5 ERC partners even after accounting for taxonomic effects between related species. The test is performed using the R packages "ape" (Paradis & Schliep, 2019) and "picante" (Kembel et al., 2010). In all cases, ACE2 continues to have a significant relationship to each protein ($p < 0.05$), indicating that ACE2's 30MY ERC relationships are not driven by taxonomic bias (Table S16).

| | ACE2 30MY ERCs | | ACE2 Contrasts Correlations | |
|---|---|---|---|---|
| **Protein** | **$\rho$** | **P-Value** | **$\rho$** | **P-Value** |
| GEN1 | 0.669 | 4.27E-08 | 0.673 | 3.39E-08 |
| XCR1 | 0.669 | 3.16E-08 | 0.503 | 1.05E-04 |
| CLU | 0.631 | 3.07E-07 | 0.604 | 1.32E-06 |
| TMEM63C | 0.630 | 2.02E-07 | 0.674 | 1.27E-08 |
| IFNAR2 | 0.616 | 2.48E-06 | 0.613 | 2.80E-06 |

Table S16: Table showing the correlation coefficients and p-values for the Spearman non-parametric independent contrasts tests on ACE2 against the top 5 ACE2 ERC proteins controlling for phylogenetic effects with the use of independent contrasts. In all cases, the proteins retain a strongly significant correlation with ACE2.

#### H. Additional Information on ACE2 Interactor Proteins

Here we provide additional summary information on ACE2 ERC proteins of interest, based on our review of data sources Gene Cards (Stelzer et al., 2016), KEGG (Kanehisa & Goto, 2000), UniProt (Bateman et al., 2021), NCBI Entrez (Maglott et al., 2005), Human Protein Atlas (Thul et al., 2017), and surveys of literature detected through Google Scholar searches. Additional information on the ERC associations of these proteins is also presented below.

*GEN1 (Flap endonuclease GEN homolog 1)*: GEN1 is ACE2's top-ranked ERC ($\rho$ = 0.67, FDR = 4.2E-05). It is a DNA nuclease whose primary functions are the resolution of DNA Holliday junctions (Chan & West, 2015), and DNA damage checkpoint signaling (Palmer & Kaldis, 2020). It also has a role in centromere stability in both meiosis and mitosis (Gao et al., 2012). Consistent with its roles in meiosis and mitosis, the second-highest ERC interactor for GEN1 is CC2D1B, a protein involved in resealing of the nuclear envelope following mitosis and assembly and disassembly of the mitotic spindle (Vietri & Stenmark, 2018).

Surprisingly, the top ERC interactor of GEN1 is Interferon λ receptor 1 (IFNLR1), and they are each other's top-ranked ERC connections (Supplementary File S3). This implies a tight association of GEN1 with the interferon pathways involved in immune response and antiviral defense (Prokunina-Olsson et al., 2020), although there is little evidence for this in the literature. Interferon pathways are important in antiviral defense, but also can contribute to cytokine storms and COVID-19 pathologies (McKechnie & Blish, 2020). Along with SLC10A6 and TESPA1, GEN1, IFNLR1, and CC2D1B form a strong reciprocal rank network (Section D, Figure 3). GEN1's top 2% ERCs are enriched for multiple terms related to viral infection, such as HPV infection (FDR = 2.0E-03), Measles (FDR = 4.0E-03), Hepatitis C (FDR = 4.6E-03), Necroptosis (FDR = 4.6E-03), Influenza A (FDR = 4.7E-03), and Kaposi sarcoma-associated herpesvirus infection (FDR = 5.5E-03). Cytokine-cytokine receptor interaction is another significantly enriched term (FDR = 1.6E-04). In contrast, based on our standard top 2% ERC list for enrichment, there are no significant terms strictly related to DNA replication, despite that being the primary identified function of GEN1 in the scientific literature. We speculate that GEN1's functions in DNA and centrosomes during mitosis could be related to DNA checkpoint signaling affecting apoptosis or necrotic cell death, perhaps explaining the enrichment for proteins involved in viral responses. Identification of binding domains between GEN1 and some of its top ERC partners could be informative for possible functional studies.

*XCR1 (X-C Motif Chemokine Receptor 1)*: XCR1 is the 2$^{nd}$ top-ranked ERC for ACE2 ($\rho$ = 0.67, FDR = 6.18E-05). XCR1 is the receptor for the chemokine XCL1. The receptor-cytokine interplay is involved in the immune response to infection and inflammation, development of regulatory T cells in the thymus, and establishing self-tolerance (Lei & Takahama, 2012). Therefore, disruptions of XCR1 due to protein interactions with ACE2 could play a role in COVID-19 complications. As well as being the top rank ACE2 ERC, these two proteins have reciprocal rank correlations at the 2% level (ACE2 is rank 37 for XCR1). Strikingly, the Severe Covid-19 GWAS Group (2020) detected a small genomic region containing six genes that significantly associates with severe COVID-19, one of which is XCR1. Our finding that ACE2's 2nd highest ERC interactor is also XCR1 is striking for two reasons. First, it lends independent support for a relationship between COVID-19 and XCR1. Second, it implicates that a direct interaction between ACE2 and XCR1 could be involved in COVID-19 pathologies. To our knowledge, there are no other reports of interactions between these two proteins. Its Top 2% ERCs show an extremely strong enrichment for cytokine-cytokine receptor interactions (FDR = 8.0E-06) and JAK-STAT related terms (FDR = 9.7E-03), and for coagulation and complement and cascades (FDR = 1.0E-02).

*CLU (Clusterin, aka Apolipoprotein J)*: CLU is the 3rd highest ACE2 ERC (ρ = 0.63, FDR = 1.5E-04), and these two proteins show strong reciprocal ranks (3, 8), likely supporting biological interactions. Relevant to this point is that both ACE2 and CLU have soluble forms that circulate in the blood (Itakura et al., 2020). CLU prevents aggregation of misfolded proteins in blood by binding to them, and also clears misfolded extracellular proteins by binding to heparan sulfate receptors on cells, leading to endocytosis and degradation of CLU and associated proteins in lysosomes (Itakura et al., 2020). This recently discovered mechanism has been referred to as a "cleaning squad" for extracellular misfolded proteins (Sánchez-Martín & Komatsu, 2020). CLU also protects cells from complement-induced apoptosis and lysis (Jenne & Tschopp, 1989). As well as being abundant in blood plasma, CLU is also found on mature sperm and abundant in seminal plasma (Uhlén et al., 2015).

CLU shows the strongest possible reciprocal ranking with GPR141 (1,1 - ρ = 0.68, FDR = 9.1E-06). GPR141 is associated with megakaryocytes (see below). Consistent with their strong evolutionary correlation, CLU is produced in megakaryocytes which subsequently mature into platelets (Tschopp et al., 1993). CLU is released by activated platelets in surrounding fluids at sites of vascular injury (Witte et al., 1993), which is consistent with their function in reducing protein aggregations. A surprising finding is the significant association of Clusterin with several coagulation pathway-related proteins (ranks shown in parentheses), including: F5 (3), F13B (9), FGG (18), and FGA (27). In addition, it has a strong reciprocal interaction with mitochondrial malic enzyme 2 (ME2, ρ = 0.62, FDR = 3.9E-05, reciprocal ranks 12,2). Analysis of CLU's top 2% strongest ERCs shows significant enrichment for 186 terms. CLU's top 4 most significantly enriched terms all relate to the coagulation cascades and clot formation. Additional significant terms are relevant to immunity, such as "Immune system" (FDR = 4.8E-03), "Signaling by Interleukins" (FDR = 4.1E-03), and "Plasma Cell", an activated immune cell type (FDR = 3.4E-05).

Of direct relevance to COVID-19, Singh et al (2021) found in an expression study of coronavirus infected cells that SARS-CoV-2, SARS-CoV, and MERS-CoV, show shared expression alterations for two genes, one of which is CLU. Therefore, the ERC results for CLU are consistent with aspects of their known function, and their interactions with coronavirus infections.

*GPR141 (G Protein-Coupled Receptor 141)*: Although GPR141 falls just outside the top 1% ACE2 ERC set (rank 24 – 1.2%), its relevance to Clusterin and our protein network analysis below warrants its inclusion here. There is limited information on GPR141 in the literature. Nevertheless,

GPR141 forms a very strong reciprocal rank with CLU (1,1), each being the top interactor with the other, and CLU-GPR141-ACE2 forms a reciprocal rank 24 triad. According to the Human Protein Atlas (Uhlén et al., 2015), it is highly expressed in the brain, bone marrow, lymphatic tissue, and blood. Cell types showing enriched expression of GPR141 include granulocytes, Kupffer cells, and macrophages, as well as alveolar cell types 1 & 2. A recent study found that GPR141 expression is a molecular signature for megakaryocytes (Lu et al., 2018), the progenitor cells for platelets and red blood cells. Noteworthy in this regard is that autopsy results of COVID-19 victims with neurological manifestations find an unusual presence of megakaryocytes in brain capillaries (Nauen et al., 2021). Additionally, elevated levels of IFN-activated megakaryocytes are observed in the blood of patients with severe COVID-19 (Bernardes et al., 2020). These findings suggest possible roles for GPR141 in COVID-19 pathologies.

Although there is limited information on GPR141, its protein interactions revealed by ERCs could be informative. The GPR141's top 2 percent ERCs show significant enrichment for 111 terms (Supplementary File S3). Most of its top enriched terms relate to the coagulation cascade (FDR = 2.9E-10), with many of the contributing proteins being similar to Clusterin's protein set. Additionally, there is significant enrichment for terms related to regulation of vasodilator nitric oxide (FDR = 3.0E-03), ceramide/sphingolipid signaling (FDR = 6.8E-03) and cytokine responses (FDR = 6.8E-03).

Recent studies implicate GPR141 in Alzheimer's Disease (AD) (Srinivasan et al., 2020; Hodges et al., 2021; Novikova et al., 2021). The finding may be noteworthy given the very strong ERC association of GPR141 with CLU and their top reciprocal ranks (1,1). Multiple lines of evidence implicate CLU in AD, including a role in amyloid Aβ processing, CLU polymorphism association with late-onset AD (Balcar et al., 2021), and correlations of CLU levels in serum and cerebrospinal fluid with AD (Shepherd et al., 2020). Since the function of GPR141 is poorly understood, the ERC results suggest that the two proteins interact closely, possibly through physical binding, and their functional relationships should be further explored.

*TMEM63C (Transmembrane Protein 63C)*: TMEM63C is the 4$^{th}$ ranking ACE2 ERC (FDR = 1.3E-04), and the two have strong reciprocal ranks (and ACE2 show a strong reciprocal rank ERCs (3,10), suggestive of direct reciprocal interactions. Along with other family members, TMEM63C forms a membrane channel and functions in osmolarity perception and regulation (Zhao et al., 2016). It plays an important role in kidney function and kidney disease (Schulz et al., 2019), with angiotensin II inducing its expression in glomerular podocyte cells (Eisenreich et al., 2020). Reduced expression of TMEM63C can result in podocyte apoptosis (Eisenreich et al., 2020). The connection between TMEM63C and angiotensin II is a further indication of a functional interaction, given that ACE2 metabolizes angiotensin II to angiotensin (1-7) as part of the RAS pathway. The RAS pathway is implicated in aspects of COVID-19 (Kai & Kai, 2020).

TMEM63C's top 2% ERC list has significant enrichment for three terms related to the coagulation cascade (FDR = 6.8E-04). Tissue enrichment reveals "adult liver" as the most enriched term (FDR = 8.0E-03). Importantly, there are significant terms related to peptidase activity and the Renin-angiotensin system (driven by the proteins ACE2 and ANPEP). ANPEP is particularly interesting as it has been previously identified as a receptor for several coronaviruses such as HCV-229E (Yeager et al., 1992). ANPEP is known to be a metallopeptidase (as is ACE2) and has been implicated in the regulation of angiogenesis (Rangel et al., 2007). Additionally, ANPEP is known to have Angiotensin III as a substrate (Danziger, 2008), tying it back to the RAS pathway, with

ACE2 and TMEM63C. Therefore, the ACE2-TMEM63C reciprocal rank ERCs may indicate direct biological interactions between the proteins, possibly involving physical binding.

*IFNAR2 (Interferon alpha/beta receptor 2)*: IFNAR2 is the 5th ranking ACE2 ERC, with highly significant correlation (ρ = 0.62, FDR = 6.1E-04). IFNAR2 combines with IFNAR1 to form the IFN-alpha/beta receptor, which acts through JAK/STAT signaling to modulate immune responses. IFNAR1/IFNAR2 is the receptor for both alpha and beta interferons and is involved in immune responses to viral infection, most notably to influenza and defense against bacterial infections (Shepardson et al., 2018). IFNAR2 was not originally in our protein set, but we added it based on a paper that implicated this protein in severe COVID-19 based on GWAS and gene expression changes (Liu et al., 2021; Pairo-Castineira et al., 2021). Another study implicates mutations in IFNAR2 with severe COVID-19 (Zhang et al., 2020). When added to our ERC protein set, it was found to be a high ERC to ACE2 (rank 5 in the ACE2 set), providing independent support for its role in COVID-19, possibly through direct ACE2-IFNAR2 interactions.

There are both soluble and membrane-bound forms of IFNAR2. The soluble form (sIIFNAR2) "exerts immunomodulatory, antiproliferative and antiviral activities" (Hurtado-Guerrero et al., 2020). The presence of soluble forms for both IFNAR2 and ACE2 suggests possible avenues for physical interaction, in addition to between their membrane-bound forms. IFNAR2 and IFNAR1 combine to form the IFN-alpha/beta receptor, and as expected, these two proteins are significantly and highly correlated (ρ = 0.79, FDR = 1.9E-09, reciprocal ranks 19,2). CD40, which ranks IFNAR2 as its top ERC, is a crucial immunity protein in the tumor necrosis factor-R (TNF-R) family, with roles in B lymphocytes, macrophages, and cytotoxic T lymphocytes (Grewal & Flavell, 1998; Van Kooten & Banchereau, 2000). IFNAR2 has eleven proteins showing RR20, which is discussed further in the analysis of reciprocal rank networks (Section D). Enrichment analysis for IFNAR2's top 2% ERCs has an expected strong enrichment for terms related to canonical IFNAR2-related pathways such as "Cytokine-cytokine receptor interaction" (FDR = 1.4E-04), "PI3K-Akt Signaling pathway" (FDR = 1.8E-03), and "JAK-STAT signaling pathway" (FDR = 4.0E-03). Some additional enriched terms of note include several terms related to: tumor necrosis factor signaling, coagulation and complement cascade, ECM receptor interaction/collagen function, and plasma membrane (Supplementary File S3).

*KIF3B (Kinesin Family Member 3B)*: KIF3B is the 6th highest ACE2 ERC. This protein is involved in chromosomal segregation during meiosis and mitosis and also participates in intracellular trafficking (Stelzer et al., 2016). Along with GEN1, it is another high-ranking ACE2 ERC involved in chromosomal processes. Among its phenotypes are ciliary assembly (Cogné et al., 2020), endocytosis (Reed et al., 2010), and regulation of dendrite structure in neurons (Joseph et al., 2020). KIF3B's top ERC is Secretogranin II (SCG2), which is a neuroendocrine protein that regulates the formation of secretory granules (Stelzer et al., 2016). Genetic variants of its 2nd ranking ERC, Inositol hexakisphosphate kinase 3 (IP6K3) are associated with Alzheimer's disease (Crocco et al., 2016) and its 4th ranking protein, Neuronal Pentraxin Receptor (NPTXR), with which it has strong reciprocal ranks (4,6), is a biomarker for Alzheimer's disease (Lim et al., 2020). The nature of KIF3Bs interactions with ACE2 is not immediately obvious, except for a possible functional connection between ACE2 at amyloid protein catalysis (Kehoe, 2018; Evans et al., 2020). KIF3B top 2% ERCs show significant enrichment only for the "Complement and coagulation cascades" term from KEGG (FDR = 1.9E-02).

*ITPRIPL2 (Inositol 1,4,5-Trisphosphate Receptor Interacting Protein-Like 2)*: ITPRIPL2 is the 7th highest among ACE2's ERC set. Information about this protein is limited in the literature. It is reported in the Human Protein Atlas to be localized to centrosomes. Examination of its ERC set could provide some information relevant to studies of this protein and possible interactions with ACE2. Among its highest ranking ERCs are two proteins associated with DNA repair and mitotic processes. FANCG (1) is involved with double-strand break repair (Yamamoto et al., 2003). CC2D1B plays a role in the reformation of the mitotic nuclear envelope (Vietri & Stenmark, 2018), has a high reciprocal rank association with ITPRIPL2 (2,6). In turn, CCD1B has high reciprocal ranks with GEN1 (2,1), which is involved in holiday junction resolution and genomic stability (see description above). These findings are consistent with the centrosome localization of ITPRIPL2 and suggest that these proteins may physically interact in a manner that results in correlated protein evolution. Three other proteins showing reciprocal rank associations (RR10) are CC2D1B (2, 6), ENAM (4,5), and STAT6 (10,9). Why ACE2 shows a high ERC with ITPRIPL2 is unclear. An ITPRIPL2 top 2% ERC enrichment analysis indicates cytokine receptor activity (FDR = 1.6E-02) and tumor necrosis factor signaling terms (FDR = 2.4E-02). Additionally, there is significant enrichment for “DNA metabolic process” (FDR = 4.9E-02).

*FAM227A (Family with Sequence Similarity 227 Member A)*: FAM227A is the 8$^{th}$ ranking ACE2 ERC. There is little information about this protein in the current literature, so its evolutionary protein correlations could be informative. The Human Protein Atlas indicates that gene expression is enhanced in the pituitary gland and testes, in ciliated cells, early and late spermatids, and cone & rod photoreceptors (Uhlén et al., 2015). The top five ERC proteins for FAM227A are F5 (involved in blood coagulation), SPZ1 (enriched in spermatids), C16orf96 (enriched in spermatids), FSCB (enriched in spermatids ), and FERIL5 (enriched in spermatids) (Uhlén et al., 2015). This ERC pattern strongly suggests functional interactions among these proteins in spermatogenesis. Moreover, ACE2 is expressed in spermatogonia (Wang & Xu, 2020) and is implicated in male fertility issues associated with COVID-19 (Liu et al., 2020; Verma, Saksena & Sadri-Ardekani, 2020). Therefore, we suggest that this effect could be mediated by FAM227A, a possibility that is worth further exploration. The top 2% of FAM227A ERCs are enriched for 40 terms and reveal a strong association with inflammatory signaling/immunity (Supplementary File S3). In particular, the most significant enrichment is the KEGG term “Cytokine-cytokine receptor interaction” (FDR = 1.3E-04). Most of the proteins driving enrichment for such terms are toll-like receptors, interferon/interleukin receptors, and cytokine receptors.

*TLR8 (Toll-like Receptor 8)*: TLR8 is the 9$^{th}$ ranking ACE2 ERC. Toll-like receptors are a class of proteins that can detect and initiate an innate immune response to foreign invaders (Takeda, Kaisho & Akira, 2003) by recognizing conserved features of pathogens (Kawai & Akira, 2010). Importantly, toll-like receptor responses are usually associated with large inflammatory responses of the immune system (Takeda, Kaisho & Akira, 2003; Kawai & Akira, 2010). TLR8 has strong ERCs to several other toll-like receptors such as TLR9 (ranks 11, 13) and a unidirectional connection to TLR7 (rank 26, $\rho = 0.71$, FDR = 9.6E-08). Consistent with these observations, enrichment of the top 2% ERC list of TLR8 shows highly significant terms associated with TLR8 such as TRAF6 mediated IRF7 activation in TLR7/8 or 9 signaling (FDR = 8.3E-07) and the toll-like receptor signaling pathway (FDR = 2.1E-06). Additionally, the other significantly enriched terms are overwhelmingly related to other immunity-related pathways (Supplementary File S3).

*COL4A4 (Collagen Type IV Alpha 4)*: COL4A4 is the 10$^{th}$ ranking ACE2 ERC. Collagen Type 4 is a complex of six proteins that are part of the extracellular matrix called the basement membrane, which resides between epithelial cells (Stelzer et al., 2016), such as those of glomerulus and capillaries. Type 4 collagen is a major constituent of glomerular basement membranes. Mutations in COL4A4 and other COL4A genes are associated with inherited kidney disease such as Alport syndrome (Buzza et al., 2001) and familial hematuria (Longo et al., 2002). Top 2% ERC list enrichment analysis shows significant enrichment for immunity signaling related terms such as Cytokine-cytokine receptor interaction (FDR = 1.7E-04), PI3k-Akt signaling pathway (FDR = 3.0E-03; of which type IV collagen subunits are canonically annotated as a part of), and JAK-STAT signaling pathway (FDR p = 7.0E-03).

*FAM3D (FAM3 Metabolism Regulating Signaling Molecule D)*: FAM3D is the 11$^{th}$ ranking ACE2 ERC. As seen in figure ACE2-RRN Net, FAM3D is one of four proteins with strong reciprocal rank correlations to ACE2. It is a chemoattractant for neutrophils and monocytes in peripheral blood, is implicated in inflammatory responses in the gastrointestinal tract (Peng et al., 2016). Studies indicate that it has a role in nutritional regulation in the gastrointestinal tract (de Wit et al., 2012), and this may provide a functional connection, given the role of ACE2 in the processing of peptides in the gut (Kuba et al., 2010). Strikingly, ACE2 and FAM3D show strong ERC reciprocal ranks and form a RR network with CLU and GPR141. It also shows strong RR with Solute Carrier Family 16 Member 11 (SLC16A11). Several coagulation cascade proteins are present in its top 1% interaction set, including F13B (its highest-ranked ERC), SERPINA5, and FGB, suggesting possible links to coagulation pathologies of COVID-19. The top 2%ERC list enrichment analysis results in the top 5 terms related to coagulation and clotting (FDR = 3.5E-09). Additionally, there is strong enrichment for various immune response-related terms such as “cytokine receptor activity” (FDR = 2.2E-03) and enrichment for plasma cell presence (FDR = 5.0E-03).

*F5 (Coagulation Factor 5, also abbreviated FV)*: F5 is the 12$^{th}$ ranking ACE2 ERC. F5 is a key regulator of hemostasis and a central cofactor involved in blood coagulation (Ivanciu et al., 2017). Our ERC analysis predicts strong interactions between ACE2 and F5 (rank 12 for ACE2, $\rho$ = 0.57, FDR = 7.2E-04), possibly mediated through the Clusterin (see below). F5 can act as a cofactor for coagulation or anticoagulation (Cramer & Gale, 2012). Approximately 20% of circulating F5 resides in platelets with the remainder in plasma (Gould, Silveira & Tracy, 2004), and whereas plasma F5 has an important role in thrombin formation in microcirculation, platelet F5 has a larger role in severe injury (Ivanciu et al., 2017). The former role could be relevant to micro thrombosis observed in COVID-19. In fact, F5 has been found to associate with COVID-19 symptom severity (elevation in F5 activity) and this may be due to the high abundance of megakaryocytes in the lungs and hearts in COVID-19 infected patients (Stefely et al., 2020). This is further supported by a gene set overlap study showing F5 being annotated to all five examined comorbidities linked to COVID-19 severity (Dolan et al., 2020).

Our ERC analysis of F5 suggests that it may have many other functions beyond the coagulation pathway. F5 is a very “connected” protein with strikingly strong ERC correlations. Twenty-one proteins have spearman rank correlations > 0.80. In addition, seven proteins rank F5 first among their ERCs and 43 rank F5 in their top 5 ERCs. The strongest enrichments of the top 2% ERCs are immune response-related terms such as “response to cytokine” (FDR = 1.1E-03) and “inflammatory response” (FDR = 1.2E-03). Notably, there is only one significant coagulation-related term in this list, “Complement and Coagulation Cascades” (FDR = 6.9E-03)

*AR (Androgen Receptor)*: AR is the 13th ranked ACE2 ERC ($\rho$ = 0.52, FDR = 8.8E-04) and is barely cut off from the RR20 criteria to ACE2 (the rank of ACE2 is 22nd in the AR ERC list). AR is encoded on the X chromosomes and is a hormonal receptor that plays a major role in male development, particularly in male reproductive systems and somatic differentiation (Matsumoto et al., 2008). It. AR’s top-ranking ERC is spermatogenesis associated 25 protein (SPATA25) with (1,2) reciprocal ranks, and its top 2% ERCs only show significant enrichment for cytokine receptor activity (FDR = 1.1E-03). In addition to its roles in sexual differentiation and behavior (Cunningham, Lumia & McGinnis, 2012), AR enhances prostate cancer cell growth (Gelmann, 2002). It may play a role in microbial infection resistance as a knockout in mice can reduce the development and proliferation of neutrophils (Chuang et al., 2009). Androgen signaling may play a role in SARs-CoV-2 infectivity, as indicated by knockdowns of AR in prostate cells result in downregulation of ACE2 and infection cofactors TMPRSS2 and FURIN (Samuel et al., 2020). Additionally, AR has been annotated as being associated with 4 of the 5 COVID comorbidities that are associated with COVID severity in Dolan et al (2020). Male fertility problems may be associated with COVID-19 infection and the ACE2 receptor is abundant in male genetical track and spermatagonia (Huang et al., 2021; Seymen, 2021). ACE2-AR protein interactions, as predicted by ERC, may play a role in these pathologies.

*TSGA13 (Testis specific gene 13 protein)*: TSGA13 is the 14$^{th}$ ranking ERC for ACE2 ($\rho = 0.57$, FDR = 8.8E-04). The function of this protein is not well understood, so it is characterized by its expression in the testes (Zhao et al., 2015). Despite its high expression in the testes, TSGA13 is expressed in other tissues (Zhao et al., 2015) and it may not play a role in fertility as mice with TSGA13 knocked out were still fertile (Miyata et al., 2016). However, this protein is highly conserved (Zhao et al., 2015) so may still play an important role in organisms. TSGA13 variation has been associated with total colonic aganglionosis in patients with Hirschsprung disease (Jung et al., 2019) and reduced expression of TSGA13 has been associated with carcinoma (Zhao et al., 2015). We, therefore, propose that ERC analysis can provide insight into the potential function of TSGA13 as it has many extraordinarily high ERCs (78 proteins show $\rho$ values of 0.7 or higher). The top ERC is C16orf96 ($\rho = 0.83$, FDR = 4.5E-12) which is not well understood, but its 2$^{nd}$ highest ERC is C3orf30 ($\rho = 0.82$, FDR = 2E-10), also known as "testis expressed 55" (TEX55) which may play a role in fertility, especially considering its strong expression in adult testes (Jamin et al., 2021). The ERC results coupled with known expression profiles suggest that TSGA13 and C3orf30 may interact with each other, although there is no external evidence to suggest this currently. Furthermore, TSGA13's potential interaction with ACE2 may be mediated through their common ERC partners such as F5 ($\rho = 0.80$, FDR = 3.1E-11), TLR8 ($\rho = 0.78$, FDR = 5.7E-10), and IFNAR2 ($\rho = 0.75$, FDR = 6.1E-09). The top 2% ERCs show enrichment for many immunity/interferon-related terms (FDR = 7.0E-05), complement and coagulation cascade (FDR = 1.4E-04), and no terms related to male fertility or male reproductive tissues.

*PLA2G7 (Platelet-activating factor acetylhydrolase)*: PLA2G7 is the 15$^{th}$ ranking ERC for ACE2 ($\rho = 0.57$, FDR = 8.4E-04). PLA2G7 is a member of the arachidonic acid pathway and is potentially associated with prostate cancer (Vainio et al., 2011). PLA2G7's strong ERC to ACE2 is particularly interesting due to its likely association with cardiovascular and heart disease (Sutton et al., 2008; Wang et al., 2010), each of which are associated with COVID-19 (Bansal, 2020; Alsaied et al., 2020). Additionally, PLA2G7's role in the arachidonic acid pathway is relevant to COVID-19 pathologies as a deficiency in arachidonic acid may lead to greater COVID-19 susceptibility and the arachidonic acid pathway is a candidate therapeutic target (Hoxha, 2020; Ripon et al., 2021). The connection to ACE2 specifically may also make biological sense as MAS (the receptor for the Angiotensin(1-7) that ACE2 can produce) can cause the release of arachidonic acid (Bader, 2013). Analysis of PLA2G7's top 2% ERC list shows significant enrichment for various terms related to immunity such as "cytokine receptor activity" (FDR = 1.9E-05) and several viral infection pathways such as Influenza A infection (FDR = 6.5E-03). Interestingly, there was also significant enrichment for terms related to DNA repair (FDR = 4.3E-02).

*MMS19 (MMS19 nucleotide excision repair homolog)*: MMS19 is the 16th ranking ERC for ACE2 (ρ = 0.56, FDR = 8.9E-04). Like ACE2's strongest ERC partner, GEN1, MMS19 is involved in DNA repair (Stehling et al., 2012). It is also specifically associated with the "cytosolic Fe-S protein assembly (CIA)", which forms a complex with MMS19 to assist in DNA metabolism, replication, and repair (Gari et al., 2012). Similar to GEN1, MMS19's mode of interaction with ACE2 is still unclear. But the top ERCs of MMS19 show several proteins directly related to DNA maintenance such as POLL (DNA polymerase lambda; ρ = 0.76, FDR = 7.2E-10) and GEN1 (ρ = 0.74, FDR = 6.2E-09). But significant enrichment on the top 2% ERC list is just shown for "death receptor activity" (FDR = 3.1E-02) and "tumor necrosis factor-activated receptor activity" (FDR = 3.1E-02).

*Angiomotin (AMOT)*: AMOT is the 17th ranking ERC for ACE2 (ρ = 0.56, FDR = 8.8E-04). Its potential relevance to COVID-19 pathologies is clear as AMOT is associated with angiogenesis and endothelial cell movement (Bratt et al., 2005; Aase et al., 2007). These associations may explain its ERC to ACE2 as well. For instance, ACE2 can promote endothelial cell migration (Jin et al., 2015). Additionally, COVID-19 infection has been associated with angiogenesis in the lungs (Ackermann et al., 2020). AMOT shares several of ACE2's top ERCs. For instance, GEN1 and TSGA13 are both among AMOT's top 20 ERCs. The top 2% ERCs of AMOT show significant enrichment for complement and coagulation cascades (FDR = 4.3E-04), inflammatory response (FDR = 1.0E-03), and spermatogenesis (FDR = 2.7E-02).

*L1CAM (L1 cell adhesion molecule)*: L1CAM is a RR20 protein to ACE2 (ρ = 0.56, FDR = 8.8E-04, ranks 18, 14). It is a part of the immunoglobulin superfamily and is best characterized for its role in the nervous system, specifically relating to the development of the brain (Schäfer & Altevogt, 2010). Interestingly, L1CAM is embedded in the extracellular membrane but can be cleaved near the membrane to allow for the circulation of the truncated protein (Schäfer & Altevogt, 2010). The metallopeptidase ADAM17 is one of the enzymes that cleaves L1CAM near the membrane (Schäfer & Altevogt, 2010), and is also known to mediate the release of the ectodomain of ACE2 from the extracellular membrane as well (Lambert et al., 2005). Thus, both proteins circulate in plasma where they may interact, although the functional basis of this postulated interaction is unclear. L1CAM has three other RR20 proteins: BMX non-receptor tyrosine kinase (BMX; ranks 1,3), cyclin-dependent kinase inhibitor 2C (CDKN2C ranks 2,20), and glycerophosphodiester phosphodiesterase domain containing 3 (GDPD3, 5,19). The top 2% enrichment for L1CAM has several significant terms for complement and coagulation cascades (FDR = 5.4E-04), positive regulation of cellular protein localization (FDR = 5.2E-03), endopeptidase activity (FDR = 5.9E-03), Alzheimer's Disease (FDR = 1.1E-02), and arachnoid cyst (FDR = 3.6E-04). It is possible, although highly speculative, that ACE2-L1CAM protein interactions could play a role in neurological pathologies associated with COVID-19.

*PDYN (Prodynorphin aka Leumorphin)*: PDYN is the 19th ranking ERC for ACE2 (ρ = 0.56, FDR = 8.8E-04). PDYN is an endogenous opioid receptor (Stelzer et al., 2016), which also inhibits vasopressin secretion (Yamada et al., 1988), suggesting a connection to ACE2 in blood pressure homeostasis. Unsurprisingly, PDYN is implicated in neurotransmission and mental disorders (such as schizophrenia, Alzheimer's, epilepsy, and cerebellar ataxia) (Clarke et al., 2012; Jezierska et al., 2013; Henriksson et al., 2014). PDYN has several proteins involved in immune function among its top ERCs such as Interferon lambda receptor 1 (IFNLR1; ρ = 0.77, FDR = 6.4E-10) and Toll-like receptor 7 (TLR7; ρ = 0.75, FDR = 2.4E-09). The top 2% ERC list of PDYN shows significant enrichment for terms related to immune system function (FDR = 6.0E-03), the complement and coagulation cascades (FDR = 6.0E-03), but no significant terms related to brain function other than “NCAM1 interactions” (FDR= 4.9E-02).

*IQ motif containing D (IQCD)*: IQCD is the 20th ranking ERC for ACE2 (ρ = 0.56, FDR = 8.9E-04). IQCD in mammals is not well studied. But it has been characterized as being involved in the “acrosome” (Zhang et al., 2019). The acrosome is an organelle that is part of the sperm and is involved in the “acrosome reaction”, which allows sperm to fuse with an egg upon fertilization (Abou-Haila & Tulsiani, 2000). It is required for spermatogenesis in mice (Harris et al., 2014) IQCD is therefore another protein with strong ERC to ACE2 implicated in male sex organs. There is also some evidence that suggests IQCD is associated with male fertility (Zhang et al., 2019). Additionally, ACE2 presence may be negatively associated with the acrosome reaction in sperm-precursor cells (Wang & Xu, 2020), but the direct mechanism for this is unclear. The top 2% ERC list for IQCD shows enrichment for tumor-necrosis factor-related terms (FDR = 9.3E-04) and “SW-620 cell” (4.9E-02) which is a human colon carcinoma cell line.

### I. Supplementary Text References


Aase K, Ernkvist M, Ebarasi L, Jakobsson L, Majumdar A, Yi C, Birot O, Ming Y, Kvanta A, Edholm D, Aspenström P, Kissil J, Claesson-Welsh L, Shimono A, Holmgren L. 2007. Angiomotin regulates endothelial cell migration during embryonic angiogenesis. *Genes and Development* 21:2055–2068. DOI: 10.1101/gad.432007.

Abou-Haila A, Tulsiani DRP. 2000. Mammalian sperm acrosome: Formation, contents, and function. *Archives of Biochemistry and Biophysics* 379:173–182. DOI: 10.1006/abbi.2000.1880.

Ackermann M, Verleden SE, Kuehnel M, Haverich A, Welte T, Laenger F, Vanstapel A, Werlein C, Stark H, Tzankov A, Li WW, Li VW, Mentzer SJ, Jonigk D. 2020. Pulmonary Vascular Endothelialitis, Thrombosis, and Angiogenesis in Covid-19. *New England Journal of Medicine* 383:120–128. DOI: 10.1056/nejmoa2015432.

Alsaied T, Aboulhosn JA, Cotts TB, Daniels CJ, Etheridge SP, Feltes TF, Gurvitz MZ, Lewin MB, Oster ME, Saidi A. 2020. Coronavirus Disease 2019 (COVID-19) Pandemic Implications in Pediatric and Adult Congenital Heart Disease. *Journal of the American Heart Association* 9:e017224. DOI: 10.1161/JAHA.120.017224.

Bader M. 2013. ACE2, angiotensin-(1-7), and Mas: The other side of the coin. *Pflugers Archiv European Journal of Physiology* 465:79–85. DOI: 10.1007/s00424-012-1120-0.

Balcar VJ, Zeman T, Janout V, Janoutová J, Lochman J, Šerý O. 2021. Single Nucleotide Polymorphism rs11136000 of CLU Gene (Clusterin, ApoJ) and the Risk of Late-Onset Alzheimer's Disease in a Central European Population. *Neurochemical Research* 46:411–422. DOI: 10.1007/s11064-020-03176-y.

Bansal M. 2020. Cardiovascular disease and COVID-19. *Diabetes and Metabolic Syndrome: Clinical Research and Reviews* 14:247–250. DOI: 10.1016/j.dsx.2020.03.013.

Bateman A, Martin MJ, Orchard S, Magrane M, Agivetova R, Ahmad S, Alpi E, Bowler-Barnett EH, Britto R, Bursteinas B, Bye-A-Jee H, Coetzee R, Cukura A, Silva A Da, Denny P, Dogan T, Ebenezer TG, Fan J, Castro LG, Garmiri P, Georghiou G, Gonzales L, Hatton-Ellis E, Hussein A, Ignatchenko A, Insana G, Ishtiaq R, Jokinen P, Joshi V, Jyothi D, Lock A, Lopez R, Luciani A, Luo J, Lussi Y, MacDougall A, Madeira F, Mahmoudy M, Menchi M, Mishra A, Moulang K, Nightingale A, Oliveira CS, Pundir S, Qi G, Raj S, Rice D, Lopez MR, Saidi R, Sampson J, Sawford T, Speretta E, Turner E, Tyagi N, Vasudev P, Volynkin V, Warner K, Watkins X, Zaru R, Zellner H, Bridge A, Poux S, Redaschi N, Aimo L, Argoud-Puy G, Auchincloss A, Axelsen K, Bansal P, Baratin D, Blatter MC, Bolleman J, Boutet E, Breuza L, Casals-Casas C, de Castro E, Echioukh KC, Coudert E, Cuche B, Doche M, Dornevil D, Estreicher A, Famiglietti ML, Feuermann M, Gasteiger E, Gehant S, Gerritsen V, Gos A, Gruaz-Gumowski N, Hinz U, Hulo C, Hyka-Nouspikel N, Jungo F, Keller G, Kerhornou A, Lara V, Le Mercier P, Lieberherr D, Lombardot T, Martin X, Masson P, Morgat A, Neto TB, Paesano S, Pedruzzi I, Pilbout S, Pourcel L, Pozzato M, Pruess M, Rivoire C, Sigrist C, Sonesson K, Stutz A, Sundaram S, Tognolli M, Verbregue L, Wu CH, Arighi CN, Arminski L, Chen C, Chen Y, Garavelli JS, Huang H, Laiho K, McGarvey P, Natale DA, Ross K, Vinayaka CR, Wang Q, Wang Y, Yeh LS, Zhang J. 2021. UniProt: The universal protein knowledgebase in 2021. *Nucleic Acids Research* 49:D480–D489. DOI: 10.1093/nar/gkaa1100.

Bernardes JP, Mishra N, Tran F, Bahmer T, Best L, Blase JI, Bordoni D, Franzenburg J, Geisen U, Josephs-Spaulding J, Köhler P, Künstner A, Rosati E, Aschenbrenner AC, Bacher P, Baran N, Boysen T, Brandt B, Bruse N, Dörr J, Dräger A, Elke G, Ellinghaus D, Fischer J, Forster M, Franke A, Franzenburg S, Frey N, Friedrichs A, Fuß J, Glück A, Hamm J, Hinrichsen F, Hoeppner MP, Imm S, Junker R, Kaiser S, Kan YH, Knoll R, Lange C, Laue G, Lier C, Lindner M, Marinos G, Markewitz R, Nattermann J, Noth R, Pickkers P, Rabe KF, Renz A, Röcken C, Rupp J, Schaffarzyk A, Scheffold A, Schulte-Schrepping J, Schunk D, Skowasch D, Ulas T, Wandinger KP, Wittig M, Zimmermann J, Busch H, Hoyer BF, Kaleta C, Heyckendorf J, Kox M, Rybniker J, Schreiber S, Schultze JL, Rosenstiel P, Banovich NE, Desai T, Eickelberg O, Haniffa M, Horvath P, Kropski JA, Lafyatis R, Lundeberg J, Meyer K, Nawijn MC, Nikolic M, Ordovas Montanes J, Pe'er D, Tata PR, Rawlins E, Regev A, Reyfman P, Samakovlis C, Schultze J, Shalek A, Shepherd D, Spence J, Teichmann S, Theis F, Tsankov A, van den Berge M, von Papen M, Whitsett J, Zaragosi LE, Angelov A, Bals R, Bartholomäus A, Becker A, Bezdan D, Bonifacio E, Bork P, Clavel T, Colme-Tatche M, Diefenbach A, Dilthey A, Fischer N, Förstner K, Frick JS, Gagneur J, Goesmann A, Hain T, Hummel M, Janssen S, Kalinowski J, Kallies R, Kehr B, Keller A, Kim-Hellmuth S, Klein C, Kohlbacher O, Korbel JO, Kurth I, Landthaler M, Li Y, Ludwig K, Makarewicz O, Marz M, McHardy A, Mertes C, Nöthen M, Nürnberg P, Ohler U, Ossowski S, Overmann J, Peter S, Pfeffer K, Poetsch AR, Pühler A, Rajewsky N, Ralser M, Rieß O, Ripke S, Nunes da Rocha U, Saliba AE, Sander LE, Sawitzki B, Schiffer P, Schulte EC, Sczyrba A, Stegle O, Stoye J, Vehreschild J, Vogel J, von Kleist M, Walker A, Walter J, Wieczorek D, Ziebuhr J. 2020. Longitudinal Multi-omics Analyses Identify Responses of Megakaryocytes, Erythroid Cells, and Plasmablasts as Hallmarks of Severe COVID-19. *Immunity* 53:1296-1314.e9. DOI: 10.1016/j.immuni.2020.11.017.

Bratt A, Birot O, Sinha I, Veitonmäki N, Aase K, Ernkvist M, Holmgren L. 2005. Angiomotin regulates endothelial cell-cell junctions and cell motility. *Journal of Biological Chemistry* 280:34859–34869. DOI: 10.1074/jbc.M503915200.

Buzza M, Wang YY, Dagher H, Babon JJ, Cotton RG, Powell H, Dowling J, Savige J. 2001. COL4A4 mutation in thin basement membrane disease previously described in Alport syndrome. *Kidney International* 60:480–483. DOI: 10.1046/j.1523-1755.2001.060002480.x.

Casino P, Gozalbo-Rovira R, Rodríguez-Diáz J, Banerjee S, Boutaud A, Rubio V, Hudson BG, Saus J, Cervera J, Marina A. 2018. Structures of collagen IV globular domains: Insight into associated pathologies, folding and network assembly. *IUCrJ* 5:765–779. DOI: 10.1107/S2052252518012459.

Chan YW, West S. 2015. GEN1 promotes Holliday junction resolution by a coordinated nick and counter-nick mechanism. *Nucleic Acids Research* 43:10882–10892. DOI: 10.1093/nar/gkv1207.

Chuang KH, Altuwaijri S, Li G, Lai JJ, Chu CY, Lai KP, Lin HY, Hsu JW, Keng P, Wu MC, Chang C. 2009. Neutropenia with impaired host defense against microbial infection in mice lacking androgen receptor. *Journal of Experimental Medicine* 206:1181–1199. DOI: 10.1084/jem.20082521.

Clarke TK, Ambrose-Lanci L, Ferraro TN, Berrettini WH, Kampman KM, Dackis CA, Pettinati HM, O'Brien CP, Oslin DW, Lohoff FW. 2012. Genetic association analyses of PDYN polymorphisms with heroin and cocaine addiction. *Genes, Brain and Behavior* 11:415–423. DOI: 10.1111/j.1601-183X.2012.00785.x.

Cogné B, Latypova X, Senaratne LDS, Martin L, Koboldt DC, Kellaris G, Fievet L, Le Meur G, Caldari D, Debray D, Nizon M, Frengen E, Bowne SJ, Buckley RM, Aberdein D, Alves PC, Barsh GS, Bellone RR, Bergström TF, Boyko AR, Brockman JA, Casal ML, Castelhano MG, Distl O, Dodman NH, Ellinwood NM, Fogle JE, Forman OP, Garrick DJ, Ginns EI, Häggström J, Harvey RJ, Hasegawa D, Haase B, Helps CR, Hernandez I, Hytönen MK, Kaukonen M, Kaelin CB, Kosho T, Leclerc E, Lear TL, Leeb T, Li RHL, Lohi H, Longeri M, Magnuson MA, Malik R, Mane SP, Munday JS, Murphy WJ, Pedersen NC, Rothschild MF, Rusbridge C, Shapiro B, Stern JA, Swanson WF, Terio KA, Todhunter RJ, Warren WC, Wilcox EA, Wildschutte JH, Yu Y, Cadena EL, Daiger SP, Bujakowska KM, Pierce EA, Gorin M, Katsanis N, Bézieau S, Petersen-Jones SM, Occelli LM, Lyons LA, Legeai-Mallet L, Sullivan LS, Davis EE, Isidor B. 2020. Mutations in the Kinesin-2 Motor KIF3B Cause an Autosomal-Dominant Ciliopathy. *American Journal of Human Genetics* 106:893–904. DOI: 10.1016/j.ajhg.2020.04.005.

Cramer TJ, Gale AJ. 2012. The anticoagulant function of coagulation factor V. *Thrombosis and Haemostasis* 107:15–21. DOI: 10.1160/TH11-06-0431.

Crocco P, Saiardi A, Wilson MS, Maletta R, Bruni AC, Passarino G, Rose G. 2016. Contribution of polymorphic variation of inositol hexakisphosphate kinase 3 (IP6K3) gene promoter to the susceptibility to late onset Alzheimer's disease. *Biochimica et Biophysica Acta - Molecular Basis of Disease* 1862:1766–1773. DOI: 10.1016/j.bbadis.2016.06.014.

Cunningham RL, Lumia AR, McGinnis MY. 2012. Androgen receptors, sex behavior, and aggression. *Neuroendocrinology* 96:131–140. DOI: 10.1159/000337663.

Danziger RS. 2008. Aminopeptidase N in arterial hypertension. *Heart Failure Reviews* 13:293–298. DOI: 10.1007/s10741-007-9061-y.

Dolan ME, Hill DP, Mukherjee G, McAndrews MS, Chesler EJ, Blake JA. 2020. Investigation of COVID-19 comorbidities reveals genes and pathways coincident with the SARS-CoV-2 viral disease. *Scientific Reports* 10:1–11. DOI: 10.1038/s41598-020-77632-8.

Eisenreich A, Orphal M, Böhme K, Kreutz R. 2020. Tmem63c is a potential pro-survival factor in angiotensin II-treated human podocytes. *Life Sciences* 258. DOI: 10.1016/j.lfs.2020.118175.

Evans CE, Miners JS, Piva G, Willis CL, Heard DM, Kidd EJ, Good MA, Kehoe PG. 2020. ACE2 activation protects against cognitive decline and reduces amyloid pathology in the Tg2576 mouse model of Alzheimer's disease. *Acta Neuropathologica* 139:485–502. DOI: 10.1007/s00401-019-02098-6.

Fricke-Galindo I, Falfán-Valencia R. 2021. Genetics Insight for COVID-19 Susceptibility and Severity: A Review. *Frontiers in Immunology* 12. DOI: 10.3389/fimmu.2021.622176.

Gao M, Rendtlew Danielsen J, Wei LZ, Zhou DP, Xu Q, Li MM, Wang ZQ, Tong WM, Yang YG. 2012. A Novel Role of Human Holliday Junction Resolvase GEN1 in the Maintenance of Centrosome Integrity. *PLoS ONE* 7:e49687. DOI: 10.1371/journal.pone.0049687.

Gari K, Ortiz AML, Borel V, Flynn H, Skehel JM, Boulton SJ. 2012. MMS19 links cytoplasmic iron-sulfur cluster assembly to DNA metabolism. *Science* 337:243–245. DOI: 10.1126/science.1219664.

Garland T, Harvey PH, Ives AR. 1992. Procedures for the analysis of comparative data using phylogenetically independent contrasts. *Systematic Biology* 41:18–32. DOI: 10.1093/sysbio/41.1.18.

Gelmann EP. 2002. Molecular biology of the androgen receptor. *Journal of Clinical Oncology* 20:3001–3015. DOI: 10.1200/JCO.2002.10.018.

Gould WR, Silveira JR, Tracy PB. 2004. Unique in vivo modifications of coagulation factor V produce a physically and functionally distinct platelet-derived cofactor: Characterization of purified platelet-derived factor V/Va. *Journal of Biological Chemistry* 279:2383–2393. DOI: 10.1074/jbc.M308600200.

Grewal IS, Flavell RA. 1998. CD40 and CD154 in cell-mediated immunity. *Annual Review of Immunology* 16:111–135. DOI: 10.1146/annurev.immunol.16.1.111.

Harris TP, Schimenti KJ, Munroe RJ, Schimenti JC. 2014. IQ motif-containing G (Iqcg) is required for mouse spermiogenesis. *G3: Genes, Genomes, Genetics* 4:367–372. DOI: 10.1534/g3.113.009563.

Henriksson R, Bäckman CM, Harvey BK, Kadyrova H, Bazov I, Shippenberg TS, Bakalkin G. 2014. PDYN, a gene implicated in brain/mental disorders, is targeted by REST in the adult human brain. *Biochimica et Biophysica Acta - Gene Regulatory Mechanisms* 1839:1226–1232. DOI: 10.1016/j.bbagrm.2014.09.001.

Hodges AK, Piers TM, Collier D, Cousins O, Pocock JM. 2021. Pathways linking Alzheimer's disease risk genes expressed highly in microglia. *Neuroimmunology and Neuroinflammation* 2020. DOI: 10.20517/2347-8659.2020.60.

Hoxha M. 2020. What about COVID-19 and arachidonic acid pathway? *European Journal of Clinical Pharmacology* 76:1501–1504. DOI: 10.1007/s00228-020-02941-w.

Huang C, Ji X, Zhou W, Huang Z, Peng X, Fan L, Lin G, Zhu W. 2021. Coronavirus: A possible cause of reduced male fertility. *Andrology* 9:80–87. DOI: 10.1111/andr.12907.

Huerta-Cepas J, Serra F, Bork P. 2016. ETE 3: Reconstruction, Analysis, and Visualization of Phylogenomic Data. *Molecular Biology and Evolution* 33:1635–1638. DOI: 10.1093/molbev/msw046.

Hurtado-Guerrero I, Hernáez B, Pinto-Medel MJ, Calonge E, Rodriguez-Bada JL, Urbaneja P, Alonso A, Mena-Vázquez N, Aliaga P, Issazadeh-Navikas S, Pavia J, Leyva L, Alcamí J, Alcamí A, Fernández Ó, Oliver-Martos B. 2020. Antiviral, Immunomodulatory and Antiproliferative Activities of Recombinant Soluble IFNAR2 without IFN-ß Mediation. *Journal of Clinical Medicine* 9:959. DOI: 10.3390/jcm9040959.

Itakura E, Chiba M, Murata T, Matsuura A. 2020. Heparan sulfate is a clearance receptor for aberrant extracellular proteins. *Journal of Cell Biology* 219. DOI: 10.1083/JCB.201911126.

Ivanciu L, Crosby J, MacLeod AR, Revenko A, Camire RM, Davidson RJ, Monia BP. 2017. Differential role of plasma and platelet-derived factor V in vivo. *Blood* 130:364–364. DOI: 10.1182/BLOOD.V130.SUPPL_1.364.364.

Jamin SP, Petit FG, Demini L, Primig M. 2021. Tex55 encodes a conserved putative A-kinase anchoring protein dispensable for male fertility in the mouse. *Biology of Reproduction* 104:731–733. DOI: 10.1093/biolre/ioab007.

Jenne DE, Tschopp J. 1989. Molecular structure and functional characterization of a human complement cytolysis inhibitor found in blood and seminal plasma: Identity to sulfated glycoprotein 2, a constituent of rat testis fluid. *Proceedings of the National Academy of Sciences of the United States of America* 86:7123–7127. DOI: 10.1073/pnas.86.18.7123.

Jezierska J, Stevanin G, Watanabe H, Fokkens MR, Zagnoli F, Kok J, Goas JY, Bertrand P, Robin C, Brice A, Bakalkin G, Durr A, Verbeek DS. 2013. Identification and characterization of novel PDYN mutations in dominant cerebellar ataxia cases. *Journal of Neurology* 260:1807–1812. DOI: 10.1007/s00415-013-6882-6.

Jin HY, Chen LJ, Zhang ZZ, Xu Y Le, Song B, Xu R, Oudit GY, Gao PJ, Zhu DL, Zhong JC. 2015. Deletion of angiotensin-converting enzyme 2 exacerbates renal inflammation and injury in apolipoprotein E-deficient mice through modulation of the nephrin and TNF-alpha-TNFRSF1A signaling. *Journal of Translational Medicine* 13:1–16. DOI: 10.1186/s12967-015-0616-8.

Joseph NF, Grinman E, Swarnkar S, Puthanveettil S V. 2020. Molecular Motor KIF3B Acts as a Key Regulator of Dendritic Architecture in Cortical Neurons. *Frontiers in Cellular Neuroscience* 14:1–11. DOI: 10.3389/fncel.2020.521199.

Jung SM, Namgoong S, Seo JM, Kim DY, Oh JT, Kim HY, Kim JH. 2019. Potential association between TSGA13 variants and risk of total colonic aganglionosis in Hirschsprung disease. *Gene* 710:240–245. DOI: 10.1016/j.gene.2019.06.007.

Kai H, Kai M. 2020. Interactions of coronaviruses with ACE2, angiotensin II, and RAS inhibitors—lessons from available evidence and insights into COVID-19. *Hypertension Research* 43:648–654. DOI: 10.1038/s41440-020-0455-8.

Kanehisa M, Goto S. 2000. KEGG: Kyoto Encyclopedia of Genes and Genomes. *Nucleic Acids Research* 28:27–30. DOI: 10.1093/nar/28.1.27.

Kawai T, Akira S. 2010. The role of pattern-recognition receptors in innate immunity: Update on toll-like receptors. *Nature Immunology* 11:373–384. DOI: 10.1038/ni.1863.

Kehoe PG. 2018. The coming of age of the angiotensin hypothesis in Alzheimer's disease: Progress toward disease prevention and treatment? *Journal of Alzheimer's Disease* 62:1443–1466. DOI: 10.3233/JAD-171119.

Kembel SW, Cowan PD, Helmus MR, Cornwell WK, Morlon H, Ackerly DD, Blomberg SP, Webb CO. 2010. Picante: R tools for integrating phylogenies and ecology. *Bioinformatics* 26:1463–1464. DOI: 10.1093/bioinformatics/btq166.

Kim S. 2015. ppcor: An R Package for a Fast Calculation to Semi-partial Correlation Coefficients. *Communications for Statistical Applications and Methods* 22:665–674. DOI: 10.5351/csam.2015.22.6.665.

Van Kooten G, Banchereau J. 2000. CD40-CD40 ligand. *Journal of Leukocyte Biology* 67:2–17. DOI: 10.1002/jlb.67.1.2.

Kriventseva E V., Kuznetsov D, Tegenfeldt F, Manni M, Dias R, Simão FA, Zdobnov EM. 2019. OrthoDB v10: Sampling the diversity of animal, plant, fungal, protist, bacterial and viral genomes for evolutionary and functional annotations of orthologs. *Nucleic Acids Research* 47:D807–D811. DOI: 10.1093/nar/gky1053.

Kuba K, Imai Y, Ohto-Nakanishi T, Penninger JM. 2010. Trilogy of ACE2: A peptidase in the renin-angiotensin system, a SARS receptor, and a partner for amino acid transporters. *Pharmacology and Therapeutics* 128:119–128. DOI:

10.1016/j.pharmthera.2010.06.003.

Kumar S, Stecher G, Suleski M, Hedges SB. 2017. TimeTree: A Resource for Timelines, Timetrees, and Divergence Times. *Molecular biology and evolution* 34:1812–1819. DOI: 10.1093/molbev/msx116.

Lambert DW, Yarski M, Warner FJ, Thornhill P, Parkin ET, Smith AI, Hooper NM, Turner AJ. 2005. Tumor necrosis factor-α convertase (ADAM17) mediates regulated ectodomain shedding of the severe-acute respiratory syndrome-coronavirus (SARS-CoV) receptor, angiotensin-converting enzyme-2 (ACE2). *Journal of Biological Chemistry* 280:30113–30119. DOI: 10.1074/jbc.M505111200.

Lei Y, Takahama Y. 2012. XCL1 and XCR1 in the immune system. *Microbes and Infection* 14:262–267. DOI: 10.1016/j.micinf.2011.10.003.

Letunic I, Bork P. 2021. Interactive Tree Of Life (iTOL) v5: an online tool for phylogenetic tree display and annotation. *Nucleic Acids Research*. DOI: 10.1093/nar/gkab301.

Lim B, Sando SB, Grøntvedt GR, Bråthen G, Diamandis EP. 2020. Cerebrospinal fluid neuronal pentraxin receptor as a biomarker of long-term progression of Alzheimer's disease: a 24-month follow-up study. *Neurobiology of Aging* 93:97.e1-97.e7. DOI: 10.1016/j.neurobiolaging.2020.03.013.

Liu X, Chen Y, Tang W, Zhang L, Chen W, Yan Z, Yuan P, Yang M, Kong S, Yan L, Qiao J. 2020. Single-cell transcriptome analysis of the novel coronavirus (SARS-CoV-2) associated gene ACE2 expression in normal and non-obstructive azoospermia (NOA) human male testes. *Science China Life Sciences* 63:1006–1015. DOI: 10.1007/s11427-020-1705-0.

Liu D, Yang J, Feng B, Lu W, Zhao C, Li L. 2021. Mendelian randomization analysis identified genes pleiotropically associated with the risk and prognosis of COVID-19. *Journal of Infection* 82:126–132. DOI: 10.1016/j.jinf.2020.11.031.

Longo I, Porcedda P, Mari F, Giachino D, Meloni I, Deplano C, Brusco A, Bosio M, Massella L, Lavoratti G, Roccatello D, Frascá G, Mazzucco G, Onetti Muda A, Conti M, Fasciolo F, Arrondel C, Heidet L, Renieri A, De Marchi M. 2002. COL4A3/COL4A4 mutations: From familial hematuria to autosomal-dominant or recessive Alport syndrome. *Kidney International* 61:1947–1956. DOI: 10.1046/j.1523-1755.2002.00379.x.

Lu YC, Xavier-Ferrucio J, Wang L, Zhang PX, Grimes HL, Venkatasubramanian M, Chetal K, Aronow B, Salomonis N, Krause DS. 2018. The Molecular Signature of Megakaryocyte-Erythroid Progenitors Reveals a Role for the Cell Cycle in Fate Specification. *Cell Reports* 25:2083-2093.e4. DOI: 10.1016/j.celrep.2018.10.084.

Maglott D, Ostell J, Pruitt KD, Tatusova T. 2005. Entrez Gene: Gene-centered information at NCBI. *Nucleic Acids Research* 33:D54–D58. DOI: 10.1093/nar/gki031.

Matsumoto T, Shiina H, Kawano H, Sato T, Kato S. 2008. Androgen receptor functions in male and female physiology. *Journal of Steroid Biochemistry and Molecular Biology* 109:236–241. DOI: 10.1016/j.jsbmb.2008.03.023.

McKechnie JL, Blish CA. 2020. The Innate Immune System: Fighting on the Front Lines or Fanning the Flames of COVID-19? *Cell Host and Microbe* 27:863–869. DOI: 10.1016/j.chom.2020.05.009.

Miyata H, Castaneda JM, Fujihara Y, Yu Z, Archambeault DR, Isotani A, Kiyozumi D, Kriseman ML, Mashiko D, Matsumura T, Matzuk RM, Mori M, Noda T, Oji A, Okabe M, Prunskaite-Hyyrylainen R, Ramirez-Solis R, Satouh Y, Zhang Q, Ikawa M, Matzuk MM. 2016. Genome engineering uncovers 54 evolutionarily conserved and testis-enriched genes that are not required for male fertility in mice. *Proceedings of the National Academy of Sciences of the United States of America* 113:7704–7710. DOI: 10.1073/pnas.1608458113.

Mosesson MW. 2005. Fibrinogen and fibrin structure and functions. In: *Journal of Thrombosis and Haemostasis*. John Wiley & Sons, Ltd, 1894–1904. DOI: 10.1111/j.1538-7836.2005.01365.x.

Nauen DW, Hooper JE, Stewart CM, Solomon IH. 2021. Assessing Brain Capillaries in Coronavirus Disease 2019. *JAMA Neurology*. DOI: 10.1001/jamaneurol.2021.0225.

Novikova G, Kapoor M, Tcw J, Abud EM, Efthymiou AG, Chen SX, Cheng H, Fullard JF, Bendl J, Liu Y, Roussos P, Björkegren JL, Liu Y, Poon WW, Hao K, Marcora E, Goate AM. 2021. Integration of Alzheimer's disease genetics and myeloid genomics identifies disease risk regulatory elements and genes. *Nature Communications* 12:1–14. DOI: 10.1038/s41467-021-21823-y.

Pairo-Castineira E, Clohisey S, Klaric L, Bretherick AD, Rawlik K, Pasko D, Walker S, Parkinson N, Fourman MH, Russell CD, Furniss J, Richmond A, Gountouna E, Wrobel N, Harrison D, Wang B, Wu Y, Meynert A, Griffiths F, Oosthuyzen W, Kousathanas A, Moutsianas L, Yang Z, Zhai R, Zheng C, Grimes G, Beale R, Millar J, Shih B, Keating S, Zechner M, Haley C, Porteous DJ, Hayward C, Yang J, Knight J, Summers C, Shankar-Hari M, Klenerman P, Turtle L, Ho A, Moore SC, Hinds C, Horby P, Nichol A, Maslove D, Ling L, McAuley D, Montgomery H, Walsh T, Pereira AC, Renieri A, Shen X, Ponting CP, Fawkes A, Tenesa A, Caulfield M, Scott R, Rowan K, Murphy L, Openshaw PJM, Semple MG, Law A, Vitart V, Wilson JF, Baillie JK. 2021. Genetic mechanisms of critical illness in COVID-19. *Nature* 591:92–98. DOI: 10.1038/s41586-020-03065-y.

Palmer N, Kaldis P. 2020. Less-well known functions of cyclin/CDK complexes. *Seminars in Cell and Developmental Biology* 107:54–62. DOI: 10.1016/j.semcdb.2020.04.003.

Paradis E, Schliep K. 2019. Ape 5.0: An environment for modern phylogenetics and evolutionary analyses in R. *Bioinformatics* 35:526–528. DOI: 10.1093/bioinformatics/bty633.

Peng X, Xu E, Liang W, Pei X, Chen D, Zheng D, Zhang Y, Zheng C, Wang P, She S, Zhang Y, Ma J, Mo X, Zhang Y, Ma D, Wang Y. 2016. Identification of FAM3D as a newendogenous chemotaxis agonist for the formyl peptide receptors. *Journal of Cell Science* 129:1831–1842. DOI: 10.1242/jcs.183053.

Prokunina-Olsson L, Alphonse N, Dickenson RE, Durbin JE, Glenn JS, Hartmann R, Kotenko S V., Lazear HM, O'Brien TR, Odendall C, Onabajo OO, Piontkivska H, Santer DM, Reich NC, Wack A, Zanoni I. 2020. COVID-19 and emerging viral infections: The case for interferon lambda. *Journal of Experimental Medicine* 217. DOI: 10.1084/jem.20200653.

Rangel R, Sun Y, Guzman-Rojas L, Ozawa MG, Sun J, Giordano RJ, Van Pelt CS, Tinkey PT, Behringer RR, Sidman RL, Arap W, Pasqualini R. 2007. Impaired angiogenesis in aminopeptidase N-null mice. *Proceedings of the National Academy of Sciences of the United States of America* 104:4588–4593. DOI: 10.1073/pnas.0611653104.

Reed AAC, Loh NY, Terryn S, Lippiat JD, Partridge C, Galvanovskis J, Williams SE, Jouret F, Wu FTF, Courtoy PJ, Andrew Nesbit M, Rorsman P, Devuyst O, Ashcroft FM, Thakker R V. 2010. CLC-5 and KIF3B interact to facilitate CLC-5 plasma membrane expression, endocytosis, and microtubular transport: Relevance to pathophysiology of Dent's disease. *American Journal of Physiology - Renal Physiology* 298:F365–F380. DOI: 10.1152/ajprenal.00038.2009.

Ripon MAR, Bhowmik DR, Amin MT, Hossain MS. 2021. Role of arachidonic cascade in COVID-19 infection: A review. *Prostaglandins and Other Lipid Mediators* 154:106539. DOI: 10.1016/j.prostaglandins.2021.106539.

Samuel RM, Majd H, Richter MN, Ghazizadeh Z, Zekavat SM, Navickas A, Ramirez JT, Asgharian H, Simoneau CR, Bonser LR, Koh KD, Garcia-Knight M, Tassetto M, Sunshine S, Farahvashi S, Kalantari A, Liu W, Andino R, Zhao H, Natarajan P, Erle DJ, Ott M, Goodarzi H, Fattahi F. 2020. Androgen Signaling Regulates SARS-CoV-2 Receptor Levels and Is Associated with Severe COVID-19 Symptoms in Men. *Cell Stem Cell* 27:876-889.e12. DOI: 10.1016/j.stem.2020.11.009.

Sánchez-Martín P, Komatsu M. 2020. Heparan sulfate and clusterin: Cleaning squad for extracellular protein degradation. *Journal of Cell Biology* 219. DOI: 10.1083/JCB.202001159.

Schäfer MKE, Altevogt P. 2010. L1CAM malfunction in the nervous system and human carcinomas. *Cellular and Molecular Life Sciences* 67:2425–2437. DOI: 10.1007/s00018-010-0339-1.

Schulz A, Müller NV, Van De Lest NA, Eisenreich A, Schmidbauer M, Barysenka A, Purfürst B, Sporbert A, Lorenzen T, Meyer AM, Herlan L, Witten A, Rühle F, Zhou W, De Heer E, Scharpfenecker M, Panáková D, Stoll M, Kreutz R. 2019. Analysis of the genomic architecture of a complex trait locus in hypertensive rat models links Tmem63C to kidney damage. *eLife* 8. DOI: 10.7554/eLife.42068.

Severe Covid-19 GWAS Group. 2020. Genomewide Association Study of Severe Covid-19 with Respiratory Failure. *New England Journal of Medicine* 383:1522–1534. DOI: 10.1056/nejmoa2020283.

Seymen CM. 2021. The other side of COVID-19 pandemic: Effects on male fertility. *Journal of Medical Virology* 93:1396–1402. DOI: 10.1002/jmv.26667.

Shepardson KM, Larson K, Johns LL, Stanek K, Cho H, Wellham J, Henderson H, Rynda-Apple A. 2018. IFNAR2 is required for anti-influenza immunity and alters susceptibility to post-influenza bacterial superinfections. *Frontiers in Immunology* 9:2589. DOI: 10.3389/fimmu.2018.02589.

Shepherd CE, Affleck AJ, Bahar AY, Carew-Jones F, Halliday GM. 2020. Intracellular and secreted forms of clusterin are elevated early in Alzheimer's disease and associate with both Aβ and tau pathology. *Neurobiology of Aging* 89:129–131. DOI: 10.1016/j.neurobiolaging.2019.10.025.

Singh MK, Mobeen A, Chandra A, Joshi S, Ramachandran S. 2021. A meta-analysis of comorbidities in COVID-19: Which diseases increase the susceptibility of SARS-CoV-2 infection? *Computers in Biology and Medicine* 130:104219. DOI: 10.1016/j.compbiomed.2021.104219.

Srinivasan K, Friedman BA, Etxeberria A, Huntley MA, van der Brug MP, Foreman O, Paw JS, Modrusan Z, Beach TG, Serrano GE, Hansen D V. 2020. Alzheimer's Patient Microglia Exhibit Enhanced Aging and Unique Transcriptional Activation. *Cell Reports* 31:107843. DOI: 10.1016/j.celrep.2020.107843.

Stefely JA, Christensen BB, Gogakos T, Cone Sullivan JK, Montgomery GG, Barranco JP, Van Cott EM. 2020. Marked factor V activity elevation in severe COVID-19 is associated with venous thromboembolism. *American Journal of Hematology* 95:1522–1530. DOI: 10.1002/ajh.25979.

Stehling O, Vashisht AA, Mascarenhas J, Jonsson ZO, Sharma T, Netz DJA, Pierik AJ, Wohlschlegel JA, Lill R. 2012. MMS19 assembles iron-sulfur proteins required for DNA metabolism and genomic integrity. *Science* 337:195–199. DOI: 10.1126/science.1219723.

Stelzer G, Rosen N, Plaschkes I, Zimmerman S, Twik M, Fishilevich S, Iny Stein T, Nudel R, Lieder I, Mazor Y, Kaplan S, Dahary D, Warshawsky D, Guan-Golan Y, Kohn A, Rappaport N, Safran M, Lancet D. 2016. The GeneCards suite: From gene data mining to disease genome sequence analyses. *Current Protocols in Bioinformatics* 2016:1.30.1-1.30.33. DOI: 10.1002/cpbi.5.

Sutton BS, Crosslin DR, Shah SH, Nelson SC, Bassil A, Hale AB, Haynes C, Goldschmidt-Clermont PJ, Vance JM, Seo D, Kraus WE, Gregory SG, Hauser ER. 2008. Comprehensive genetic analysis of the platelet activating factor acetylhydrolase (PLA2G7) gene and cardiovascular disease in case-control and family datasets. *Human Molecular Genetics* 17:1318–1328. DOI: 10.1093/hmg/ddn020.

Takeda K, Kaisho T, Akira S. 2003. Toll-like receptors. *Annual Review of Immunology* 21:335–376. DOI: 10.1146/annurev.immunol.21.120601.141126.

Thomas C, Moraga I, Levin D, Krutzik PO, Podoplelova Y, Trejo A, Lee C, Yarden G, Vleck SE, Glenn JS, Nolan GP, Piehler J, Schreiber G, Garcia KC. 2011. Structural linkage between ligand discrimination and receptor activation by Type i interferons. *Cell* 146:621–632. DOI: 10.1016/j.cell.2011.06.048.

Thul PJ, Akesson L, Wiking M, Mahdessian D, Geladaki A, Ait Blal H, Alm T, Asplund A, Björk L, Breckels LM, Bäckström A, Danielsson F, Fagerberg L, Fall J, Gatto L, Gnann C, Hober S, Hjelmare M, Johansson F, Lee S, Lindskog C, Mulder J, Mulvey CM, Nilsson P, Oksvold P, Rockberg J, Schutten R, Schwenk JM, Sivertsson A, Sjöstedt E, Skogs M, Stadler C, Sullivan DP, Tegel H, Winsnes C, Zhang C, Zwahlen M, Mardinoglu A, Pontén F, Von Feilitzen K, Lilley KS, Uhlén M, Lundberg E. 2017. A subcellular map of the human proteome. *Science* 356. DOI: 10.1126/science.aal3321.

Tschopp J, Jenne DE, Hertig S, Preissner KT, Morgenstern H, Sapino - AP, French L. 1993. Human megakaryocytes express clusterin and package it without apolipoprotein A-1 into α-granules. *Blood* 82:118–125. DOI: 10.1182/blood.v82.1.118.bloodjournal821118.

Uhlén M, Fagerberg L, Hallström BM, Lindskog C, Oksvold P, Mardinoglu A, Sivertsson Å, Kampf C, Sjöstedt E, Asplund A, Olsson IM, Edlund K, Lundberg E, Navani S, Szigyarto CAK, Odeberg J, Djureinovic D, Takanen JO, Hober S, Alm T, Edqvist PH, Berling H, Tegel H, Mulder J, Rockberg J, Nilsson P, Schwenk JM, Hamsten M, Von Feilitzen K, Forsberg M, Persson L, Johansson F, Zwahlen M, Von Heijne G, Nielsen J, Pontén F. 2015. Tissue-based map of the human proteome. *Science* 347:1260419–1260419. DOI: 10.1126/science.1260419.

Vainio P, Gupta S, Ketola K, Mirtti T, Mpindi JP, Kohonen P, Fey V, Perälä M, Smit F, Verhaegh G, Schalken J, Alanen KA, Kallioniemi O, Iljin K. 2011. Arachidonic acid pathway members PLA2G7, HPGD, EPHX2, and CYP4F8 identified as putative novel therapeutic targets in prostate cancer. *American Journal of Pathology* 178:525–536. DOI: 10.1016/j.ajpath.2010.10.002.

Verma S, Saksena S, Sadri-Ardekani H. 2020. ACE2 receptor expression in testes: Implications in coronavirus disease 2019 pathogenesis. *Biology of Reproduction* 103:449–451. DOI: 10.1093/biolre/ioaa080.

Vietri M, Stenmark H. 2018. Orchestrating Nuclear Envelope Sealing during Mitosis. *Developmental Cell* 47:541–542. DOI: 10.1016/j.devcel.2018.11.020.

Wang Q, Hao Y, Mo X, Wang L, Lu X, Huang J, Cao J, Li H, Gu D. 2010. PLA2G7 gene polymorphisms and coronary heart disease risk: A meta-analysis. *Thrombosis Research* 126:498–503. DOI: 10.1016/j.thromres.2010.09.009.

Wang Z, Xu X. 2020. scRNA-seq Profiling of Human Testes Reveals the Presence of the ACE2 Receptor, A Target for SARS-CoV-2 Infection in Spermatogonia, Leydig and Sertoli Cells. *Cells* 9:920. DOI: 10.3390/cells9040920.

de Wit NJW, IJssennagger N, Oosterink E, Keshtkar S, Hooiveld GJEJ, Mensink RP, Hammer S, Smit JWA, Müller M, der Meer R van. 2012. Oit1/Fam3D, a gut-secreted protein displaying nutritional status-dependent regulation. *Journal of Nutritional Biochemistry* 23:1425–1433. DOI: 10.1016/j.jnutbio.2011.09.003.

Witte DP, Aronow BJ, Stauderman ML, Stuart WD, Clay MA, Gruppo RA, Jenkins SH, Harmony JAK. 1993. Platelet activation releases megakaryocyte-synthesized apolipoprotein J, a highly abundant protein in atheromatous lesions. *American Journal of Pathology* 143:763–773.

Yamada T, Nakao K, Itoh H, Morii N, Shiono S, Sakamoto M, Sugawara A, Saito Y, Mukoyama M, Arai H, Eigyo M, Matsushita A, Imura H. 1988. Inhibitory action of leumorphin on vasopressin secretion in conscious rats. *Endocrinology* 122:985–990. DOI: 10.1210/endo-122-3-985.

Yamamoto K, Ishiai M, Matsushita N, Arakawa H, Lamerdin JE, Buerstedde J-M, Tanimoto M, Harada M, Thompson LH, Takata M. 2003. Fanconi Anemia FANCG Protein in Mitigating Radiation- and Enzyme-Induced DNA Double-Strand Breaks by Homologous Recombination in Vertebrate Cells. *Molecular and Cellular Biology* 23:5421–5430. DOI: 10.1128/mcb.23.15.5421-5430.2003.

Yeager CL, Ashmun RA, Williams RK, Cardellichio CB, Shapiro LH, Look AT, Holmes K V. 1992. Human aminopeptidase N is a receptor for human coronavirus 229E. *Nature* 357:420–422. DOI: 10.1038/357420a0.

Zhang P, Jiang W, Luo N, Zhu W, Fan L. 2019. IQ motif containing D (IQCD), a new acrosomal protein involved in the acrosome reaction and fertilisation. *Reproduction, Fertility and Development*. DOI: 10.1071/RD18416.

Zhang Q, Liu Z, Moncada-Velez M, Chen J, Ogishi M, Bigio B, Yang R, Arias AA, Zhou Q, Han JE, Ugurbil AC, Zhang P, Rapaport F, Li J, Spaan AN, Boisson B, Boisson-Dupuis S, Bustamante J, Puel A, Ciancanelli MJ, Zhang SY, Béziat V, Jouanguy E, Abel L, Cobat A, Casanova JL, Bastard P, Korol C, Rosain J, Philippot Q, Chbihi M, Lorenzo L, Bizien L, Neehus AL, Kerner G, Seeleuthner Y, Manry J, Le Voyer T, Boisson B, Boisson-Dupuis S, Bustamante J, Puel A, Zhang SY, Béziat V, Jouanguy

E, Abel L, Cobat A, Casanova JL, Bastard P, Rosain J, Philippot Q, Chbihi M, Lorenzo L, Bizien L, Neehus AL, Kerner G, Seeleuthner Y, Manry J, Le Voyer T, Boisson B, Boisson-Dupuis S, Bustamante J, Puel A, Zhang SY, Béziat V, Jouanguy E, Abel L, Cobat A, Casanova JL, Le Pen J, Schneider WM, Razooky BS, Hoffmann HH, Michailidis E, Rice CM, Sabli IKD, Hodeib S, Sancho-Shimizu V, Bilguvar K, Ye J, Maniatis T, Bolze A, Arias AA, Arias AA, Zhang Y, Notarangelo LD, Su HC, Zhang Y, Notarangelo LD, Su HC, Onodi F, Korniotis S, Karpf L, Soumelis V, Bonnet-Madin L, Amara A, Dorgham K, Gorochov G, Smith N, Duffy D, Moens L, Meyts I, Meade P, García-Sastre A, Krammer F, Corneau A, Masson C, Schmitt Y, Schlüter A, Pujol A, Khan T, Marr N, Fellay J, Fellay J, Fellay J, Roussel L, Vinh DC, Shahrooei M, Shahrooei M, Alosaimi MF, Alsohime F, Hasanato R, Mansouri D, Mansouri D, Mansouri D, Al-Saud H, Almourfi F, Al-Mulla F, Al-Muhsen SZ, Al Turki S, Al Turki S, van de Beek D, Biondi A, Bettini LR, D'Angio M, Bonfanti P, Imberti L, Sottini A, Paghera S, Quiros-Roldan E, Rossi C, Oler AJ, Tompkins MF, Alba C, Dalgard CL, Vandernoot I, Smits G, Goffard JC, Migeotte I, Haerynck F, Soler-Palacin P, Martin-Nalda A, Colobran R, Morange PE, Keles S, Çölkesen F, Ozcelik T, Yasar KK, Senoglu S, Karabela ŞN, Rodríguez-Gallego C, Rodríguez-Gallego C, Novelli G, Hraiech S, Tandjaoui-Lambiotte Y, Tandjaoui-Lambiotte Y, Duval X, Laouénan C, Duval X, Laouénan C, Laouénan C, Snow AL, Dalgard CL, Milner JD, Mogensen TH, Mogensen TH, Marr N, Spaan AN, Bustamante J, Ciancanelli MJ, Meyts I, Maniatis T, Soumelis V, Nussenzweig M, Nussenzweig M, Casanova JL, García-Sastre A, García-Sastre A, García-Sastre A, Lifton RP, Lifton RP, Lifton RP, Casanova JL, Foti G, Bellani G, Citerio G, Contro E, Pesci A, Valsecchi MG, Cazzaniga M, Abad J, Blanco I, Rodrigo C, Aguilera-Albesa S, Akcan OM, Darazam IA, Aldave JC, Ramos MA, Nadji SA, Alkan G, Allardet-Servent J, Allende LM, Alsina L, Alyanakian MA, Amador-Borrero B, Mouly S, Sene D, Amoura Z, Mathian A, Antolí A, Blanch GR, Riera JS, Moreno XS, Arslan S, Assant S, Auguet T, Azot A, Bajolle F, Bustamante J, Lévy R, Oualha M, Baldolli A, Ballester M, Feldman HB, Barrou B, Beurton A, Bilbao A, Blanchard-Rohner G, Blandinières A, Rivet N, Blazquez-Gamero D, Bloomfield M, Bolivar-Prados M, Clavé P, Borie R, Bosteels C, Lambrecht BN, van Braeckel E, Bousfiha AA, Bouvattier C, Vincent A, Boyarchuk O, Bueno MRP, Castro M V., Matos LRB, Zatz M, Agra JJC, Calimli S, Capra R, Carrabba M, Fabio G, Casasnovas C, Vélez-Santamaria V, Caseris M, Falck A, Poncelet G, Castelle M, Castelli F, de Vera MC, Catherinot E, Chalumeau M, Toubiana J, Charbit B, Li Z, Pellegrini S, Cheng MP, Clotet B, Codina A, Colkesen F, Çölkesen F, Colobran R, Comarmond C, Dalmau D, Dalmau D, Darley DR, Dauby N, Dauger S, Le Bourgeois F, Levy M, de Pontual L, Dehban A, Delplancq G, Demoule A, Diehl JL, Dobbelaere S, Durand S, Mircher C, Rebillat AS, Vilaire ME, Eldars W, Elgamal M, Elnagdy MH, Emiroglu M, Erdeniz EH, Aytekin SE, Euvrard R, Evcen R, Faivre L, Fartoukh M, Philippot Q, Faure M, Arquero MF, Flores C, Flores C, Flores C, Flores C, Francois B, Fumadó V, Fumadó V, Fumadó V, Fusco F, Ursini MV, Solis BG, de Diego RP, van Den Rym AM, Gaussem P, Gil-Herrera J, Gilardin L, Alarcon MG, Girona-Alarcón M, Goffard JC, Gok F, Yosunkaya A, González-Montelongo R, Íñigo-Campos A, Lorenzo-Salazar JM, Muñoz-Barrera A, Guerder A, Gul Y, Guner SN, Gut M, Hadjadj J, Haerynck F, Halwani R, Hammarström L, Hatipoglu N, Hernandez-Brito E, Heijmans C, Holanda-Peña MS, Horcajada JP, Hoste L, Hoste E, Hraiech S, Humbert L, Mordacq C, Thumerelle C, Vuotto F, Iglesias AD, Jamme M, Arranz MJ, Jordan I, Jorens P, Kanat F, Kapakli H, Kara I, Karbuz A, Yasar KK, Senoglu S, Keles S, Demirkol YK, Klocperk A, Król ZJ, Kuentz P, Kwan YWM, Lagier JC, Lau YL, Leung D, Leo YS, Young BE, Lopez RL, Levin M, Linglart A, Loeys B, Louapre C, Lubetzki C, Luyt CE, Lye DC, Mansouri D, Marjani M, Pereira JM, Martin A, Soler-Palacín P, Pueyo DM, Martinez-Picado J, Marzana I, Matthews G V., Mayaux J, Parizot C, Quentric P, Mège JL, Raoult D, Melki I, Meritet JF, Metin O, Meyts I, Mezidi M, Migeotte I, Taccone F, Millereux M, Mirault T, Mirsaeidi M, Melián AM, Martinez AM, Morange P, Morelle G, Naesens L, Nafati C, Neves JF, Ng LFP, Medina YN, Cuadros EN, Gonzalo Ocejo-Vinyals J, Orbak Z, Özçelik T, Pan-Hammarström Q, Pascreau T, Paz-Artal E, Philippe A, Planas-Serra L, Schluter A, Ploin D, Viel S, Poissy J, Pouletty M, Reisli I, Ricart P, Richard JC, Rivière JG, Rodriguez-Gallego C, Rodriguez-Gallego C, Rodríguez-Palmero A, Romero CS, Rothenbuhler A, Rozenberg F, del Prado MYR, Sanchez O, Sánchez-Ramón S, Schmidt M, Schweitzer CE, Scolari F, Sediva A, Seijo LM, Seppänen MRJ, Ilovich AS, Slabbynck H, Smadja DM, Sobh A, Solé-Violán J, Soler C, Stepanovskiy Y, Stoclin A, Tandjaoui-Lambiotte Y, Taupin JL, Tavernier SJ, Terrier B, Tomasoni G, Alvarez JT, Trouillet-Assant S, Troya J, Tucci A, Uzunhan Y, Vabres P, Valencia-Ramos J, van de Velde S, van Praet J, Vandernoot I, Vatansev H, Vilain C, Voiriot G, Yucel F, Zannad F, Belot A, Bole-Feysot C, Lyonnet S, Masson C, Nitschke P, Pouliet A, Schmitt Y, Tores F, Zarhrate M, Shahrooei M, Abel L, Andrejak C, Angoulvant F, Bachelet D, Bhavsar K, Bouadma L, Chair A, Couffignal C, Silveira C Da, Debray MP, Duval X, Eloy P, Esposito-Farese M, Ettalhaoui N, Gault N, Ghosn J, Gorenne I, Hoffmann I, Kafif O, Kali S, Khalil A, Laouénan C, Laribi S, Le M, Le Hingrat Q, Lescure FX, Lucet JC, Mentré F, Mullaert J, Peiffer-Smadja N, Peytavin G, Roy C, Schneider M, Mohammed NS, Tagherset L, Tardivon C, Tellier MC, Timsit JF, Trioux T, Tubiana S, Basmaci R, Behillil S, Beluze M, Benkerrou D, Dorival C, Meziane A, Téoulé F, Bompart F, Bouscambert M, Gaymard A, Lina B, Rosa-Calatrava M, Terrier O, Caralp M, Cervantes-Gonzalez M, D'Ortenzio E, Puéchal O, Semaille C, Coelho A, Diouf A, Hoctin A, Mambert M, Couffin-Cadiergues S, Deplanque D, Descamps D, Visseaux B, Desvallées M, Khan C, Diallo A, Mercier N, Paul C, Petrov-Sanchez V, Dubos F, Enouf VVE, Mouquet H, Esperou H, Jaafoura S, Papadopoulos A, Etienne M, Gigante T, Rossignol B, Guedj J, Le Nagard H, Lingas G, Neant N, Kaguelidou F, Lévy Y, Wiedemann A, Lévy Y, Wiedemann A, Levy-Marchal C, Malvy D, Noret M, Pages J, Picone O, Rossignol P, Tual C, Veislinger A, van der Werf S, Vanel N, Yazdanpanah Y, Alavoine L, Costa Y, Duval X, Ecobichon JL, Frezouls W, Ilic-Habensus E, Leclercq A, Lehacaut J, Letrou S, Mandic M, Nouroudine M, Quintin C, Rexach J, Tubiana S, Vignali V, Amat KKA, Behillil S, Enouf V, van der Werf S, Bielicki J, Bruijning P, Burdet C, Burdet C, Caumes E, Charpentier C, Damond F, Descamps D, Le Hingrat Q, Visseaux B, Coignard B, Couffin-Cadiergues S, Delmas C, Espérou H, Roufai L, Dechanet A, Houhou N, Kafif O, Kikoine J, Manchon P, Piquard V, Postolache A, Terzian Z, Lebeaux D, Lina B, Lucet JC, Malvy D, Meghadecha M, Motiejunaite J, Thy M, van Agtmael M, Bomers M, Chouchane O, Geerlings S, Goorhuis B, Grobusch MP, Harris V, Hermans SM, Hovius JW, Nellen J, Peters E, van der Poll T, Prins JM, Reijnders T, Schinkel M, Sigaloff K, Stijnis CS, van der Valk M, van Vugt M, Joost Wiersinga W, Algera AG, van Baarle F, Bos L, Botta M, de Bruin S, Bulle E, Elbers P, Fleuren L, Girbes A, Hagens L, Heunks L, Horn J, van Mourik N, Paulus F, Raasveld J, Schultz MJ, Smit M, Stilma W, Thoral P, Tsonas A, de Vries H, Bax D, Cloherty A, Beudel M, Brouwer MC, Koning R, van de Beek D, Bogaard HJ, de Brabander J, de Bree G, Bugiani M, Geerts B, Hollmann MW, Preckel B, Veelo D, Geijtenbeek T, Hafkamp F, Hamann J, Hemke R, de Jong MD, Schuurman A, Teunissen C, Vlaar APJ, Wouters D, Zwinderman AH, Abel L, Aiuti A, Muhsen S Al, Al-Mulla F, Anderson MS, Arias AA, Feldman HB, Bogunovic D, Itan Y, Bolze A, Cirulli E, Barrett KS, Washington N, Bondarenko A, Bousfiha AA, Brodin P, Bryceson Y, Bustamante CD, Butte M, Casari G, Chakravorty S, Christodoulou J, Le Mestre S, Condino-Neto A, Cooper MA, Dalgard CL, David A, DeRisi JL, DeRisi JL, Desai M, Drolet BA, Espinosa S, Fellay J, Flores C, Franco JL, Gregersen PK, Haerynck F, Hagin D, Halwani R, Heath J,

Henrickson SE, Hsieh E, Imai K, Karamitros T, Kisand K, Ku CL, Lau YL, Ling Y, Lucas CL, Maniatis T, Mansouri D, Marodi L, Meyts I, Milner J, Mironska K, Mogensen T, Morio T, Ng LFP, Notarangelo LD, Su HC, Novelli A, Novelli G, O'Farrelly C, Okada S, Ozcelik T, de Diego RP, Planas AM, Prando C, Pujol A, Quintana-Murci L, Renia L, Renieri A, Rodríguez-Gallego C, Sancho-Shimizu V, Sankaran V, Shahrooei M, Snow A, Soler-Palacín P, Spaan AN, Tangye S, Turvey S, Uddin F, Uddin MJ, Uddin MJ, van de Beek D, Vazquez SE, Vinh DC, von Bernuth H, Zawadzki P, Casanova JL, Jing H, Tung W, Meguro K, Shaw E, Jing H, Tung W, Shafer S, Zheng L, Zhang Z, Kubo S, Chauvin SD, Meguro K, Shaw E, Lenardo M, Luthers CR, Bauman BM, Shafer S, Zheng L, Zhang Z, Kubo S, Chauvin SD, Lenardo M, Lack J, Karlins E, Hupalo DM, Rosenberger J, Sukumar G, Wilkerson MD, Zhang X. 2020. Inborn errors of type I IFN immunity in patients with life-threatening COVID-19. *Science* 370. DOI: 10.1126/science.abd4570.

Zhao H, Lai X, Xu X, Sui K, Bu X, Ma W, Li D, Guo K, Xu J, Yao L, Li W, Su J. 2015. Histochemical analysis of testis specific gene 13 in human normal and malignant tissues. *Cell and Tissue Research* 362:653–663. DOI: 10.1007/s00441-015-2227-3.

Zhao X, Yan X, Liu Y, Zhang P, Ni X. 2016. Co-expression of mouse TMEM63A, TMEM63B and TMEM63C confers hyperosmolarity activated ion currents in HEK293 cells. *Cell Biochemistry and Function* 34:238–241. DOI: 10.1002/cbf.3185.
